# Variation in relaxation of non-photochemical quenching between the founder genotypes of the soybean (Glycine max) nested association mapping population

**DOI:** 10.1101/2024.06.03.597170

**Authors:** Dhananjay Gotarkar, Anthony Digrado, Yu Wang, Lynn Doran, Bethany Blakey, Brian W. Diers, Daniel J. Eck, Steven J. Burgess

## Abstract

Improving the efficiency of crop photosynthesis has the potential to increase yields. Genetic manipulation showed photosynthesis can be improved by speeding up relaxation of photoprotective mechanisms during sun to shade transitions. However, it is unclear if natural variation in relaxation of non-photochemical quenching (NPQ) can be exploited in crop breeding programs. To address this issue, we measured six NPQ parameters in the 40 founder lines and common parent of a Soybean Nested Association Mapping (SoyNAM) panel over two field seasons in Illinois. NPQ parameters did not show consistently variable trends throughout development, and variation between sampling days suggests environmental impacts on NPQ which last more than 24 hours. 17 genotypes were found to show small but consistent differences in NPQ relaxation kinetics relative to a reference line providing a basis for future mapping studies. Finally, a soybean canopy model predicted available phenotypic variation could result in a 1.6% difference in carbon assimilation when comparing fastest and slowest relaxing NPQ values.

**Significance Statement:** Evidence suggests increasing the rate of relaxation of photoprotection can lead to improved biomass and yield. We compare photoprotection relaxation rates in 41 diverse soybean genotypes grown in the field, identifying lines with faster rates of relaxation, and predict a potential 1.6% difference in daily carbon assimilation which could contribute to improving soybean performance.

## Introduction

Leaves within a canopy are exposed to sunflecks and shadeflecks, caused by intermittent cloud cover, wind induced leaf movements, and the changing angle of the sun (Kaiser *et al*., 2018). Balancing variable energy supply with demand for reducing equivalents is essential for efficient photosynthesis (Kramer and Evans, 2011) and a delay in adjustment of biochemical or gas diffusional processes can lead to reduced carbon assimilation (Sakoda *et al*., 2022).

In response to excess light, plants activate photoprotective mechanisms (Demmig-Adams *et al*., 2012; Jahns and Holzwarth, 2012) which deal with production of reactive oxygen species and limit photoinhibition (Pinnola and Bassi, 2018). Excess absorbed light energy can be dissipated by non-photochemical quenching (NPQ) of chlorophyll excited states, reducing the likelihood that damaging reactive oxygen species are formed (Patricia Müller *et al*., 2001; Ruban and Wilson, 2021). Transgenic approaches have shown increasing NPQ in rice can lead to increased biomass production in glasshouse conditions by alleviating photoinhibition (Hubbart *et al*., 2018). However, excess NPQ, or delays in relaxing NPQ during shadeflecks, are predicted to cause unnecessary dissipation of energy, reducing the efficiency of photosynthesis (Zhu *et al*., 2004; Burgess *et al*., 2019). As a result, speeding up NPQ activation and relaxation can improve photosynthetic efficiency (Kromdijk et al., 2016; Garcia-Molina and Leister, 2020; De Souza et al., 2022; Lehretz et al., 2022). Support for a link between fast relaxation of NPQ and increased biomass accumulation comes from African rice genotypes grown under controlled conditions (Cowling *et al*., 2022). Further, analysis of transgenic plants in small-scale field experiments suggested an increase biomass in Tobacco (Kromdijk *et al*., 2016) and seed production in soybean (De Souza *et al*., 2022) could be achieved if NPQ relaxation is accelerated. However, there have been contrasting results in Arabidopsis (Garcia-Molina and Leister, 2020) and potato (Lehretz *et al*., 2022), while the manipulating NPQ on soybean seed production differed between years (De Souza *et al*., 2022). Taken together, these data point to the need to further understand species the relation of photoprotection to whole plant physiology if the potential benefits of altering NPQ are to be translated to commercial crop varieties (Kaiser *et al*., 2019; Leister, 2023).

Most experiments investigating NPQ have been performed under controlled conditions or at a single time point during development. However, NPQ is highly dynamic and sensitive to any perturbation which impacts carbon assimilation or cellular redox state. This means in a field environment, NPQ is likely to vary in response to weather conditions (Zhu *et al*., 2009; Porcar-Castell, 2011; Sun *et al*., 2020), but the relationship between genotype x environment interactions on NPQ kinetics is poorly understood. Two studies with field grown soybean have assessed photosynthetic parameters with analysis of chlorophyll fluorescence, using canopy reflectance to calculate photochemical reflectance index (PRI) as a proxy for NPQ (Herritt *et al*., 2016), and OJIP transients to look at variation in fluorescence kinetics (Herritt *et al*., 2018). However, the individual components of NPQ relaxation were not assessed. Therefore, the extent to which individual components of NPQ co-vary in natural populations and whether they can be selected in breeding programs remains unclear.

A nested association mapping panel has been developed for soybean with the aim of identifying beneficial alleles possessed by elite and exotic germplasm (SoyNAM)(Song *et al*., 2017; Diers *et al*., 2018). This population has proved useful for studying agronomic traits and how they impact yield (Song *et al*., 2017; Diers *et al*., 2018; Lopez *et al*., 2019; Montes *et al*., 2022). We sought to use this resource for analysis of NPQ relaxation, and kinetics were measured for the 41 SoyNAM founder lines over the course of a field season in 2021 and 2022. The goals of this study were to (1) assess if NPQ kinetics vary in response to developmental and field environmental conditions, (2) identify soybean genotypes with fast relaxation kinetics that could serve as the basis for genetic mapping, and (3) test the impact of altering NPQ relaxation on carbon assimilation using a soybean canopy model given existing diversity.

Six parameters were calculated by fitting a double exponential function to the decay in NPQ on transition to low light (Dall’Osto *et al*., 2014), including fast (qE) (Krause *et al*., 1982) and intermediate (qM) relaxing NPQ and their respective rate constants (r_qE_ and r_qM_), in addition to long term NPQ (qI) and maximum NPQ reached during high light treatment (Table 1). Repeat measurements were taken throughout the growing season and a mixed effects linear modeling approach was used to identify lines with significantly different values for NPQ relaxation. Finally, a canopy photosynthesis model (Wang *et al*., 2020) was used to estimate the potential impact of genetic improvement in NPQ relaxation on soybean photosynthesis.

**Table 1.**
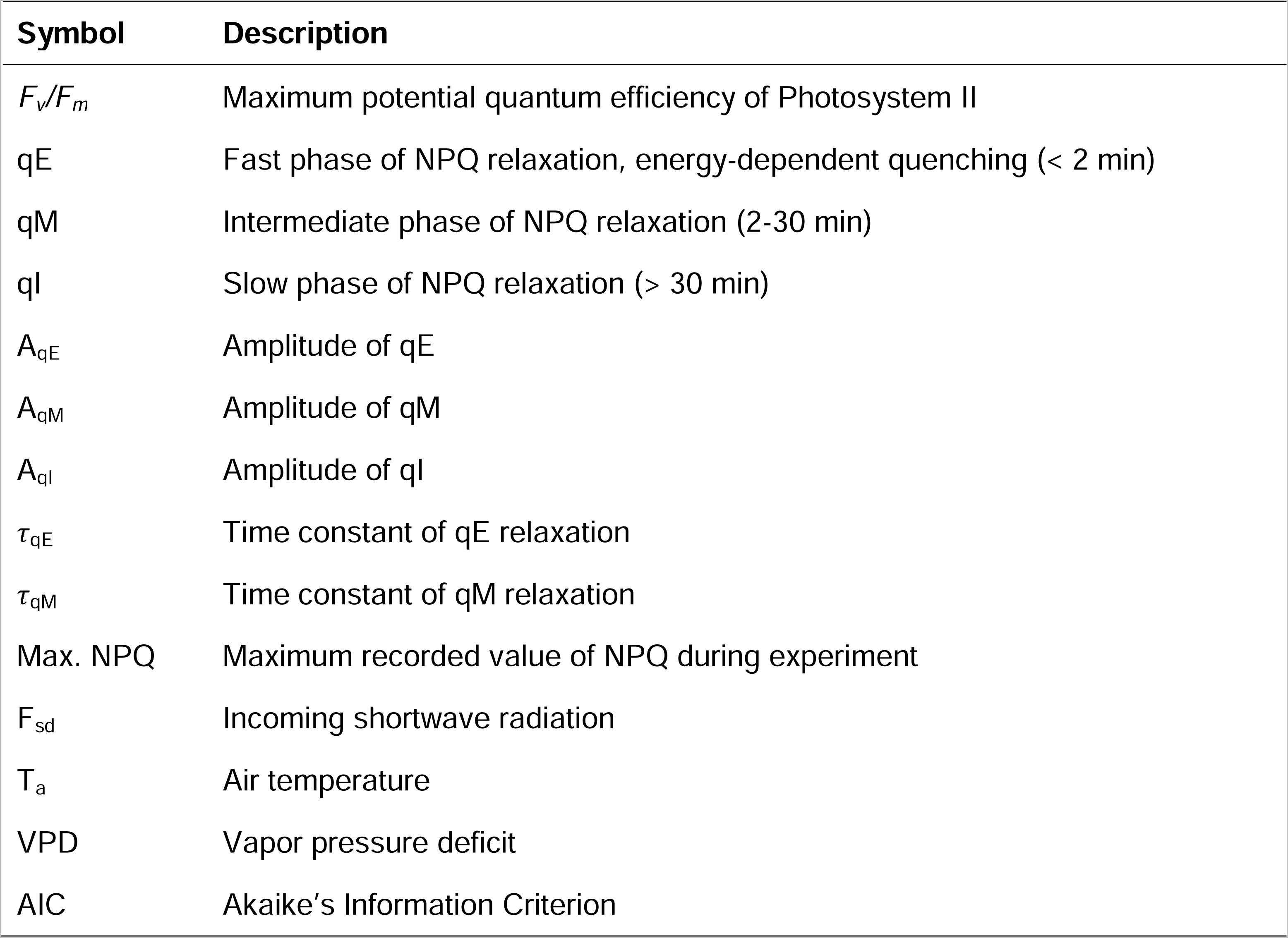
List of Abbreviations.

## Results

Measurements of NPQ relaxation in field grown plants were taken between V1 to R6 maturity stages in 2021 and 2022 (Figure 1, Table S1-2). Although NPQ parameters remained largely consistent over the course of the growing season, some days exhibited variations in mean values and variance. For instance, declines in r_qE_ and A_qM_ were observed on the fourth and fifth sampling days in 2022 (Figure 1a and b).

**Figure 1:**
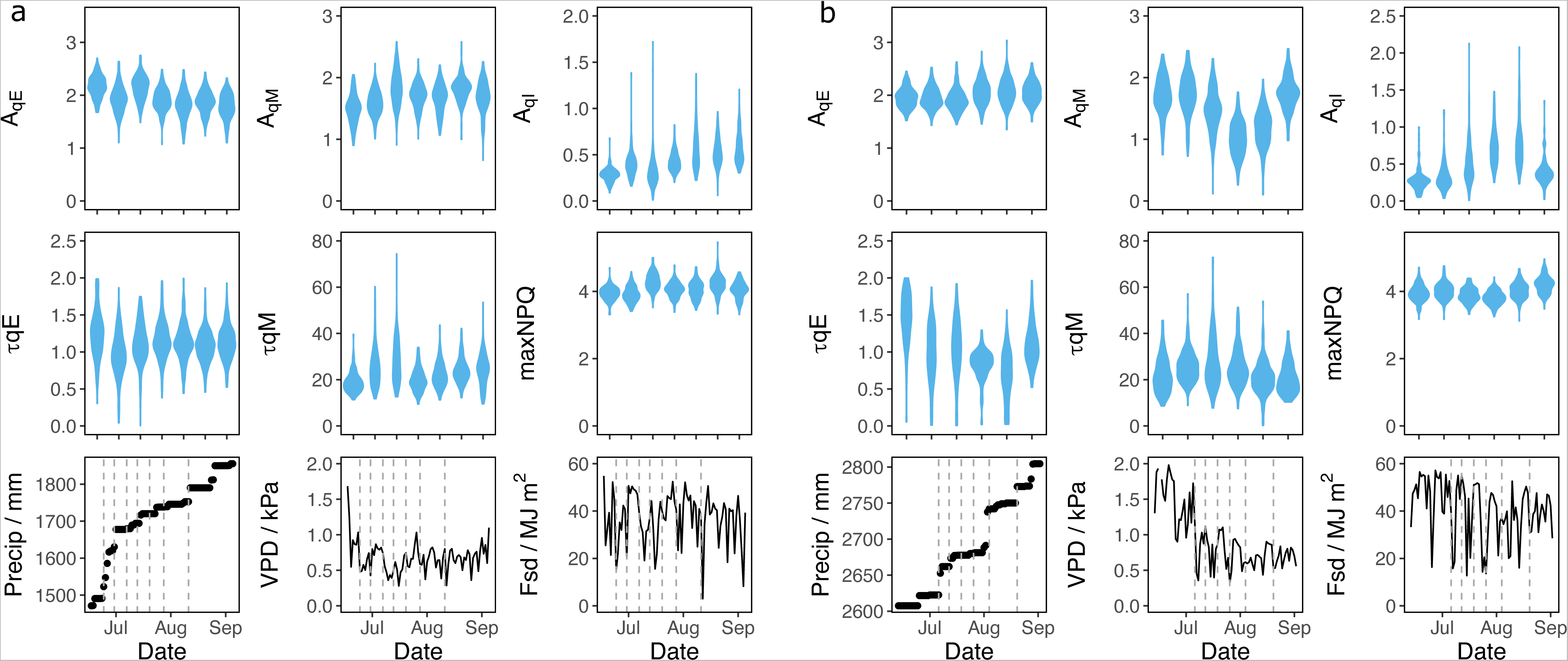
Comparison of NPQ relaxation kinetics and weather conditions over the course of two field seasons, showing field level summaries of (a) 2021 and (b) 2022. Violin plots are combined plot-level averages for all SoyNAM genotypes on a given sampling day. Sampling dates are indicated by grey dotted lines. Precipitation is represented as cumulative values from the beginning of the year, whereas VPD and Fsd are mean daily values.

### Variation in NPQ relaxation parameters

For all parameters measured there were significant differences between genotypes (G) and days (E) (Table 2). This indicated significant differences between genotypes and a strong impact of the environment on NPQ relaxation. However, there was no significant G x E interaction, suggesting genotypes responded similarly to the environment (Table 2). A stepwise AIC-based model revealed that the environment explained 10-47% of the observed variance in NPQ relaxation depending on the parameter and year (Figure 2). The parameter Aq_I_ was the best explained by the environment, with a R^2^>0.3 in both years. The parameter r_qM_ remained the parameter for which the environment had the least predictive power, with a R^2^<0.16 for both years. The R^2^ for the other parameters varied between years with, for instance, r_qE_ showing a R^2^ of 0.08 in 2021 and 0.32 in 2022. This could be explained by the range of variation observed in these parameters during those years. For both 2021 and 2022, the high values for the Precip and F_sd_ coefficients indicate their strong impact on the parameters, followed by VPD. The years 2021 and 2022 distinguished themselves as VPD_7day had a stronger impact in 2021 compared to 2022 where VPD was preferred by the model. The variable Ta_7day had a significant impact on the NPQ parameters in 2022. Between years, the parameters were affected differently by their environment, with A_qI_ being affected positively by Ta_7day, Precip, and F_sd_ in 2021, but showing the opposite behavior in 2022. This might be due different environmental patterns between the two years, with 2022 showing higher VPD and lower precipitation at the beginning of the growing season compared to 2021. However, A_qE_, r_qE_, and maxNPQ were affected in the same manner by the environment. When the two years were combined in a single model, Precip_7day, Precip_cum, Fsd_7day, and cumFn were included in the final models (Figure S1). Their absence in the 2021 and 2022 models suggest these variables mostly contributed to explain the year to year variation in NPQ relaxation parameters.

**Figure 2.**
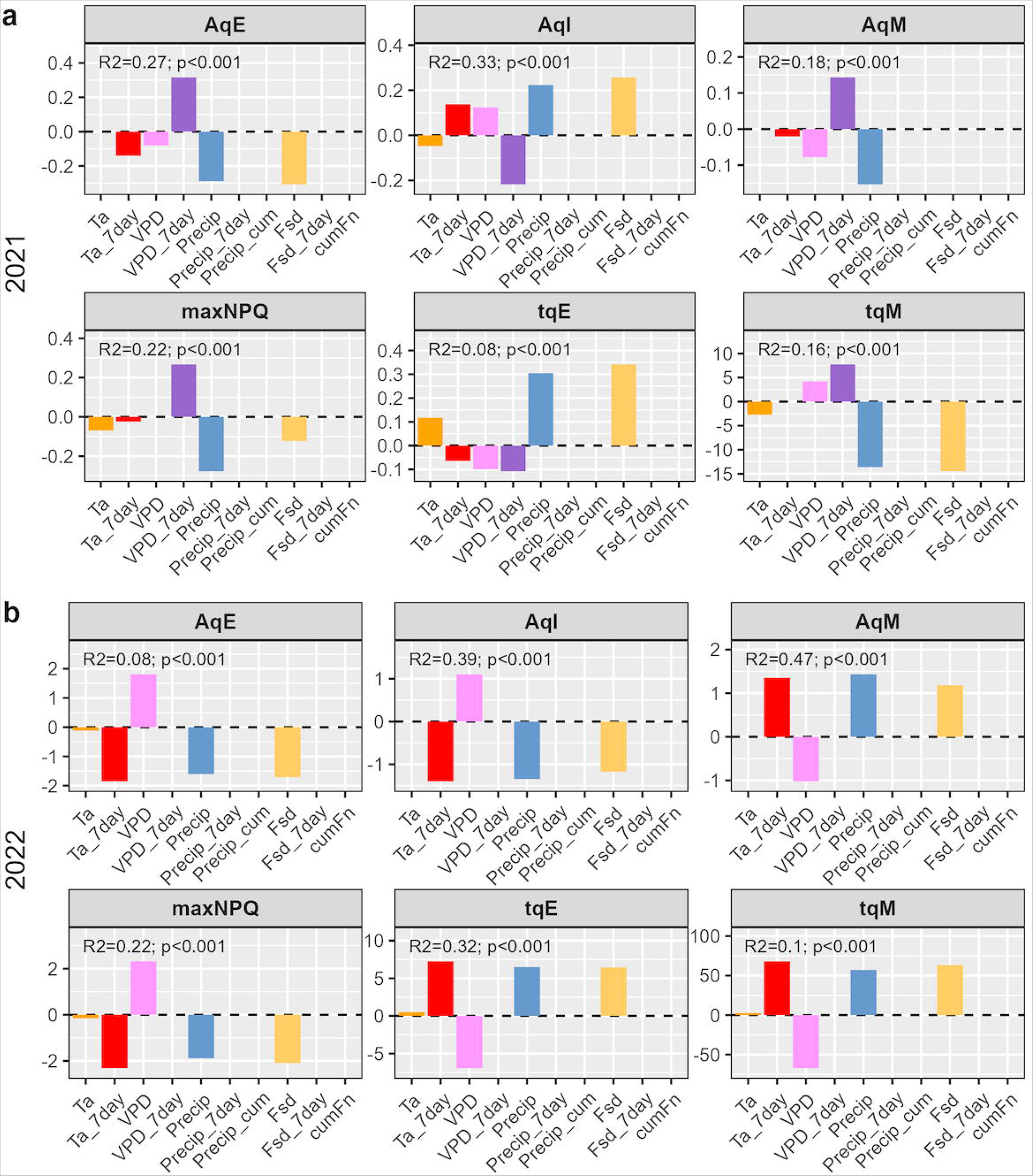
Coefficients for the best minimum adequate model (lowest AIC) for each NPQ relaxation parameters in 2021 and 2022. The R^2^ and p-value are shown for each model.

**Table 2.** Table of F-test P-values obtained by fitting an ANOVA model for each NPQ relaxation parameters with the genotypes (G) and days (or environment, E) set as fixed factors (i.e., variable ∼ Day + Genotype + Day * Genotype).

| Variable | G | E | GxE |
| --- | --- | --- | --- |
| $A_{qI}$ | <0.001 | <0.001 | 0.8325 |
| $A_{qE}$ | <0.001 | <0.001 | 0.9754 |
| $\tau_{qE}$ | <0.001 | <0.001 | 0.1193 |
| $A_{qM}$ | <0.001 | <0.001 | 0.9613 |
| $\tau_{qM}$ | <0.001 | <0.001 | 0.9463 |
| Max. NPQ | <0.001 | <0.001 | 0.8849 |

Canonical correlation analysis (CCA) was used to further explore the relationship between NPQ relaxation parameters and the environment within the NAM founders. The canonical correlations between pairs were ≥0.4 (p-value<0.001, Table 3), allowing association of changes in the environment with changes in NPQ relaxation. For both years, CCA revealed seasonal patterns in the NPQ relaxation parameters (Figure 3). In 2021, the CCA showed a change in environmental conditions throughout the season, with an increase in VPD, Ta, F_sd_, cumFn, and a decrease in precipitation between June 24^th^ and July 20^th^ (Figure 3b, d). This was associated with an increase in A_qI_ and a decline in A_qE_ (Figure 3a, c). The NPQ parameter r_qE_ remained constant. This is in accordance with the stepwise regression analysis which showed a low predictive power of environmental variable on r_qE_ in 2021 (Figure 2). Past July 20th, the environmental conditions showed less variation. This was associated with less fluctuation in the different NPQ parameters. July 7^th^ distinguished itself from the other days by an increase in A_qM_, maxNPQ, and r_qM_. In 2022, the CCA showed a somewhat similar pattern with an increase in A_qI_ at the beginning of the growing season (from July 6th to July 26th) but accompanied by a decline in A_qM_ (Figure 3e, g). This was associated with lower Ta, VPD, and F_sd_ over the same period (Figure 3f, h). The transition from August to July was marked by increasing precipitation and a decline in Ta_7day. This was associated with increasing maxNPQ and a decline in r_qM_. Between August 4th and August 20th, the opposite behavior was observed with an increase in A_qM_ and a decline in A_qI_. Overall, both years showed an increase in A_qI_ at the beginning of the season and highlighted a seasonal behavior with a period of important changes in NPQ relaxation associated with changes in environmental conditions. CCA also revealed different behavior in some NPQ relaxation parameters between years, with r_qE_ showing no variations in response to its environment in 2021 but being positively associated with Ta_7day, VPD, and cumFn in 2022. The CCA revealed no striking differences between the different NAM groups as they showed the same seasonal behavior in both years, which suggest a stronger impact of the environment on NPQ relaxation than a genotypic variation over the season.

**Figure 3.**
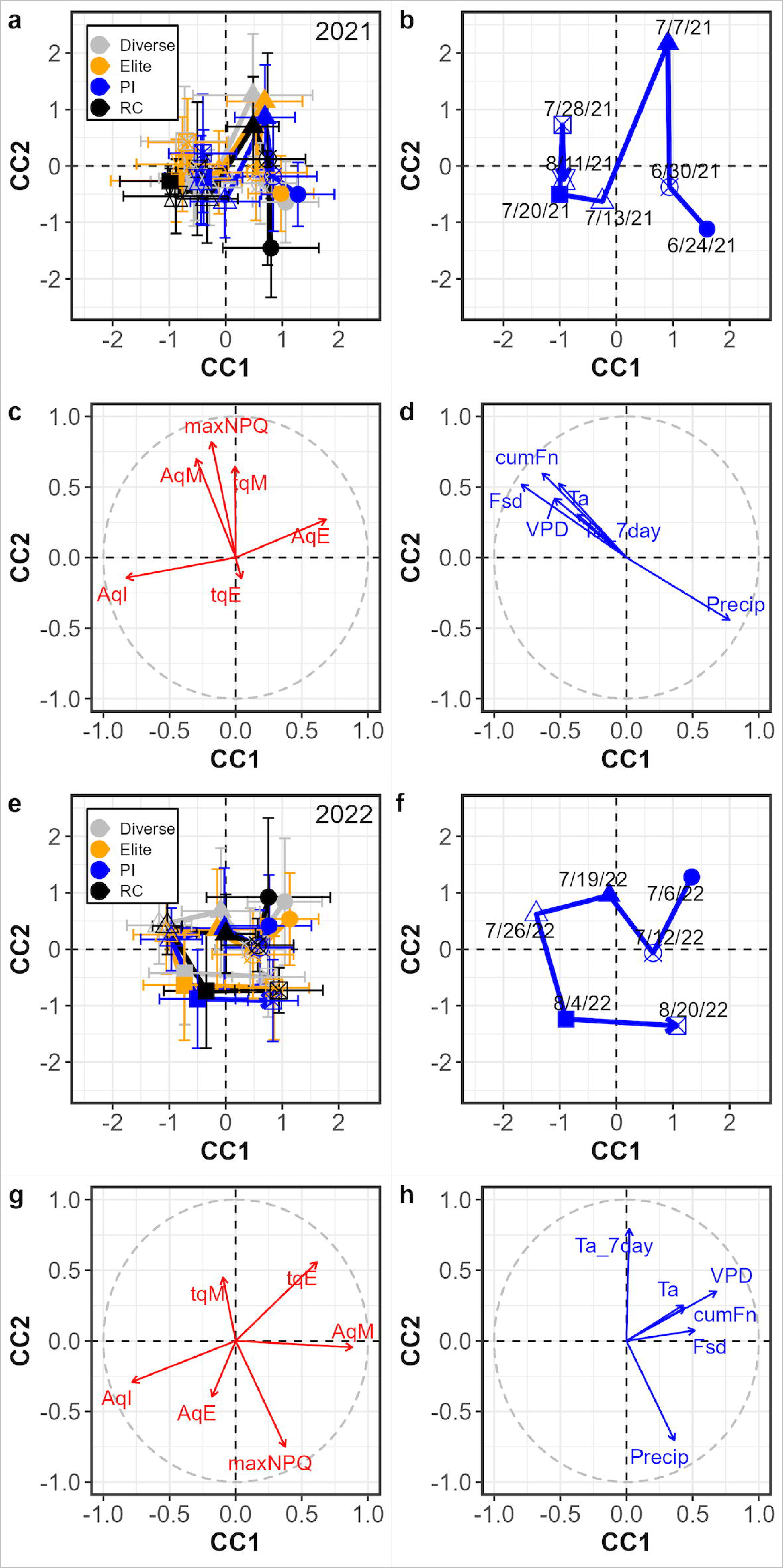
Canonical correlation analysis (CCA) displaying the relationships between the NPQ relaxation parameters and the environmental variables in (a-d) 2021 and (e-h) 2022. (a-b, e-f) CCA showing the spatial distribution of the different observations on the canonical axis (CC). The average (+-standard deviation) for the different NAM groups (with diverse, elite, PI, and RC groups in grey, orange, blue and black, respectively) at different days is represented by different shapes. For each group, a solid line connects those points to represent their evolution throughout the season. (c-d, g-h) The correlation circle showing the relationships between variables. Variables related to the NPQ relaxation parameters, and the environments are represented in red and blue, respectively. The canonical correlation and associated statistics are shown in table 4.

**Table 3.** Canonical correlations between pairs and associated Wilks lambda tests for the years 2021 and 2022.

|  | Canonical pairs for 2021 |  |  | Canonical pairs for 2022 |  |  |
| --- | --- | --- | --- | --- | --- | --- |
|  | CC1 | CC2 | CC3 | CC1 | CC2 | CC3 |
| Can. Corr. | 0.667 | 0.513 | 0.397 | 0.76 | 0.491 | 0.393 |
| Wilks | 0.331 | 0.597 | 0.81 | 0.258 | 0.61 | 0.804 |
| F | 44.83 | 28.458 | 17.562 | 44.724 | 21.473 | 14.364 |
| df | 36 | 25 | 16 | 36 | 25 | 16 |
| p-value | <0.001 | <0.001 | <0.001 | <0.001 | <0.001 | <0.001 |

**Table 4.** Table of F-test P-values obtained from a one-way ANOVA on the principal component (PC) coordinates with the NAM population groups set as a fixed factor (i.e., PC ∼ Groups).

| Day | PC1 | PC2 | PC3 |
| --- | --- | --- | --- |
| 6/24/2021 | 0.1857 | 0.0378 | 0.8604 |
| 6/30/2021 | 0.577 | 0.9843 | 0.2048 |
| 7/7/2021 | 0.8924 | 0.9776 | 0.7902 |
| 7/13/2021 | 0.1689 | 0.3434 | 0.0026 |
| 7/20/2021 | 0.1532 | 0.3742 | 0.6809 |
| 7/28/2021 | 0.6763 | 0.1174 | 0.5885 |
| 8/11/2021 | 0.1156 | 0.3003 | 0.3555 |
| 7/6/2022 | 0.1556 | 0.2845 | 0.0379 |
| 7/12/2022 | 0.132 | 0.9386 | 0.5951 |
| 7/19/2022 | 0.4572 | 0.1662 | 0.5331 |
| 7/26/2022 | 0.4417 | 0.3223 | 0.3501 |
| 8/4/2022 | 0.9584 | 0.0289 | 0.5888 |
| 8/20/2022 | 0.049 | 0.7222 | 0.2524 |

### Variation in NPQ parameters between NAM founders

To identify any variation between the NAM founders within a day, a Principal Components Analysis (PCA) was run for each day (Figure S2-14). The Diverse, Elite, PI, and RC groups are highlighted. The results showed no distinction between those groups, which tended to overlap (Figure S2-14). To determine if the groups were significantly distinct from each other, a linear model followed by an ANOVA was used on the principal component coordinate with the ‘NAM groups’ set as a fixed factor. This analysis showed some significant differences between groups on some dates (Table 4). Groups clusters also separated between group on 6/24/21 and 8/4/22 on the PC2. Groups also separated on the PC3 on 7/13/21 (Figure S5) and 7/6/22 (Figure S9). This suggest that some groups distinguished themselves from the others on the parameters that were the most strongly correlated with these axes. For instance, on 8/4/22 (Figure S13), groups separated on the second component, with PIs showing higher values in A_qE_ and maximum NPQ on average (Figure S13). Founders of NAM10, NAM31, and NAM33 detached themselves from the population cluster on two or more occasions. For instance, on those occasions, NAM33 founder tended to locate in region of the PCA associated with higher maximum NPQ on 6/24/21 (Figure S2) and 7/28/21 (Figure S7).

When considering all days combined the largest range in values between genotypes was seen in mean r_qM_ values, which varied ∼1.5 fold between the founder genotypes of NAM5 (19.94) and NAM17 (29.3) (Figure 4; Table 5; Table S3). A small range of values was observed for all other parameters (Table 2; Figure 4) and no observable difference was seen between elite, diverse or PI genotypes (Figure 4) consistent with the PCA (Figure S2-15). A mixed effects linear model was employed to test whether genotypes showed consistent variation relative to the reference genotype RC, while considering development stages and environmental conditions on NPQ relaxation parameters. Nested model comparisons via AIC indicated 17 genotypes varied consistently in at least one parameter over six different models compared to RC (Figure 5). Three genotypes consistently varied in two parameters, with founders of NAM33 and 37 having lower A_qI_ and MaxNPQ compared to RC, while the NAM54 founder had slower relaxation of A_qM_ represented by a larger r_qM_, and a higher maximum NPQ (Figure 5).

**Figure 4.**
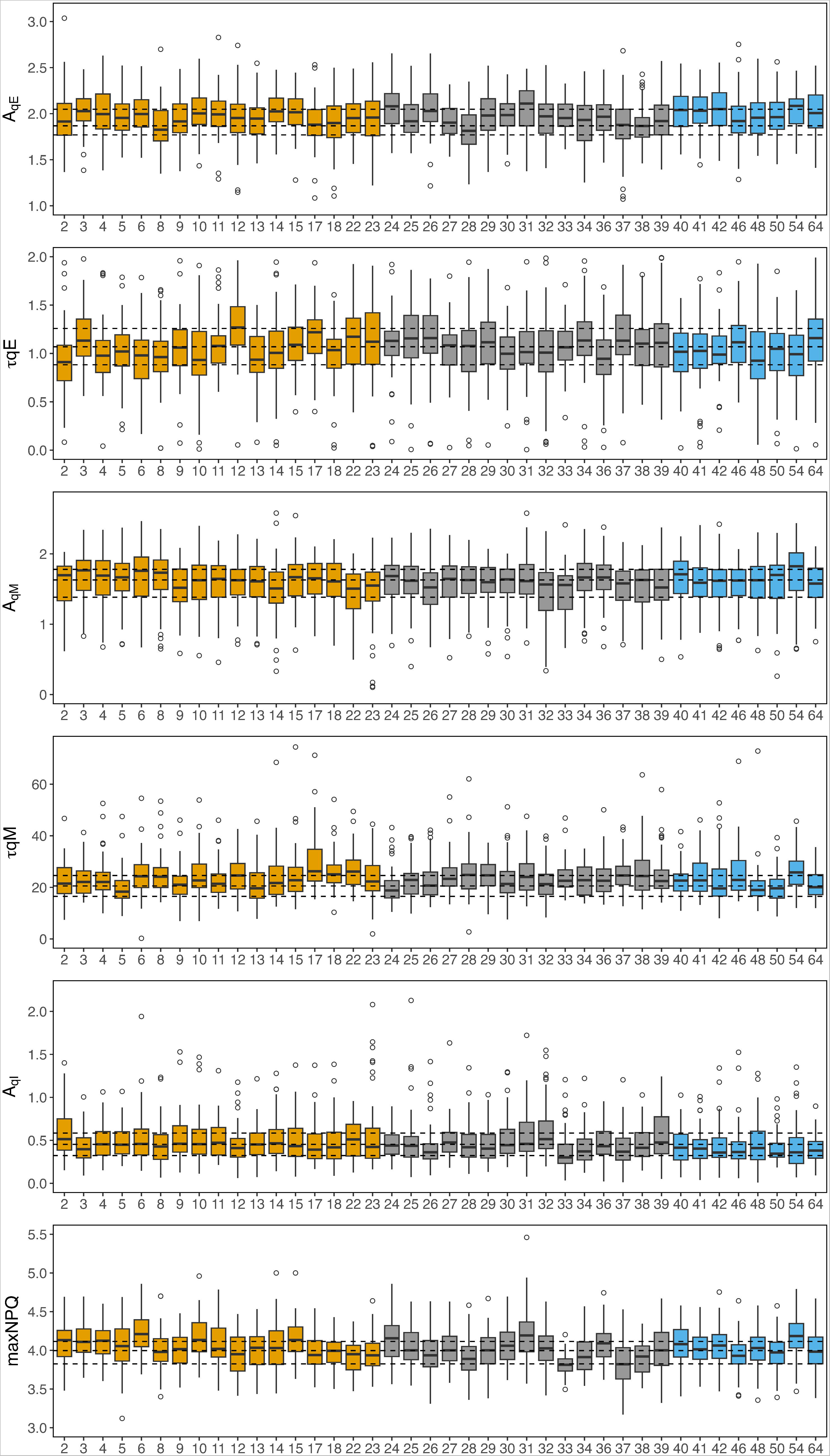
Comparison of NPQ relaxation kinetics in 41 founder genotypes of the SoyNAM population and common parent. Boxplots of six calculated NPQ relaxation parameters (A_qE_, A_qM_, A_qI_, r_qE_, r_qM_, maximum NPQ), plots are colored based on genotype group: Elite (yellow), Diverse (grey), PI (blue). Dotted black lines represent the median, upper and lower bounds of the interquartile range of reference line RC.

**Figure 5.**
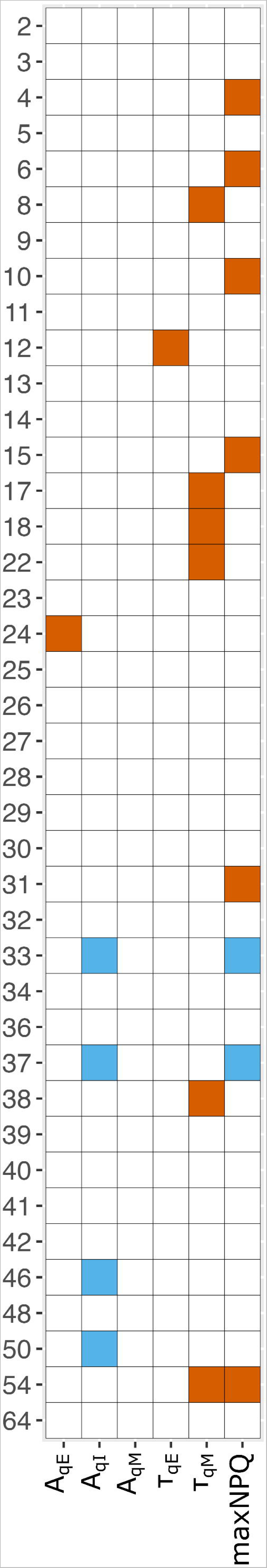
Summary representation of AIC trials for significant difference between genotypes and the common parent (RC). Genotypes which showed a consistently larger (orange) or smaller (blue) values in all six model comparisons are shown. Larger values for τ_qE_ and τ_qM_ represent slower relaxation.

**Table 5.**
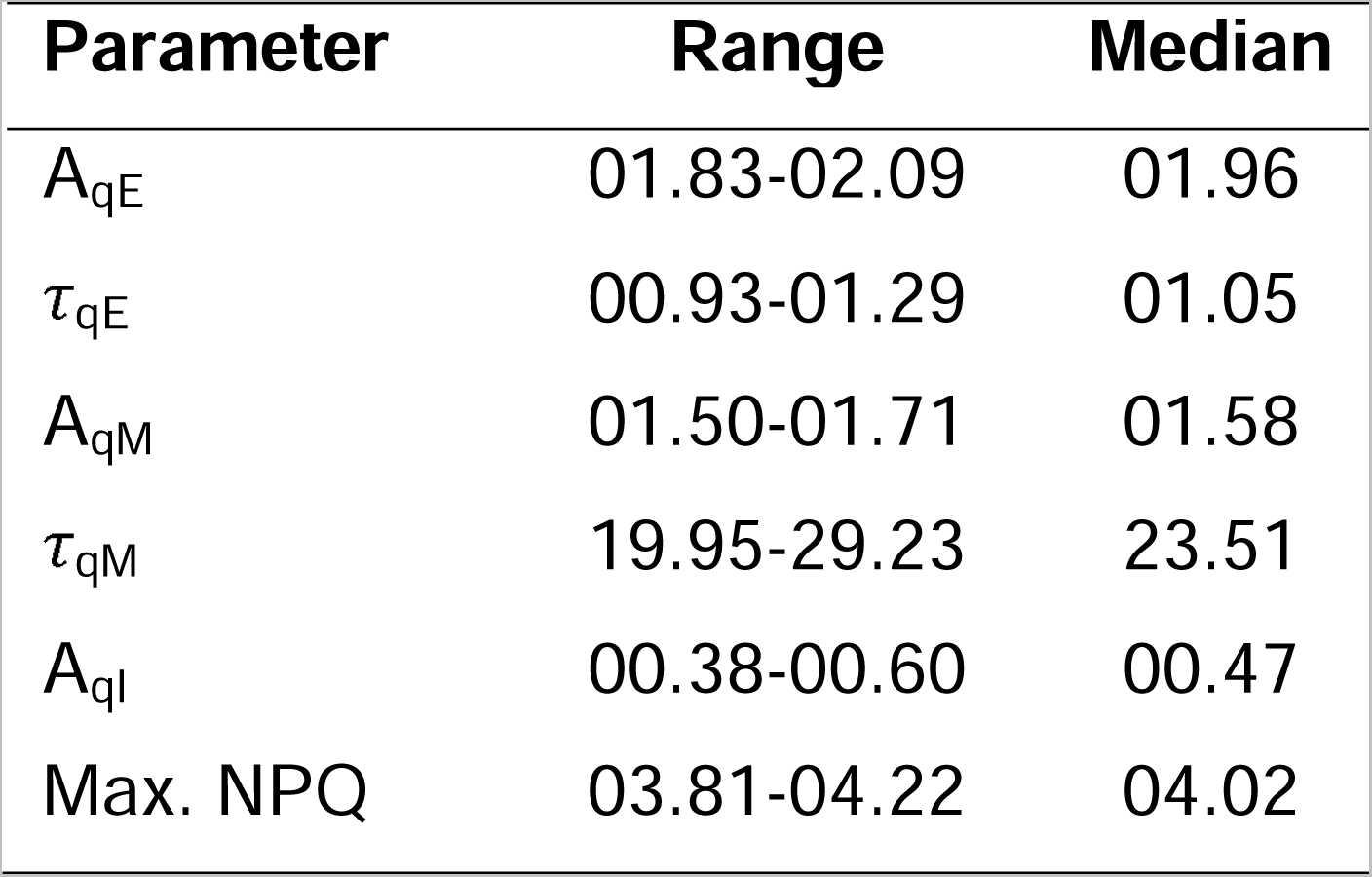
Summary of 41 NAM founder genotype NPQ relaxation parameters. Data represent comparison of mean values per genotype across all samples.

| Parameter | Range | Median |
| --- | --- | --- |
| $A_{qE}$ | 01.83-02.09 | 01.96 |
| $\tau_{qE}$ | 00.93-01.29 | 01.05 |
| $A_{qM}$ | 01.50-01.71 | 01.58 |
| $\tau_{qM}$ | 19.95-29.23 | 23.51 |
| $A_{qI}$ | 00.38-00.60 | 00.47 |
| Max. NPQ | 03.81-04.22 | 04.02 |

### Modeling impact of NPQ variation on canopy photosynthesis

To assess the potential for improving the efficiency of soybean photosynthesis a canopy model, parameterized for reference line RC was used to calculate the impact of varying NPQ parameters based on the phenotypic diversity present in the SoyNAM founders. The minimum and maximum recorded genotypic values for NPQ relaxation (r_qE_ and r_qM_) were used to estimate the potential decrease in CO_2_ assimilation caused by slow relaxation of NPQ on a cloudy (226) and sunny (227) day in Illinois in 2021 (Figure 6a and b). Simulations estimate the difference in CO_2_ assimilation between canopies with the fastest and slowest relaxing NPQ kinetics observed would equate to 1.6% on intermittently cloudy days, and 1.1% on a sunny day (Figure 6c).

**Figure 6.**
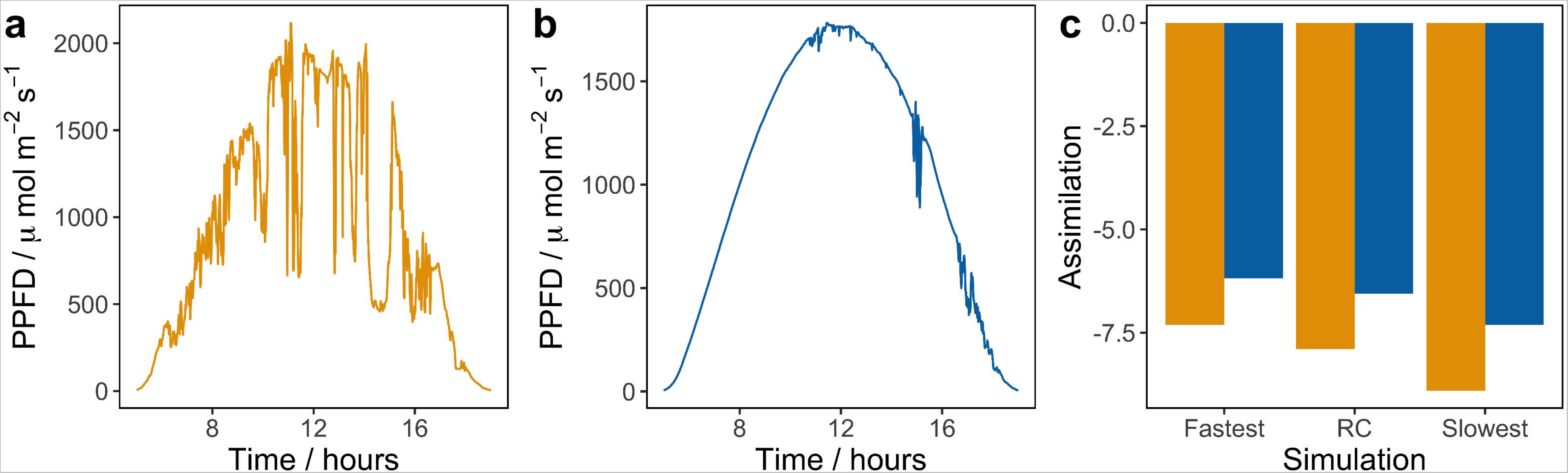
Canopy simulations comparing impact of SoyNAM phenotypic variability in NPQ relaxation kinetics on carbon assimilation on an intermittently cloudy day and sunny day. Measured light intensity on an intermittently cloudy day 266 (A) and sunny day 227 (B) in Illinois in 2021. (C) Representation of canopy model. (D) Illustrated are losses in Ac resulting from the measured rates of r_qE_ and r_qM_ comparing the slowest, fastest, mean and RC values from the SoyNAM population on a cloudy (orange bars) and sunny (blue bars) day.

## Discussion

The aims of this study included assessing the extent of variation in NPQ relaxation parameters between SoyNAM founder genotypes and predicting the impact on canopy photosynthesis. Given that we identified strong environmental effects on measured NPQ parameters, (Figure 2-3, Figure S1), our data are consistent with differences caused by physiological adaptation that lasts longer than 24h. This was brought to light by Figure S1, highlighting the importance of 7 day-averaged variables to explain the variation between years that daily averages couldn’t explain. The mechanisms behind this phenomenon are unclear, but it is possible that the phenotype is influenced by the status of xanthophyll cycle pigments in the leaf. Zeaxanthin (Zx), plays an important role in modulating NPQ and is created from the de-epoxidation of violaxanthin (Vx) via the intermediate antheraxanthin (Ax) (Jahns and Holzwarth, 2012). Zx decelerates the relaxation of qE (Niyogi *et al*., 1998) and the rate of conversion to Vx appears to be indirectly associated with slower relaxing phases of NPQ referred to as qZ (Nilkens *et al*., 2010; Kress and Jahns, 2017). The amount is Zx is dependent on the combined xanthophyll pool size (VAZ), which is regulated by carotene hydroxylase (Davison *et al*., 2002), and the steady state de-epoxidation state of xanthophyll pigments which is influenced by the activity of zeaxanthin epoxidase (ZEP). The VAZ pool size is adjusted in response to environmental conditions, with studies showing that leaves grown in full sun as opposed to shade conditions have larger pools (Demmig-Adams, 1998), with a greater de-epoxidation state, and more Zx would be expected to lead to an increase in r_qE_. However, there is currently a limited understanding of regulation of xanthophyll cycle enzymes in response to environmental stress, which appears to vary between tissues, species and cultivars (Schwarz *et al*., 2015; Bethmann *et al*., 2019; Grieco *et al*., 2020). For example, ZEP degraded in leaves but accumulated in roots of Arabidopsis exposed to drought stress (Schwarz *et al*., 2015). Whereas ZEP was shown to increase in abundance of leaves from an Iranian cultivar of wheat exposed to a water stress, but not one from the UK, while violaxanthin de-epoxidase (VDE) remained stable (Grieco *et al*., 2020). Interestingly, the wheat plants with increased ZEP were characterized by a higher qZ amplitude (Zx-dependent quenching on 10-30 min time scale) and a higher r_qZ_ relative to control as the drought treatment established, which would have been expected if VDE had increased or ZEP decreased. This has led the authors to hypothesize that the slowdown in NPQ was caused by a change in enzyme activity rather than in stoichiometry (Grieco *et al*., 2020). While it is difficult to determine whether the soybeans have been experiencing water stress during our study, the decline in r_qM_ (which would be the parameter the most related to r_qZ_) in 2022 coincided with precipitation events that had followed a sustained period with low precipitations (Figure 2) which would be consistent with increased ZEP activity. However, further investigation is required to determine if xanthophyll pigment content and enzyme abundance can account for the observed impact of other environmental variables on NPQ relaxation.

Fast conformational change in the thylakoid membrane is also known to play an important role in NPQ and its components (Schaller *et al*., 2010; Ruban *et al*., 2012; Sacharz *et al*., 2017). PsbS has been proposed to facilitate thylakoid membrane re-organization by regulating the interaction between LHCII and PSII which then results in NPQ (Kiss *et al*., 2008). A study conducted on a PsbS knock-out rice mutant revealed that r_qE_ depended on the level of phosphorylation of Lhcb1 and Lhcb2 (Pashayeva *et al*., 2021). High temperature has been shown to lead to an increase in the thylakoid membrane stiffness, causing a decline in the amplitude of the long-lifetime components of fluorescence decay without affecting its lifetime (Pollastri *et al*., 2019). While it difficult in our study to isolate the effect of air temperature on NPQ relaxation from the effect of other environmental variables, our results showed an important impact of temperature on NPQ relaxation and its varied components (Figure 2, Figure S1), though its effect varied depending on the year and the components. How multiple environmental stressors may interact and affect NPQ relaxation remains understudied.

Our study also revealed a seasonal pattern in NPQ relaxation (Figure 3) and the importance of aggregated environmental variables (Figure S1) when comparing NPQ relaxation between years, suggesting a lasting effect of the environment on NPQ relaxation. Photoinhibitory quenching has been shown to operate at a seasonal timescale (Demmig-Adams and Adams III, 2006) and could have a lasting impact on NPQ relaxation. Long-term adaptation of NPQ has been observed in *Taxus baccata* with needles exposed to high light in winter showing a slowed NPQ relaxation weeks after (Robakowski and Wyka, 2009). Studies have also shown that repeated excess-light exposure can lead to a faster onset kinetics of pH-dependent NPQ (Demmig-Adams *et al*., 2022). This was true for both sun-grown and shade leaves. Still, more studies are needed to understand how NPQ relaxation is affected by its environment on a longer timescale in crops.

The SoyNAM population is known to possess genotypic diversity with respect to photosynthetic variables: there is large variation in rates of rubisco activation, which was reported to cause a >5 fold difference in carbon fixation during the first five minutes following transition between dark and light conditions (Soleh *et al*., 2017), and loci have been identified influencing carbon assimilation and electron transport (Montes *et al*., 2022). Intriguingly the founder genotype of NAM12, which has previously been shown to possess highest levels of steady state electron transport (J_max_) (Montes *et al*., 2022), and slowest rate of rubisco activation (Soleh *et al*., 2017), was the only genotype that showed significantly slower rates of relaxation of qE relative to the reference (Figure 4). But further analysis of RIL population will be required to determine if these traits are linked or segregate independently. A known variant is found at the *e2* locus, which encodes homolog of the Arabidopsis circadian clock gene GIGANTEA (Watanabe *et al*., 2011). This gene have been shown to have pleiotropic effects which influence soybean photosynthesis (Montes *et al*., 2022), canopy coverage (Xavier *et al*., 2017) and yield traits (Diers *et al*., 2018). The late maturity *E2* allele is segregating in seven of the NAM founders, three of which (NAM33, 37 and 50) were identified as having significantly less A_qI_ compared to the reference line, and it may therefore also influence the traits measured here.

Although a small number of lines were found to vary in NPQ kinetics compared to RC (Figure 4), the overall diversity in NPQ relaxation parameters between SoyNAM founders was limited, for example, only a 1.5-fold change between the smallest and highest r_qM_ (Table 5). This small amount of variation is not a general phenomenon related to NPQ, with several studies finding substantial variation in species ranging from Arabidopsis to Maize (Jung and Niyogi, 2009; Rungrat *et al*., 2019; Cowling *et al*., 2022; Sahay *et al*., 2023). For example, analysis of rice genotypes found large variation maximum NPQ (Kasajima *et al*., 2011), which was attributed to an insertion in the promoter region of photosystem II subunit S (PsbS) resulting in higher expression and higher NPQ in japonica rice (Wang *et al*., 2017). In the case of soybean, this lack of variation may reflect an unusual number of genetic bottlenecks in domestication and then introduction into the USA (Hyten *et al*., 2006), with only 28 ancestors contributing to 95% of the genes in cultivars released between 1947-1988 (Gizlice *et al*., 1994). The lines used in this study were chosen for their diversity, but it remains unclear if there is more variation in NPQ to be found in the wider soybean germplasm and whether more may be found in collections of the wild ancestor *Glycine soja* which is considered a largely untapped source of genetic variation in cultivated soybean (Kofsky *et al*., 2018).

Previous transgenic manipulations suggested that speeding up relaxation of NPQ can increase photosynthetic efficiency leading to improved growth (Kromdijk *et al*., 2016) or seed production (De Souza *et al*., 2022), although it is not always the case (Garcia-Molina and Leister, 2020; Lehretz *et al*., 2022). The reason for this is not clear, but the interplay between total photoprotection, ability to use assimilate, and the rates of activation and relaxation are likely to be important. Wang et al. (2020) previously used a ray tracing algorithm and canopy model to estimate the predicted benefit of manipulating NPQ or photosynthetic induction rates in soybean given available natural variation. The data used was based on preliminary measurements made on the same SoyNAM panel as in this publication (Song *et al*., 2017; Diers *et al*., 2018). However, predictions of the impact on altering NPQ relaxation were based on light-to-dark measurements, which are shown to be unlikely to be realistic for a field environment (Fig. 1B) and did not focus on identifying which soybean genotypes could be beneficial for genomic selection from the perspective of NPQ relaxation. Here, the differences in NPQ parameters between lines was smaller, ∼28% difference in r_qE_ between fastest and slowest relaxing lines as compared to 40% (Wang *et al*., 2020), which equates to 1.6% daily assimilation, likely due to the much deeper sampling and measurement conditions. Analysis of the response of soybean to artificial increase of photosynthesis in the field by elevation of [CO_2_], show that while older cultivars appear sink limited, modern cultivars appear to be source limited and can fully use any increase in photosynthesis (Ainsworth and Long, 2021). Given the limited diversity of NPQ relaxation rates in soybean germplasm, achieving increases in photosynthetic efficiency through manipulating NPQ is likely to be best achieved through transgenic approaches.

## Experimental procedures

### Plants and growth conditions

The 41 parents of the Soybean Nested Association Mapping (NAM) population (Song et al., 2017, Diers et al., 2018) were grown in the field at the Crop Sciences Research and Education Center at the University of Illinois at Urbana-Champaign in 2021 (latitude 40.084604, longitude −88.227952) and 2022 (latitude 40.064866, longitude −88.193084). Seeds were planted in 1.2 m single-row plots with a 0.75 m row spacing, with 40 seed m^-1^ in a North-South orientation on 5 June 2021 and 13 June 2022 (Table S4). The experiment was arranged in a randomized complete block design, with five replicate plots per genotype. Standard agronomic practices were employed.

For greenhouse experiments, seeds of the NAM Recurrent Parent IA3023 were planted 2 mm deep in a Soil: Perlite:Torpedo Sand mixture (1:1:1), in classic 600 2-gallon pots on May 5^th^, 2021. Plants were watered and fertilized as needed, and grown under 1500 μmol m^−2^ s^−1^ light, 14 h daylength, with temperatures set a 27-30°C during the day, 23-26°C at night.

### Meteorological data collection

Meteorological variables were measured every 30 min by a weather station at the University of Illinois Energy Farm, approximately 1 km from the Crop Sciences Research and Education Center (latitude 40.062832, longitude −88.198417). Air temperature (T_a_, °C) and relative humidity (RH, %) were recorded by a HMP45C probe (Campbell Scientific, Logan, UT, USA), and incoming shortwave radiation (Fsd, W m^-2^) was from a CNR1 radiometer (Kipp & Zonen, The Netherlands), both instruments were installed 4 m above the ground. T_a_ and RH were used to calculate saturation vapor pressure (e_s_) and actual vapor pressure (e_a_) for each 30 min period, which were then used to calculate vapor pressure deficit (VPD, kPa) as per Equations 1-3:

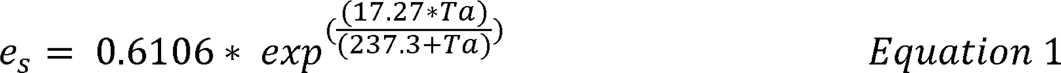

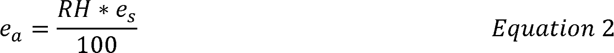

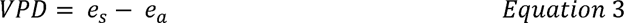

Occasional gaps in meteorological data are inevitable when measuring at these time scales, so data gaps were filled where needed, whereby an artificial neural network was used to generate a complete time-series with external data sourced from the University of Illinois Willard Airport weather station (station ID: 725315-94870) and ERA-interim data from the European Centre for Medium Range Forecasts. In total, less than 5 % of data required gap filling. Daily summary values were then calculated for each variable, with T_a_ and VPD presented as daily mean, Fsd as daytime-only (i.e. 06:00-18:00) mean, and rainfall as a daily sum. The variables T_a__7day, VPD_7day, and Fsd_7day represent the average for the past 7 days for these values. The variable Precip_7day represent the sum in precipitation for the past 7 days. The cumulated net radiation (cumFn) was calculated as the sum of net radiation between 6:00 and 12:00

### Chlorophyll fluorescence analysis

For field experiments, plants were sampled between 8:00-10:00 at seven time points across development in 2021 and six time points in 2022 (Table S5) according to (Gotarkar *et al*., 2022). Briefly, five 4.8 mm leaf disks were collected per plot, sampling from the upper-most mature leaf using a cork borer (H9663; Humboldt Mfg. Co.). Leaf disks were then transferred face down into wells of a flat-bottomed 96 well plate (FB012929; Fisher Scientific), humidity was maintained by placing a half-wet nasal aspirator filter into each well (iHank-Nose B07P6XCTGV; Amazon), and plates were sealed and wrapped in aluminum foil followed by overnight dark adaptation. Measurements were taken with modulated chlorophyll fluorescence imaging system (CF imager; Technologica, UK). To induce low-high-low light fluctuations, samples were illuminated for 10 min at 50 μmol m^−2^ s^−1^, followed by 15 min at 2000 μmol m^−2^ s^−1^ and 50 min of 50 μmol m^−2^ s^−1^. *F* _m’_, was determined by applying saturating pulses (4000 μmol m^−2^ s^−1^) at 2.5, 5, 7.5, 10 min after the actinic light was turned on (50 μmol m^−2^ s^−1^), 2.5, 5, 7.5, 10, 12.5 and 15 min after high light exposure (2000 μmol m^−2^ s^−1^), and 2.5, 5, 10, 15, 20, 25, 30, 35, 40, 45 and 50 min following return to low light (50 μmol m^−2^ s^−1^). The background was excluded manually and NPQ values at each pulse were calculated.

NPQ values were calculated for each time point using custom MatLab scripts according to (Gotarkar *et al*., 2022). NPQ relaxation parameters A_qE_, A_qM_, A_qI_, r_qE_ and r_qM_ were then calculated by fitting the sum of a double exponential function to measured NPQ values following shut off of the actinic light, according to Equation 4 (Dall’Osto *et al*., 2014):

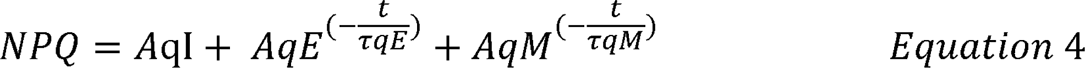

using the fit function in MatLab R0218b, where t is the measured fluorescence at a given time point. Maximum NPQ values are defined as the maximum value reached during the 15 min illumination at high light.

### Measurement of light intensity at layers in the soybean canopy

Light intensity was measured simultaneously within and above a soybean canopy using quantum sensors (LI-190R, LICOR, Lincoln, NE, USA) mounted on an aluminum frame reading at 8 different levels (9, 24, 44, 64, 84, 104, 124, and 144 cm height). Measurements were manually logged on a datalogger (CR100x; Campbell Scientific, Logan, UT, USA) after carefully placing the frame within the canopy with the sensors close to the stems and waiting for stable readings. Those measurements were taken on six different cultivars from a germplasm collection at (or close to) the reproductive stage R5. Measurements were conducted on August 17th, 2022 between 12:00 and 12:15.

### 3D soybean model and light distribution

The dynamics of lighting within a soybean canopy were predicted with a 3D architectural representation, using our previously presented soybean canopy model (Song et al.,2019; Wang et al., 2020). The model was parameterized on the measured architecture of soybean hub parent IA3023 (RC) at the University of Illinois Energy Farms in August 2022 (measured canopy parameters are listed in Tables S6-8). Leaf area was measured when the soybean plants on 18^th^ August 2021. The youngest mature leaf (∼3^rd^ trifoliate from the top) was selected for analysis, the area of all three leaves in the trifoliate from three plots was measured with a leaf area meter (LI3100C; LI-COR Environmental, Lincoln, NE, USA) (Table S6). Detailed parameters were measured for 5 genotypes on August 19^th^ 2021 and 8 genotypes on August 30^th^ 2022. Plant height was measured from the base to tip after the plants were cut from the base and stretched. Leaf width was measured at the widest point from each leaflet in a trifoliate and averaged. Leaf length was measured from base to tip for each trifoliate and averaged. Internode length was measured for the 6^th^ internode from the top, with one value recorded per plant, and branch angle was measured for the 6^th^ branch from the top using a digital protractor, one measurement per plant. The total number of trifoliate, number of primary and secondary branches and number of pods per plant were counted manually for one plant per plot (Table S7). Leaf Area Index (LAI) was measured using the SunScan canopy analysis system (SS1; Delta-T Devices Ltd, UK) with a 1 m probe according to the manufacturer’s instructions.

LAI was measured at R5 developmental stage between 11:56 and 13:50 on 20^th^ August 2021 on both sides of each plot and averaged, with the probe positioned parallel to the rows (Table S8). To calculate the actual light environment of the soybean canopy, measured PAR data on DOY 226 and 227 of 2021 in Bondville, IL. (Earth System Research Laboratory, Global Monitoring Division https://www.esrl.noaa.gov/gmd/grad/surfrad/dataplot.html) was incorporated (Table S9). A forward ray-tracing algorithm (FastTracer; Song et al. 2013) was used to predict the light absorption of each leaf pixel (ca. 5 mm^2^) every 1min from 05:00 – 19:00 in Champaign IL, US (40.11N, 88.21 W).

### Simulation of dynamic photosynthesis

Dynamic photosynthetic rates were calculated for every 10 s (Δ*t*) of the day using the absorbed light for each leaf pixel, considering rates of Rubisco activation and NPQ relaxation (Wang et al., 2020). Then the canopy net CO_2_ uptake (*A*_c_) was calculated as:

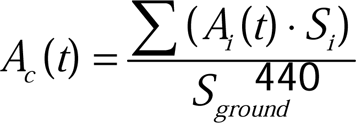

Where *A_i_(t)* is the CO_2_ uptake rate of a leaf pixel; *S_i_* is the surface area of each pixel, *S_ground_*represents the occupied ground area of the simulated canopy. All simulations were conducted in MATLAB 2021a (The Mathworks, Inc^®^).

Time constants of NPQ relaxation and Rubisco activation across the NAM population were measured and used as input of the dynamic photosynthetic model (model inputs were listed in Table S10). The time constant of Rubisco de-activation was assumed to be double the time required for activation for each genotype (Taylor & Long, 2017).

### Statistical Analysis

Technical replicates from the chlorophyll fluorescence analysis were averaged prior to statistical analyses. The impact of genotypes, the environment, and their interaction on NPQ relaxation was assessed by fitting an ANOVA model in R v4.1.2 (R Core Team, 2016) and RStudio v2024.04.0 (RStudio Team, 2015) using the stats R package v4.1.2. The ANOVA model was written as NPQ parameter ∼ genotype (G) + day (or the environment, E) + genotype * day with G and E set as fixed factors. The impact of the environment was assessed for each NPQ relaxation parameter by using a stepwise regression model to determine which environmental variables had the strongest impact on the parameters. The initial linear model included all the environmental variables. A both-direction stepwise algorithm then tested the model by removing and re-including environmental variables one by one. The Akaike’s information criterion (AIC) was used to select the best minimum adequate model (lowest AIC) using the stats R package v4.1.2. This test was performed with data collected in 2021, 2022, and for the two years together. Centered and scaled environmental variables were used as input in the model. The relationships between genotypes, the environment, and NPQ relaxation were explored using canonical correlation analysis (CCA). This multivariate statistical approach identifies linear combinations of variables for the two datasets (here, NPQ and environmental data) to construct a pair of canonical variates that are maximally correlated with each other on the first canonical axis (CC1). Then, a second pair of canonical variates is made with a maximized correlation on the second canonical axis (CC2) but uncorrelated with the CC1. The process is repeated for each canonical variates’ pairs. A significant correlation between pairs enables associations among the different variables. The CCA analysis was performed in R using the CCA R package v1.2.1 (González and Déjean, 2021). The statistical significance of canonical correlation coefficients was carried out by using Wilks’ Lambda with the CCP R package v1.2 (Menzel, 2022). All NPQ relaxation parameters were used as input for the first dataset. Environmental variables were used as input for the second dataset with the exception of VPD_7day, Precip_7day, Fsd_7day, and cumulated precipitation based on the stepwise regression model to reduce co-linearity.

For chlorophyll fluorescence analysis, leaf disks with *F*_v_/*F*_m_ value <0.75 were excluded to ensure measured values were from healthy disks representative of whole plant kinetics. Additional outliers were defined and removed if calculated parameter values were <0. The effect of genotypic differences from the RC for each NPQ relaxation parameter were estimated using linear mixed-effects models (Bates *et al*., 2015). Linear fixed effects included days, air temperature (Ta), vapor pressure deficit (VPD), precipitation (Precip), incoming shortwave radiation (Fsd), a 7-day rolling mean air temperature (Ta 7day), and a 7-day rolling mean vapor pressure deficit (VPD 7day), with random effects for plot and leaf disks (Data S1). Akaike’s Information Criterion (AIC) was used to compare linear mixed-effects models (Akaike, 1974; Faraway, 2016). Briefly, nested model comparisons via AIC determined which genotypes exhibit different measures from the baseline genotype RC, following a procedure to fit a smaller model with one genotype removed. This smaller model assumes that there is no difference between that genotype and the RC baseline. The computed AIC value for this smaller model was compared with the AIC value of the full model with all genotypes considered. If the smaller model has a lower AIC value, then the removed genotype does not exhibit an effect on NPQ relaxation parameters that is different from the RC baseline. This procedure was repeated for all genotypes and all linear mixed-effects models involving NPQ relaxation parameters as a response. In addition to this approach, a principal component analysis (PCA) was performed for each days separately to identify any patterns among NAM groups (i,e, diverse, elite, PI, and RC) and/or notable genotypes. The analysis was performed using the ade4 (v1.7-22) R package (Dray and Dufour, 2007). Centered and scaled NPQ parameters were used as input variables. Differences between NAM groups in their principal component coordinates were assessed by using a one-way ANOVA with groups as a fixed factor with the R stats package (v4.1.2).

### Data Statement and Accession Numbers

All raw NPQ data values prior to filtering and processing are provided in csv format in Table S1. All raw chlorophyll fluorescence imager files and custom scripts can be accessed via FigShare 10.6084/m9.figshare.21574509 and 10.6084/m9.figshare.25939504.

## Acknowledgements

We thank Troy Cary for setting up the soybean NAM population plots; Wanne Kromdijk for help setting up scripts for analysis of NPQ relaxation parameters; Caitlin Moore for assistance with data from 2018 and 2019 used in previous versions of this manuscript; Meghan Burns, Meghan Blaszynski, Jacob Milo, and Abigail Hinkle for their assistance collecting samples and Stephen P. Long for commenting and feedback on the manuscript. This work was supported by the project Realizing Increased Photosynthetic Efficiency (RIPE), funded from 2017-2023 by the Bill and Melinda Gates Foundation (BMGF), Foundation for Food and Agriculture Research (FFAR) and the UK Department for International Development (UKAid) under grant number OPP117215. SJB was supported by Carl R. Woese Institute for Genomic Biology Fellowship.

## Short legends for Supporting Information

Table S1: Raw NPQ timeseries values for all technical replicates prior to filtering and processing.

Table S2. Calculated NPQ relaxation parameters for all plots and genotypes.

Table S3. Genotype means for calculated NPQ relaxation parameters combining all data from 2021 and 2022.

Table S4: Field design with plot numbers and orientation in 2021 and 2022.

Table S5. Sampling time points in 2021 and 2022 including days after sowing and development stage.

Table S6. Leaf area measurements for 41 SoyNAM founders recorded in 2022, data is provided for three biological replicates.

Table S7. Canopy measurements for seven SoyNAM founders and reference genotype RC recorded on August 30^th^ 2022.

Table S8. Leaf area index measurements for 41 SoyNAM founders recorded in 2022, data is provided for two technical and 5 biological replicates.

Table S9. Measured PAR values on day of the year (DOY) 226 and 227 of 2021 in Bondville, IL. Earth System Research Laboratory, Global Monitoring Division.

Table S10. Input values for 3D canopy model.

Data S1. Description of statistical analysis comparing NPQ relaxation kinetics between genotypes.

## Supplemental Figure Legends

Figure S1. Coefficients for the best minimum adequate model (lowest AIC) for each NPQ relaxation parameters in 2021 and 2022 combined. The R2 and p-value are shown for each model.

Figure S2. PCA of NPQ relaxation in the NAM population on 6/24/21. Diverse, elite, and PI lines are shown in grey, orange, and blue, respectively. Coordinates for replicate measurements on the same line were averaged. RC line is shown in black. Error bars represent the standard deviation. Lines with a z-score>2.5 for one of their components were labeled.

Figure S3. PCA of NPQ relaxation in the NAM population on 6/30/21. Diverse, elite, and PI lines are shown in grey, orange, and blue, respectively. Coordinates for replicate measurements on the same line were averaged. RC line is shown in black. Error bars represent the standard deviation. Lines with a z-score>2.5 for one of their components were labeled.

Figure S4. PCA of NPQ relaxation in the NAM population on 7/7/21. Diverse, elite, and PI lines are shown in grey, orange, and blue, respectively. Coordinates for replicate measurements on the same line were averaged. RC line is shown in black. Error bars represent the standard deviation. Lines with a z-score>2.5 for one of their components were labeled.

Figure S5. PCA of NPQ relaxation in the NAM population on 7/13/21. Diverse, elite, and PI lines are shown in grey, orange, and blue, respectively. Coordinates for replicate measurements on the same line were averaged. RC line is shown in black. Error bars represent the standard deviation. Lines with a z-score>2.5 for one of their components were labeled.

Figure S6. PCA of NPQ relaxation in the NAM population on 7/20/21. Diverse, elite, and PI lines are shown in grey, orange, and blue, respectively. Coordinates for replicate measurements on the same line were averaged. RC line is shown in black. Error bars represent the standard deviation.

Figure S7. PCA of NPQ relaxation in the NAM population on 7/28/21. Diverse, elite, and PI lines are shown in grey, orange, and blue, respectively. Coordinates for replicate measurements on the same line were averaged. RC line is shown in black. Error bars represent the standard deviation.

Figure S8. PCA of NPQ relaxation in the NAM population on 8/11/21. Diverse, elite, and PI lines are shown in grey, orange, and blue, respectively. Coordinates for replicate measurements on the same line were averaged. RC line is shown in black. Error bars represent the standard deviation.

Figure S9. PCA of NPQ relaxation in the NAM population on 7/6/22. Diverse, elite, and PI lines are shown in grey, orange, and blue, respectively. Coordinates for replicate measurements on the same line were averaged. RC line is shown in black. Error bars represent the standard deviation.

Figure S10. PCA of NPQ relaxation in the NAM population on 7/12/22. Diverse, elite, and PI lines are shown in grey, orange, and blue, respectively. Coordinates for replicate measurements on the same line were averaged. RC line is shown in black. Error bars represent the standard deviation.

Figure S11. PCA of NPQ relaxation in the NAM population on 7/19/22. Diverse, elite, and PI lines are shown in grey, orange, and blue, respectively. Coordinates for replicate measurements on the same line were averaged. RC line is shown in black. Error bars represent the standard deviation.

Figure S12. PCA of NPQ relaxation in the NAM population on 7/26/22. Diverse, elite, and PI lines are shown in grey, orange, and blue, respectively. Coordinates for replicate measurements on the same line were averaged. RC line is shown in black. Error bars represent the standard deviation.

Figure S13. PCA of NPQ relaxation in the NAM population on 8/4/22. Diverse, elite, and PI lines are shown in grey, orange, and blue, respectively. Coordinates for replicate measurements on the same line were averaged. RC line is shown in black. Error bars represent the standard deviation.

Figure S14. PCA of NPQ relaxation in the NAM population on 8/20/22. Diverse, elite, and PI lines are shown in grey, orange, and blue, respectively. Coordinates for replicate measurements on the same line were averaged. RC line is shown in black. Error bars represent the standard deviation.

Figure S15. PCA displaying the relationships between the NPQ relaxation parameters and the environment in (a-d) 2021 (1296 observations) and (e-h) 2022 (1025 observations). (a, c, e, and g) PCA showing the spatial distribution of the different observations on the principal components (PC). The circular shapes represent observations, with different colors representing different days. The triangular shapes show the average of different NAM groups at different day, with diverse, elite, PI, and RC groups shown in grey, orange, blue and black, respectively. For each group, a solid line connect those points to represent their evolution throughout the season. (b, d, f, and h) The correlation circle showing the relationships between variables. Variables related to the NPQ relaxation parameters, and the environments are represented in red and black, respectively. The percentage of total variance explained by the PC1, PC2, and PC3 is shown on the axis title.

