## Supplementary material for "Variation in relaxation of non-photochemical quenching between the founder genotypes of the soybean (Glycine max) nested association mapping population": Data S1

### Soybean Analysis with Random Effects (2021-2022)

Daniel J. Eck

### Initial preprocessing

We first walk through initial preprocessing steps before we get to our visualizations. We load in the following R packages

```
library(tidyverse)
library(lubridate)
library(lme4)
library(Renvlp)
library(parallel)
```

We load in the data and perform some manipulations such as averaging across technical replicates.

```
CFdat2022 = read_csv("CF_and_weather_data_2021_2022.csv")
#str(CFdat2022)
CFdat2022$ID[grepl("RC", CFdat2022$ID)] = "aRC"
CFdat2022$ID = gsub(" ", "", CFdat2022$ID)

dat = CFdat2022 %>%
  group_by(plot_number, Date, ID) %>%
  summarise(mAqI = mean(AqI),
            mAqE = mean(AqE),
            mtqE = mean(tqE),
            mAqM = mean(AqM),
            mtqM = mean(tqM),
            maxNPQ = mean(maxNPQ),
            ID = unique(ID),
            plot_number = unique(plot_number),
            Date = unique(Date),
            Ta = unique(Ta),
            VPD = unique(VPD),
            Precip = unique(Precip),
            Fsd = unique(Fsd),
            Ta_7day = unique(Ta_7day),
            VPD_7day = unique(VPD_7day),
            Precip_7day = unique(Precip_7day),
            Fsd_7day = unique(Fsd_7day),
            Precip_7day_sum = unique(Precip_7day_sum),
            Precip_cum = unique(Precip_cum)) %>%
  mutate(year = year(mdy(Date))) %>%
  mutate(plot_number_year = paste(plot_number, year, sep = "_"))

## Precip_7day_sum is redundant
#dat %>%
#  mutate(ratio = Precip_7day/Precip_7day_sum) %>%
#  pull(ratio) %>% table()
dat = dat %>% select(-Precip_7day_sum)
```

We remove some extreme positive values and all negative values from the data set. We remove the anomalous 07/16/2021 values from the data set. Code that explores extreme values is in the accompanying .Rmd file.

```
dat2 = dat
dat2 = dat2 %>%
  filter(Date != "7/16/21") %>%
  filter(mAqI >= 0) %>%
  filter(mAqE >= 0) %>%
  filter(mtqE <= 2000) %>%
  filter(mtqM <= 2000) %>%
  filter(mtqE <= 2) %>%
  filter(mAqM <= 1000, mAqM >= 0) %>%
  mutate(Date = as.Date(Date, format = "%m/%d/%y"))
dat2$Date_num = dat2 %>% dplyr::select(Date) %>%
  mutate(Date_num = as.numeric(Date)) %>%
  pull(Date_num) %>% scale() %>% as.numeric()
```

We remove the last date of data collection for both years. The responses for these last dates are expected to be anomalous due to changing development stages over the course of data collection. We scale weather variables.

```
last_day = dat2 %>% group_by(year) %>%
  summarise(last_day = max(Date_num)) %>%
  pull(last_day)

dat3 = dat2 %>%
  filter( !(year == 2021 & Date_num == last_day[1]) ) %>%
  filter( !(year == 2022 & Date_num == last_day[2]) ) %>%
  ungroup()

dat4 = dat3 %>%
  mutate(Ta = as.numeric(scale(Ta)),
         VPD = as.numeric(scale(VPD)),
         Precip = as.numeric(scale(Precip)),
         Fsd = as.numeric(scale(Fsd)),
         Ta_7day = as.numeric(scale(Ta_7day)),
         VPD_7day = as.numeric(scale(VPD_7day)),
         Precip_7day = as.numeric(scale(Precip_7day)),
         Fsd_7day = as.numeric(scale(Fsd_7day)),
         Precip_cum = as.numeric(scale(Precip_cum)))
```

Prior analyses suggested that there were problematic data points in the modeling of maxNPQ. This could be the result of a faulty disc. We remove these points for all responses.

```
m1_maxNPQ = lmer(maxNPQ ~ ID + Date_num + I(Date_num^2) + Ta + VPD + Precip + Fsd +
             Ta_7day + VPD_7day + Precip_7day + Fsd_7day  +
             Precip_cum + (1|plot_number_year),
           data = dat4, REML = FALSE, control = lmerControl(optimizer ="Nelder_Mead"))

dat5 = dat4 %>%
  mutate(resid = residuals(m1_maxNPQ)) %>%
  filter(resid < 2) %>%
```

```
  dplyr::select(-resid)
```

We investigate the marginal means for each response across date and year.

##### AqE response

```
## Last day of data collection may be worth removing due to plants senescening
## More thorough data analysis needed (worth looking at Date)
ggplot(dat5 %>%
         group_by(Date) %>%
         summarise(mAqE = mean(mAqE), year = unique(year)),
       aes(x = Date, y = mAqE)) +
  geom_point() +
  geom_line() +
  facet_wrap(~year, scales = "free_x")
```

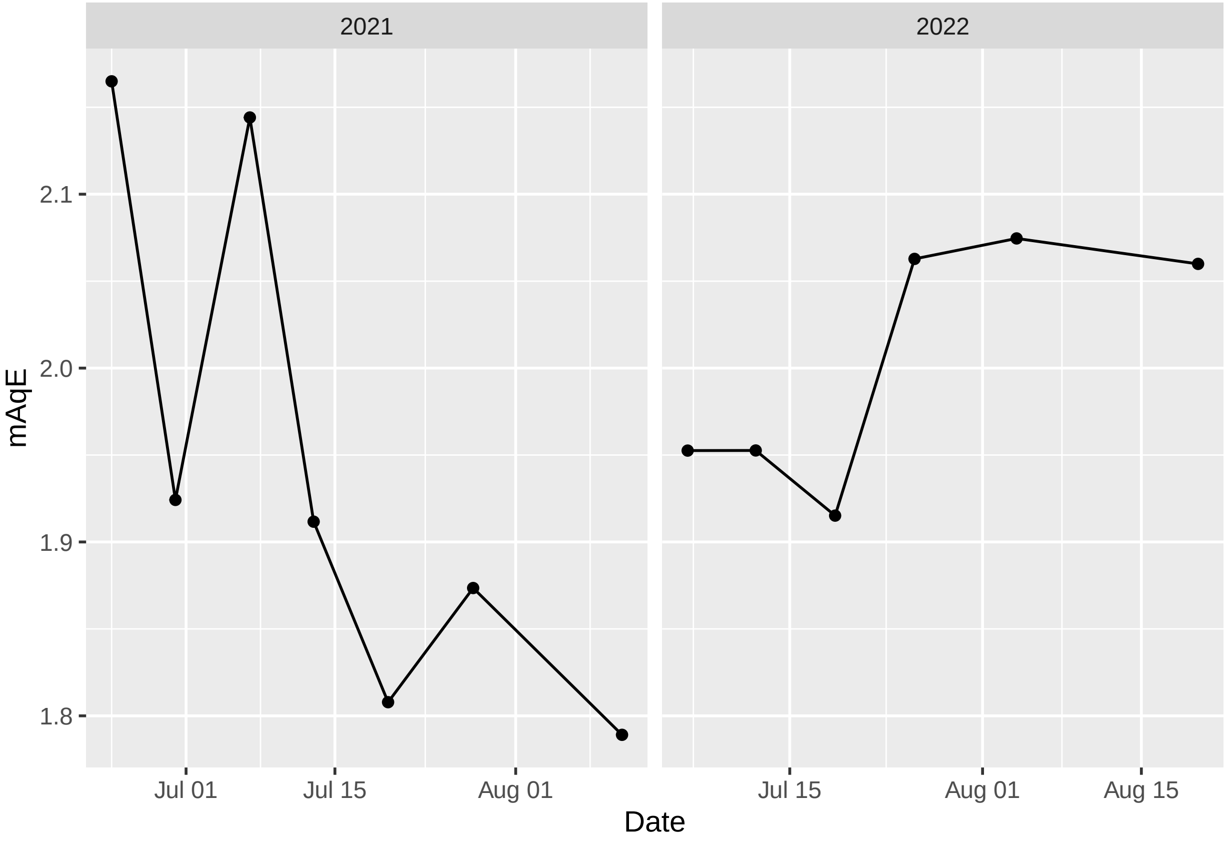

##### AqI response

```
ggplot(dat5 %>%
         group_by(Date) %>%
         summarise(mAqI = mean(mAqI), year = unique(year)),
       aes(x = Date, y = mAqI)) +
  geom_point() +
  geom_line() +
  facet_wrap(~year, scales = "free_x")
```

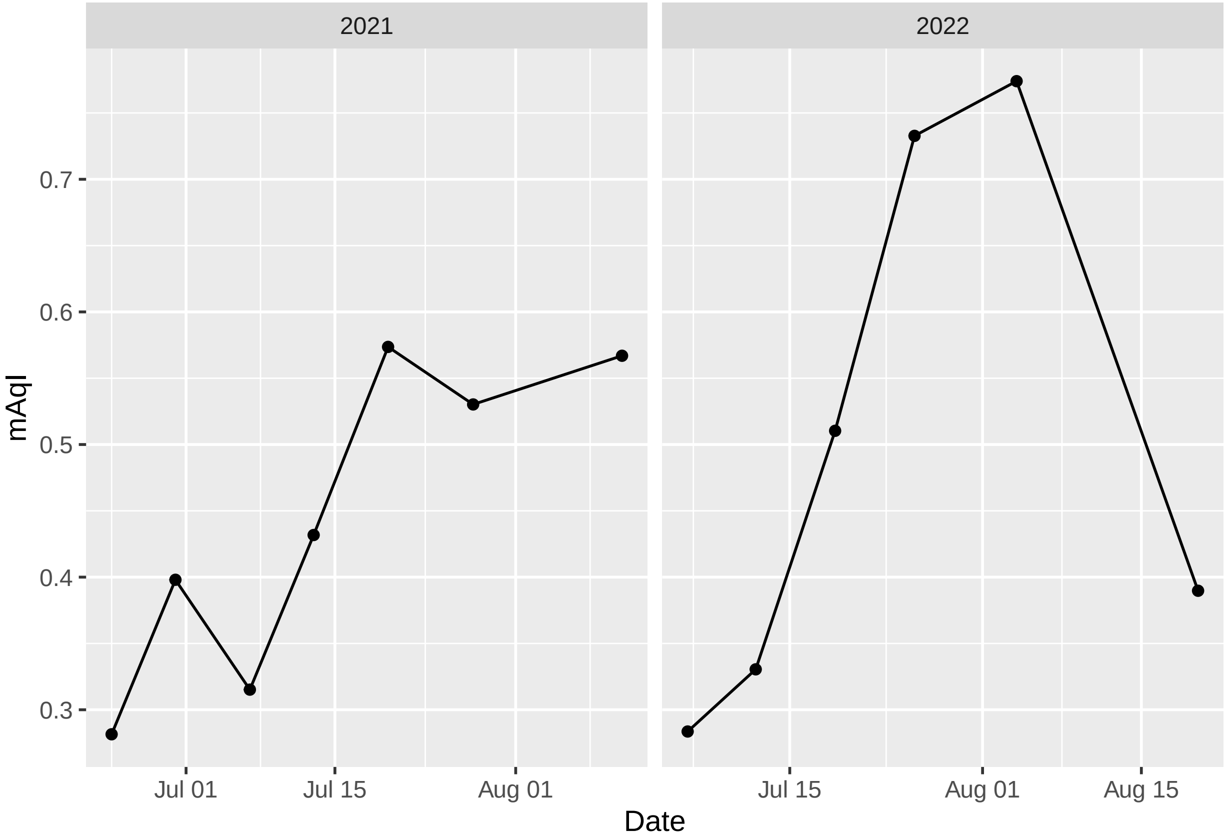

##### AqM response

```
ggplot(dat5 %>%
         group_by(Date) %>%
         summarise(mAqM = mean(mAqM), year = unique(year)),
       aes(x = Date, y = mAqM)) +
  geom_point() +
  geom_line() +
  facet_wrap(~year, scales = "free_x")
```

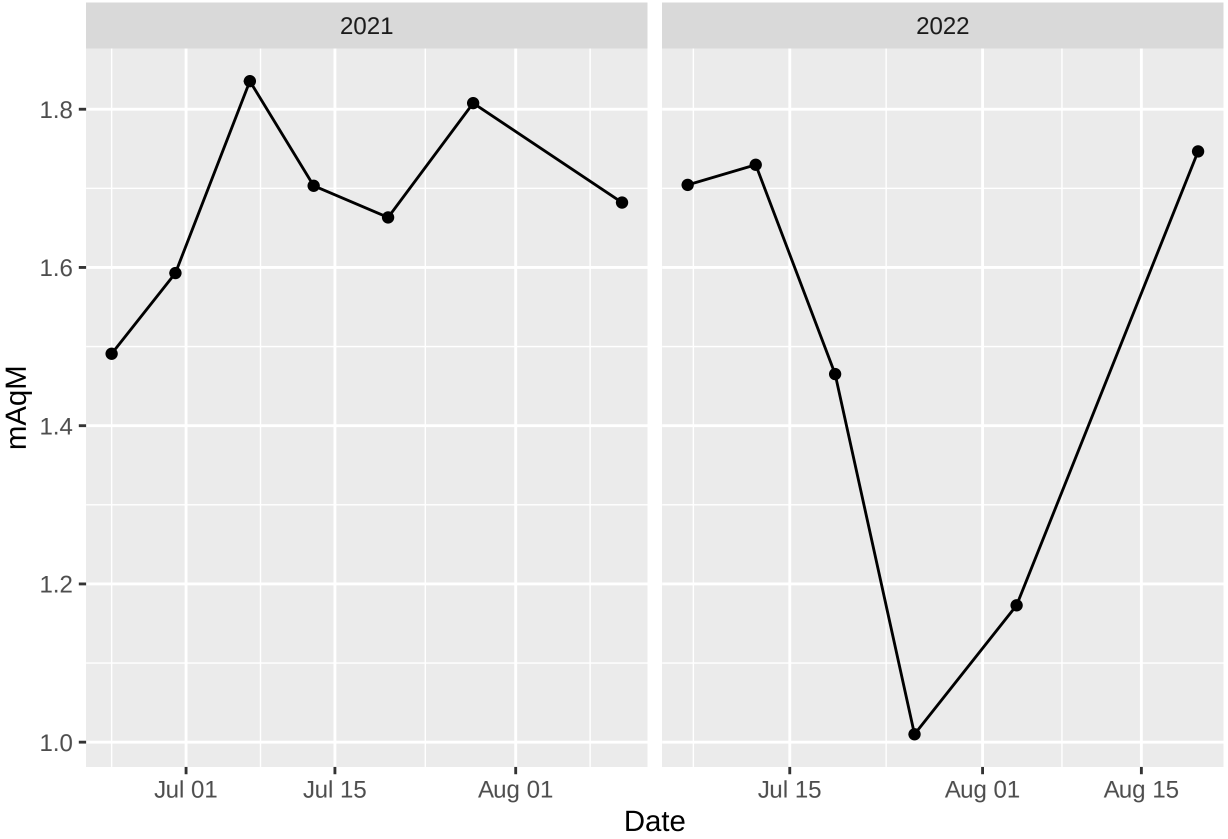

##### tqE response

```
ggplot(dat5 %>%
         group_by(Date) %>%
         summarise(mtqE = mean(mtqE), year = unique(year)),
       aes(x = Date, y = mtqE)) +
  geom_point() +
  geom_line() +
  facet_wrap(~year, scales = "free_x")
```

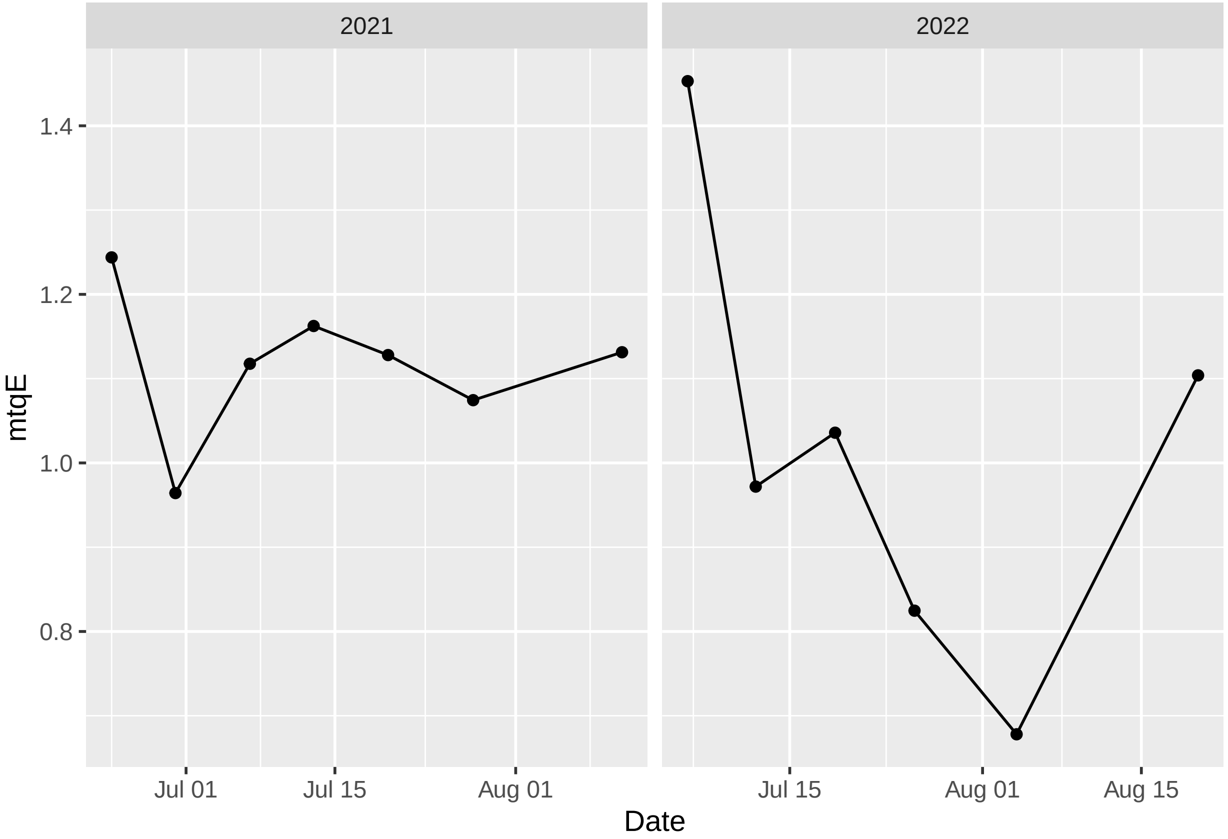

##### tqM response

```
ggplot(dat5 %>%
         group_by(Date) %>%
         summarise(mtqM = mean(mtqM), year = unique(year)),
       aes(x = Date, y = mtqM)) +
  geom_point() +
  geom_line() +
  facet_wrap(~year, scales = "free_x")
```

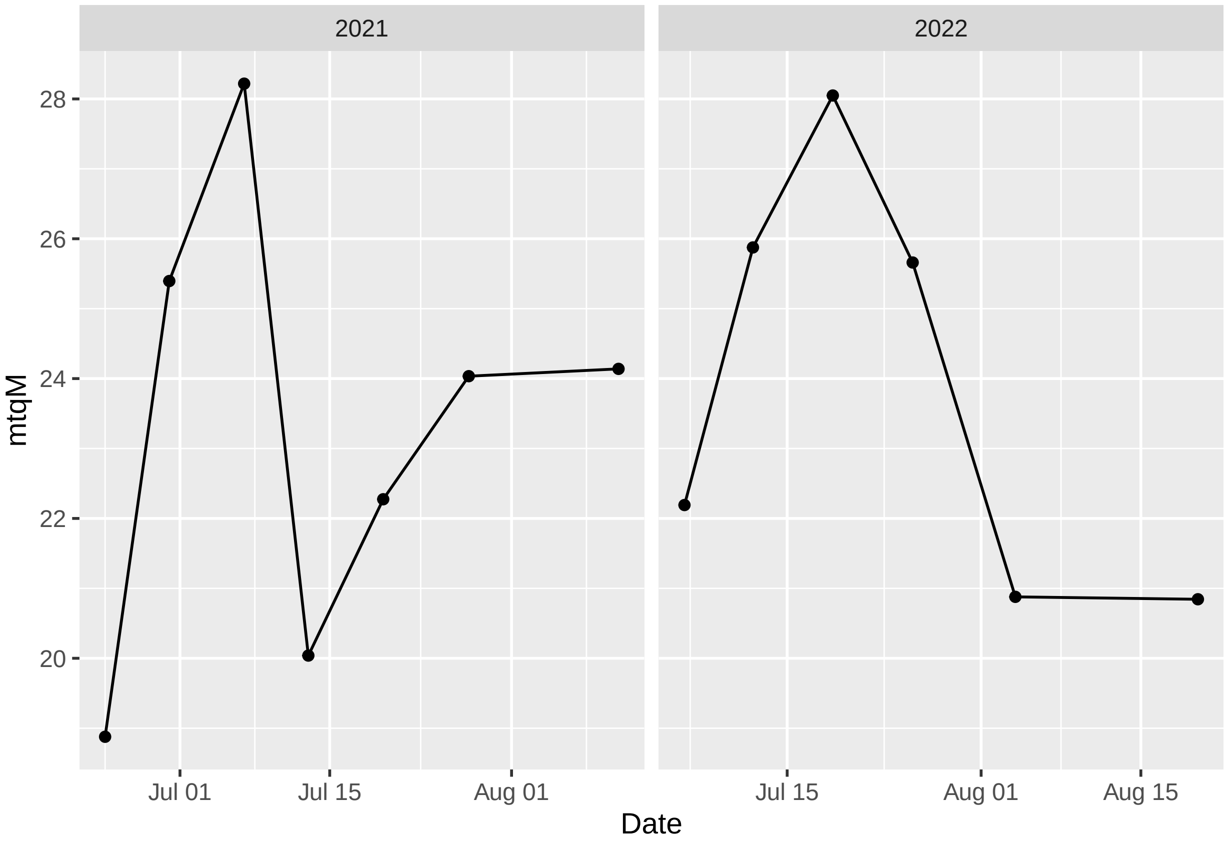

##### maxNPQ response

```
ggplot(dat5 %>%
         group_by(Date) %>%
         summarise(maxNPQ = mean(maxNPQ), year = unique(year)),
       aes(x = Date, y = maxNPQ)) +
  geom_point() +
  geom_line() +
  facet_wrap(~year, scales = "free_x")
```

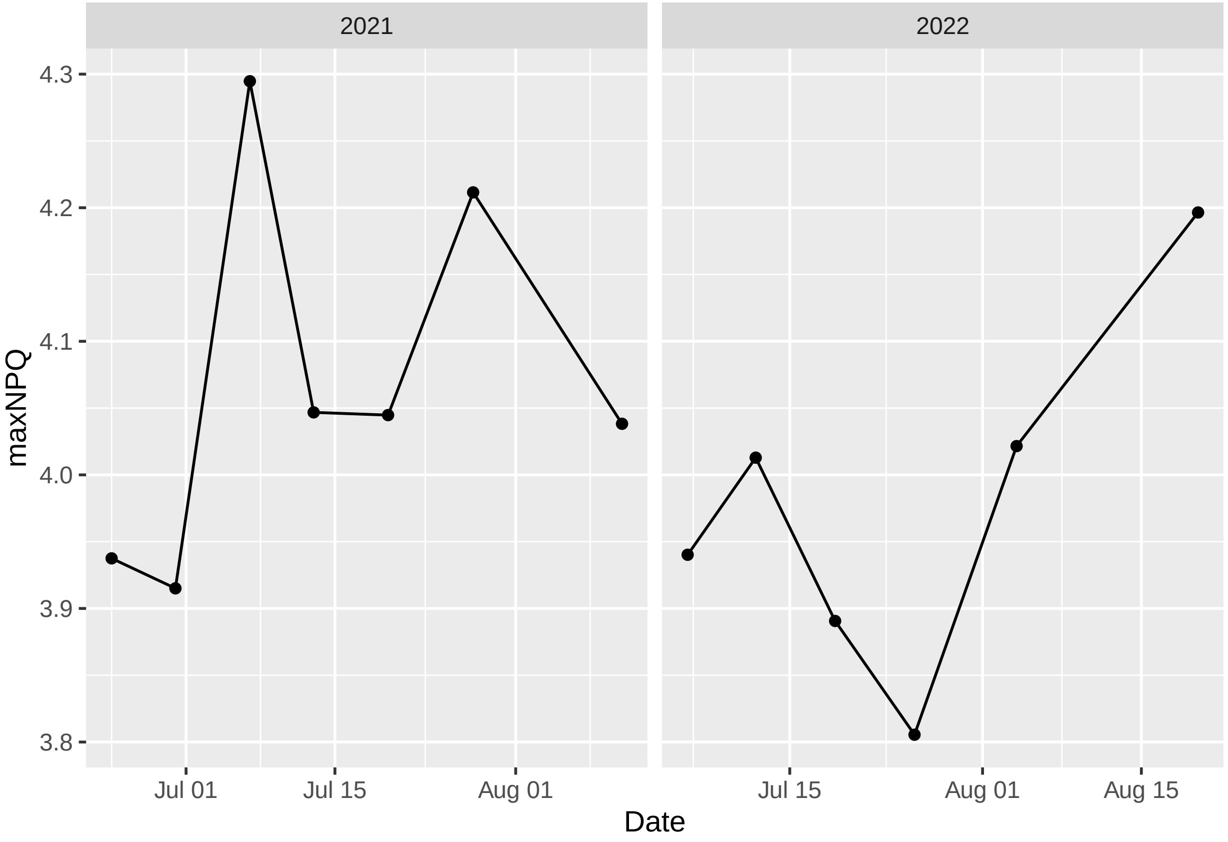

### Modeling

We fit and compare models under three circumstances:

1. model fit to all data with plot random effects
2. model fit to 2021 data with plot random effects
3. model fit to 2022 data with plot random effects

We will consider two functional forms for weather variables and date variables:

1. quadratic model in date only
2. quadratic model in date with main effect terms for additional weather variables (denoted full model)

This gives a total of six candidate models fit in a common order:

1. m1_response
2. m1_response_full
3. m2_response
4. m2_response_full
5. m3_response
6. m3_response_full

Our procedure for determining influential genotypes for each response will follow these steps:

1. Investigate model diagnostics for the six candidate models. Refit with transformations if needed.
2. Determine if each genotype provides a better fit over reference level through a leave-one-variable-out AIC procedure. We refit the model with the genotype under investigation removed and record AIC values for this model and the original model with the genotype included. The models are fit so that the genotype effect is interpreted as its difference from the RC reference level (the RC level is encoded in the model intercept).
3. Obtain coefficient estimates for the genotype under study.
4. Report genotypes that have consistent sign estimates and are determined to differentiate from the reference level across all six candidate models. These are the genotypes that we call influential.

#### AqE response

```
m1_mAqE = lmer(mAqE ~ ID + Date_num + I(Date_num^2) +
               (1|plot_number_year),
          data = dat5, REML = FALSE, control = lmerControl(optimizer ="Nelder_Mead"))
m1_mAqE_full = lmer(mAqE ~ ID + Date_num + I(Date_num^2) +
                      Ta + VPD + Fsd + Precip + Precip_cum +
                      Ta_7day + VPD_7day + Fsd_7day + Precip_7day +
                      (1|plot_number_year),
          data = dat5, REML = FALSE, control = lmerControl(optimizer ="Nelder_Mead"))
AIC(m1_mAqE)
```

```
## [1] -239.5221
```

```
AIC(m1_mAqE_full)
```

```
## [1] -513.6005
```

```
m2_mAqE = lmer(mAqE ~ ID + Date_num + I(Date_num^2) +
               (1|plot_number),
          data = dat5 %>% filter(year == 2021), REML = FALSE,
          control = lmerControl(optimizer ="Nelder_Mead"))
m2_mAqE_full = lmer(mAqE ~ ID + Date_num + I(Date_num^2) +
                      Ta + VPD + Fsd + Precip + Precip_cum +
                      Ta_7day + VPD_7day + Fsd_7day + Precip_7day +
                      (1|plot_number),
          data = dat5 %>% filter(year == 2021), REML = FALSE,
          control = lmerControl(optimizer ="Nelder_Mead"))
AIC(m2_mAqE)
```

```
## [1] 7.947379
```

```
AIC(m2_mAqE_full)
```

```
## [1] -165.8554
```

```
m3_mAqE = lmer(mAqE ~ ID + Date_num + I(Date_num^2) +
               (1|plot_number),
          data = dat5 %>% filter(year == 2022), REML = FALSE,
          control = lmerControl(optimizer ="Nelder_Mead"))
m3_mAqE_full = lmer(mAqE ~ ID + Date_num + I(Date_num^2) +
                      Ta + VPD + Fsd + Precip + Precip_cum +
                      Ta_7day + VPD_7day + Fsd_7day + Precip_7day +
                      (1|plot_number),
          data = dat5 %>% filter(year == 2022), REML = FALSE,
          control = lmerControl(optimizer ="Nelder_Mead"))
AIC(m3_mAqE)
```

```
## [1] -304.2494
```

```
AIC(m3_mAqE_full)
```

```
## [1] -350.7777
```

**Diagnostics**

Diagnostic plots for the AqE response are provided (small model followed by full model for models 1 through 3). Modeling assumptions appear to be satisfied.

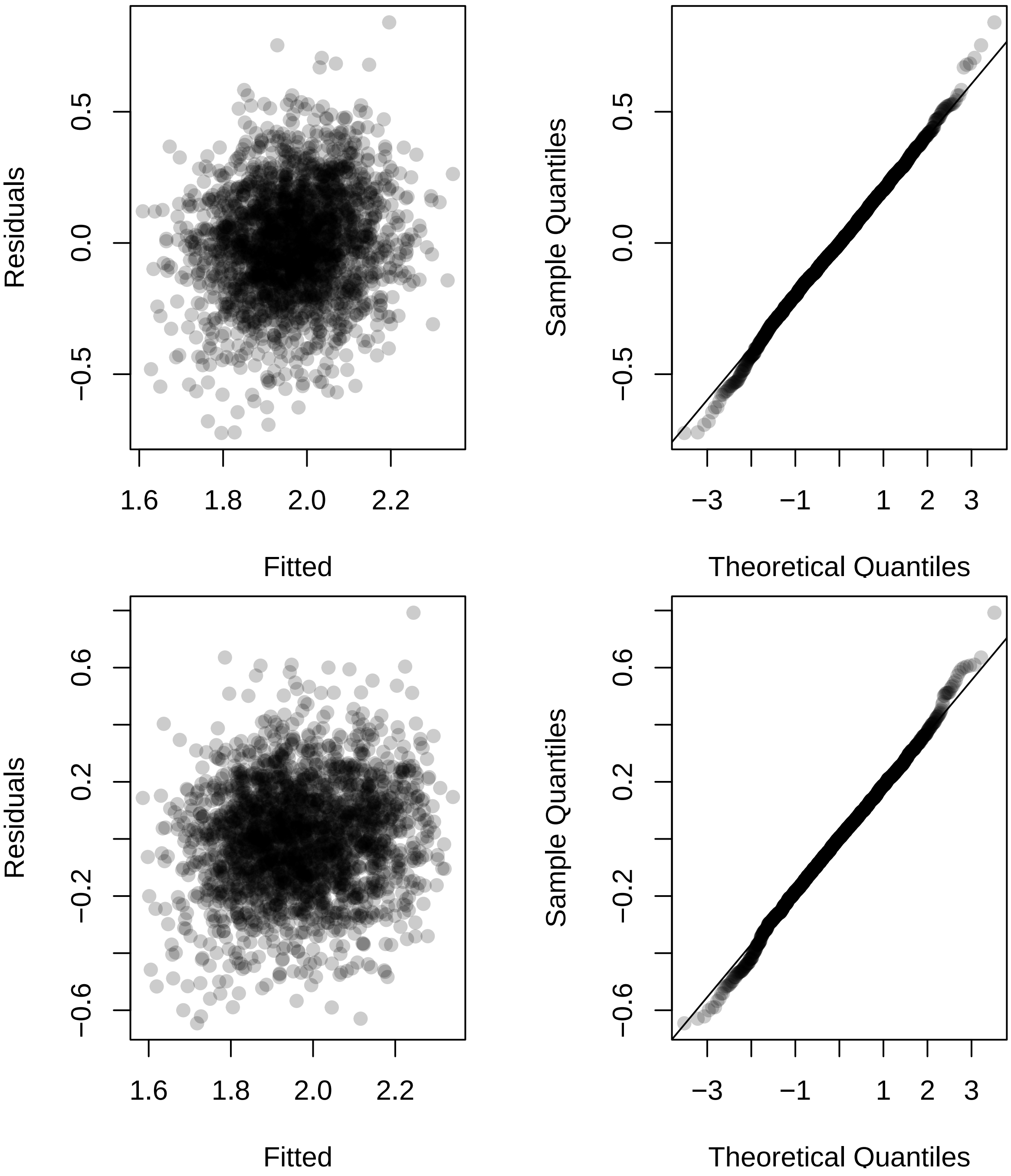

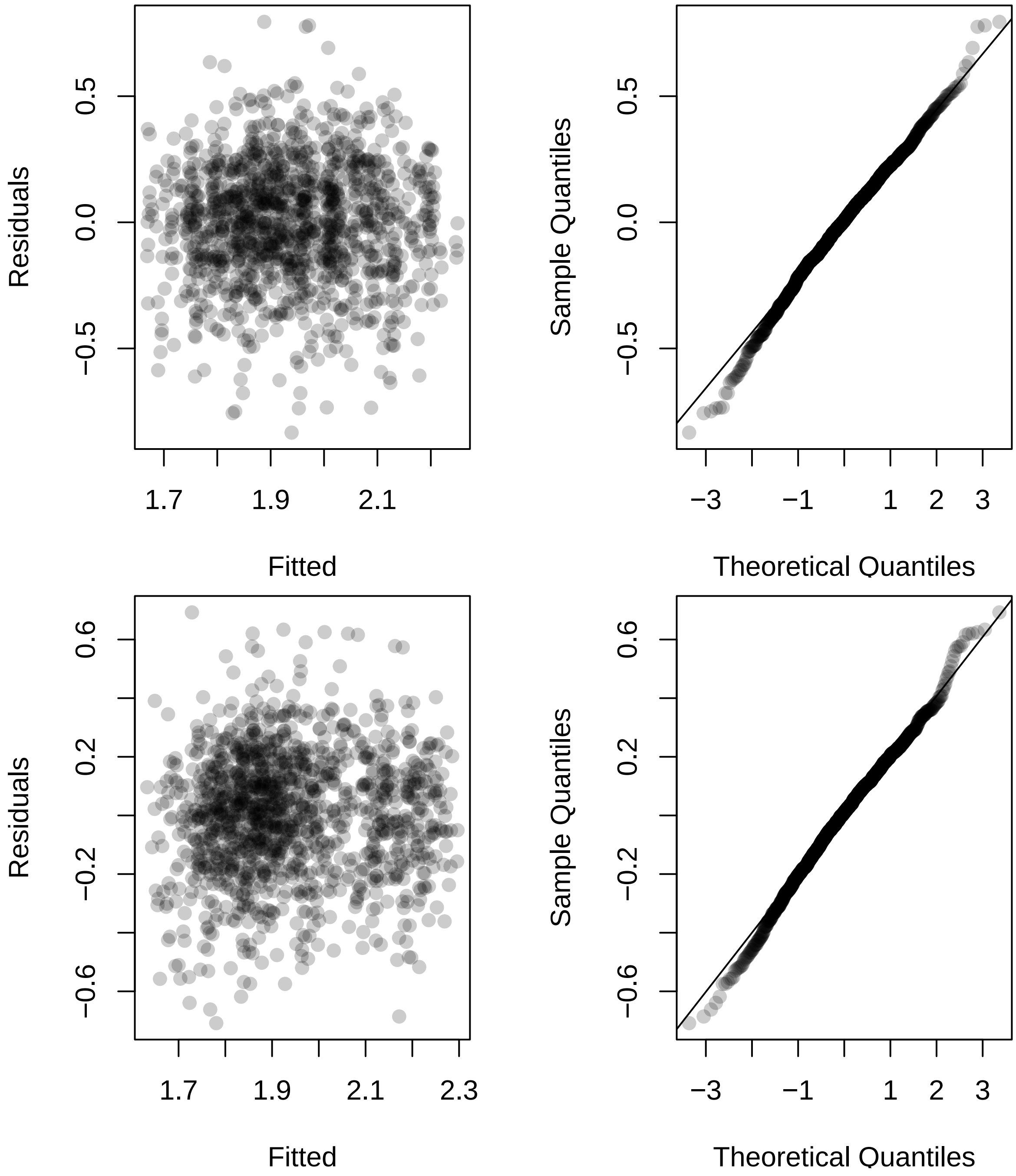
Residuals
0.5
0.0
−0.5
1.7
1.9
2.1
Fitted
Sample Quantiles
0.5
0.0
−0.5
−3
−1
1
2
3
Theoretical Quantiles
Residuals
0.6
0.2
−0.2
−0.6
1.7
1.9
2.1
2.3
Fitted
Sample Quantiles
0.6
0.2
−0.2
−0.6
−3
−1
1
2
3
Theoretical Quantiles

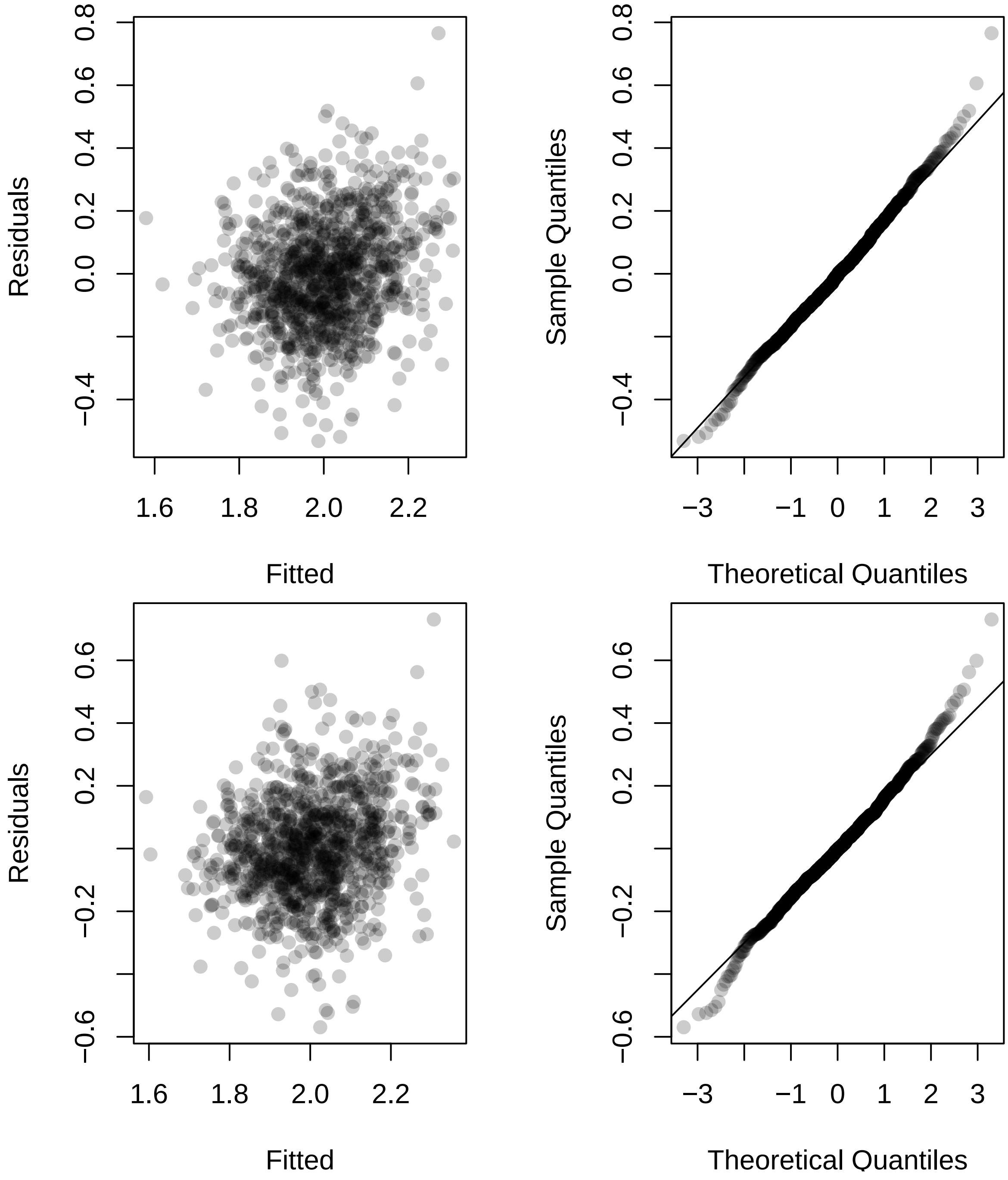
Residuals
Fitted
Sample Quantiles
Theoretical Quantiles
Residuals
Fitted
Sample Quantiles
Theoretical Quantiles

##### Investigate genotypes

We now perform an AIC based procedure to find genotypes that differ from the RC reference level.

```r
## AIC for each ID variable from full AqE fixed-effects model
M = model.matrix(mAqE ~ ID + Date_num + I(Date_num^2), data = dat5)
M_full = model.matrix(mAqE ~ ID + Date_num + I(Date_num^2) +
                      Ta + VPD + Fsd + Precip + Precip_cum +
                      Ta_7day + VPD_7day + Fsd_7day + Precip_7day,
                 data = dat5)

M2021 = model.matrix(mAqE ~ ID + Date_num + I(Date_num^2),
                 data = dat5 %>% filter(year == 2021))
M2021_full = model.matrix(mAqE ~ ID + Date_num + I(Date_num^2) +
                      Ta + VPD + Fsd + Precip + Precip_cum +
                      Ta_7day + VPD_7day + Fsd_7day + Precip_7day,
                 data = dat5 %>% filter(year == 2021))

M2022 = model.matrix(mAqE ~ ID + Date_num + I(Date_num^2),
                 data = dat5 %>% filter(year == 2022))
M2022_full = model.matrix(mAqE ~ ID + Date_num + I(Date_num^2) +
                            Ta + VPD + Fsd + Precip + Precip_cum +
                            Ta_7day + VPD_7day + Fsd_7day + Precip_7day,
                 data = dat5 %>% filter(year == 2022))

ncores = detectCores() - 2
AIC_IDs_mAqE = do.call(rbind, mclapply(
  grep("IDNA", colnames(M)), function(j){
    M1 = M[, -j]
    foo = lmer(mAqE ~ -1 + M1 + (1|plot_number_year),
                data = dat5, REML = FALSE)
    M1_full = M_full[, -j]
    foo_full = lmer(mAqE ~ -1 + M1_full + (1|plot_number_year),
                data = dat5, REML = FALSE)
    M12021 = M2021[, -j]
    bar = lmer(mAqE ~ -1 + M12021 + (1|plot_number),
                data = dat5 %>% filter(year == 2021), REML = FALSE)
    M12021_full = M2021_full[, -j]
    bar_full = lmer(mAqE ~ -1 + M12021_full + (1|plot_number),
                data = dat5 %>% filter(year == 2021), REML = FALSE)
    M12022 = M2022[, -j]
    baz = lmer(mAqE ~ -1 + M12022 + (1|plot_number),
                data = dat5 %>% filter(year == 2022), REML = FALSE)
    M12022_full = M2022_full[, -j]
    baz_full = lmer(mAqE ~ -1 + M12022_full + (1|plot_number),
                data = dat5 %>% filter(year == 2022), REML = FALSE)

    c(AIC(m1_mAqE) - AIC(foo),
      AIC(m1_mAqE_full) - AIC(foo_full),
      AIC(m2_mAqE) - AIC(bar),
      AIC(m2_mAqE_full) - AIC(bar_full),
      AIC(m3_mAqE) - AIC(baz),
      AIC(m3_mAqE_full) - AIC(baz_full))
  }, mc.cores = ncores))
```

```r
rownames(AIC_IDs_mAqE) = colnames(M)[grep("IDNA", colnames(M))]
```

Negative values indicate that the genotype in question provides better model fit when differentiated from the RC reference level.

```r
round(AIC_IDs_mAqE, 3)
```

```
##            [,1]   [,2]   [,3]    [,4]   [,5]   [,6]
## IDNAM10 -2.298 -1.826 -2.173  -2.569  1.390  1.501
## IDNAM11 -0.713 -0.866 -1.952  -2.958  1.845  1.885
## IDNAM12  1.931  1.914  1.499   0.696  1.716  1.651
## IDNAM13  1.498  1.486 -1.282  -1.999  1.441  1.407
## IDNAM14 -2.445 -2.428 -4.651  -5.191  1.561  1.698
## IDNAM15 -1.288 -1.259 -5.021  -6.216  1.998  2.000
## IDNAM17  1.690  1.673  2.016   1.388 -0.293 -0.237
## IDNAM18  1.950  1.947  2.525   1.931  2.000  1.984
## IDNAM2   1.619  1.583  1.181   0.462  1.871  1.895
## IDNAM22  1.492  1.245  0.496  -0.314  1.980  1.975
## IDNAM23  1.956  1.891  1.730   0.773  1.782  1.771
## IDNAM24 -8.752 -8.908 -9.907 -10.626 -0.065 -0.009
## IDNAM25  1.428  1.627  0.523   0.318  1.889  1.890
## IDNAM26 -3.482 -3.312 -4.456  -4.706  1.271  1.196
## IDNAM27  1.983  1.965  2.017   1.266  1.770  1.764
## IDNAM28 -0.732 -1.040  2.348   1.710 -1.603 -1.730
## IDNAM29  0.181  0.413 -0.917  -1.135  1.957  1.962
## IDNAM3  -2.038 -2.061 -4.648  -4.974  1.801  1.747
## IDNAM30  0.734  0.713 -1.868  -2.528  1.919  1.896
## IDNAM31 -4.187 -3.909 -4.408  -5.431  0.968  1.014
## IDNAM32  1.560  1.597  1.948   1.409  1.980  1.980
## IDNAM33  1.594  1.463  2.109   1.044  1.999  1.986
## IDNAM34  1.993  1.957  2.605   1.977  1.850  1.864
## IDNAM36  1.211  1.193  1.827   1.191  1.815  1.821
## IDNAM37  1.394  1.294  2.584   1.975  1.023  0.820
## IDNAM38  1.546  1.608  2.566   1.930  0.751  0.756
## IDNAM39  1.939  1.954  0.755   0.172  1.347  1.275
## IDNAM4  -3.070 -2.860  1.631   0.847 -2.902 -2.864
## IDNAM40 -1.669 -1.988 -3.050  -3.708  1.576  1.588
## IDNAM41 -1.555 -1.395 -3.221  -4.141  1.823  1.873
## IDNAM42 -4.047 -4.385 -4.515  -5.464  0.574  0.649
## IDNAM46  1.887  1.888  1.274   0.687  1.699  1.716
## IDNAM48  0.416  0.459  1.640   0.719  1.474  1.494
## IDNAM5   1.507  1.630  1.137   0.697  1.971  1.960
## IDNAM50  1.140  1.133  2.050   1.634  1.619  1.563
## IDNAM54 -3.226 -2.787 -5.249  -5.237  1.568  1.649
## IDNAM6  -0.799 -1.100 -6.588  -7.802  1.926  1.903
## IDNAM64 -1.564 -1.388 -4.064  -4.067  1.882  1.891
## IDNAM8   1.134  1.028  2.358   1.704  1.232  1.174
## IDNAM9   1.782  1.815  0.010  -0.463  1.601  1.537
```

```r
sign(AIC_IDs_mAqE < 0)
```

```
##         [,1] [,2] [,3] [,4] [,5] [,6]
## IDNAM10    1    1    1    1    0    0
## IDNAM11    1    1    1    1    0    0
## IDNAM12    0    0    0    0    0    0
```

```
## IDNAM13    0    0    1    1    0    0
## IDNAM14    1    1    1    1    0    0
## IDNAM15    1    1    1    1    0    0
## IDNAM17    0    0    0    0    1    1
## IDNAM18    0    0    0    0    0    0
## IDNAM2     0    0    0    0    0    0
## IDNAM22    0    0    0    1    0    0
## IDNAM23    0    0    0    0    0    0
## IDNAM24    1    1    1    1    1    1
## IDNAM25    0    0    0    0    0    0
## IDNAM26    1    1    1    1    0    0
## IDNAM27    0    0    0    0    0    0
## IDNAM28    1    1    0    0    1    1
## IDNAM29    0    0    1    1    0    0
## IDNAM3     1    1    1    1    0    0
## IDNAM30    0    0    1    1    0    0
## IDNAM31    1    1    1    1    0    0
## IDNAM32    0    0    0    0    0    0
## IDNAM33    0    0    0    0    0    0
## IDNAM34    0    0    0    0    0    0
## IDNAM36    0    0    0    0    0    0
## IDNAM37    0    0    0    0    0    0
## IDNAM38    0    0    0    0    0    0
## IDNAM39    0    0    0    0    0    0
## IDNAM4     1    1    0    0    1    1
## IDNAM40    1    1    1    1    0    0
## IDNAM41    1    1    1    1    0    0
## IDNAM42    1    1    1    1    0    0
## IDNAM46    0    0    0    0    0    0
## IDNAM48    0    0    0    0    0    0
## IDNAM5     0    0    0    0    0    0
## IDNAM50    0    0    0    0    0    0
## IDNAM54    1    1    1    1    0    0
## IDNAM6     1    1    1    1    0    0
## IDNAM64    1    1    1    1    0    0
## IDNAM8     0    0    0    0    0    0
## IDNAM9     0    0    0    1    0    0
```

We now investigate the specific coefficient effects from our fitted models. Specific interest is in determining the sign of the effect. Here are the effect estimates from each fitted model:

```
coefs_mAqE = round(cbind(summary(m1_mAqE)$coefficients[2:41, 1],
                   summary(m1_mAqE_full)$coefficients[2:41, 1],
                   summary(m2_mAqE)$coefficients[2:41, 1],
                   summary(m2_mAqE_full)$coefficients[2:41, 1],
                   summary(m3_mAqE)$coefficients[2:41, 1],
                   summary(m3_mAqE_full)$coefficients[2:41, 1]), 3)
coefs_mAqE
```

```
##            [,1]   [,2]   [,3]   [,4]   [,5]   [,6]
## IDNAM10   0.110  0.103  0.136  0.132  0.064  0.058
## IDNAM11   0.086  0.087  0.130  0.135  0.032  0.028
## IDNAM12   0.014  0.015  0.065  0.069 -0.044 -0.048
## IDNAM13   0.037  0.037  0.121  0.123 -0.060 -0.062
## IDNAM14   0.110  0.109  0.164  0.163  0.053  0.044
## IDNAM15   0.094  0.092  0.169  0.175  0.003  0.000
## IDNAM17  -0.029 -0.030  0.047  0.048 -0.123 -0.122
## IDNAM18  -0.012 -0.012 -0.017 -0.016  0.002 -0.010
## IDNAM2    0.032  0.033  0.071  0.075 -0.028 -0.025
## IDNAM22   0.038  0.045  0.090  0.094 -0.012 -0.013
## IDNAM23   0.011  0.017  0.057  0.067 -0.038 -0.039
## IDNAM24   0.171  0.170  0.217  0.217  0.116  0.114
## IDNAM25   0.040  0.032  0.088  0.079 -0.027 -0.027
## IDNAM26   0.123  0.119  0.166  0.160  0.068  0.071
## IDNAM27   0.007  0.010  0.046  0.052 -0.040 -0.040
## IDNAM28  -0.086 -0.090 -0.031 -0.032 -0.156 -0.158
## IDNAM29   0.071  0.065  0.113  0.107  0.017  0.016
## IDNAM3    0.105  0.104  0.169  0.164  0.036  0.040
## IDNAM30   0.059  0.058  0.128  0.128 -0.023 -0.026
## IDNAM31   0.131  0.127  0.163  0.167  0.083  0.081
## IDNAM32   0.034  0.032  0.048  0.046  0.011  0.011
## IDNAM33   0.037  0.042  0.054  0.075 -0.002 -0.010
## IDNAM34   0.005  0.011 -0.003  0.009  0.034  0.033
## IDNAM36   0.046  0.046  0.052  0.054  0.035  0.034
## IDNAM37  -0.041 -0.043 -0.009 -0.010 -0.081 -0.089
## IDNAM38  -0.035 -0.032  0.013  0.016 -0.091 -0.090
## IDNAM39   0.013  0.011  0.083  0.082 -0.066 -0.069
## IDNAM4    0.119  0.115  0.060  0.065  0.184  0.183
## IDNAM40   0.100  0.103  0.145  0.145  0.052  0.052
## IDNAM41   0.099  0.096  0.147  0.150  0.034  0.029
## IDNAM42   0.129  0.130  0.164  0.167  0.096  0.093
## IDNAM46   0.017  0.017  0.069  0.069 -0.045 -0.043
## IDNAM48   0.066  0.064  0.059  0.069  0.058  0.057
## IDNAM5    0.037  0.032  0.072  0.069 -0.014 -0.016
## IDNAM50   0.048  0.048  0.046  0.037  0.049  0.052
## IDNAM54   0.119  0.113  0.172  0.164  0.053  0.047
## IDNAM6    0.087  0.091  0.186  0.191 -0.022 -0.025
## IDNAM64   0.098  0.095  0.159  0.151  0.028  0.027
## IDNAM8   -0.049 -0.051 -0.031 -0.033 -0.072 -0.074
## IDNAM9    0.025  0.023  0.100  0.097 -0.053 -0.057
```

Here are the corresponding signs:

```r
sign(coefs_mAqE > 0)
```

```
##          [,1] [,2] [,3] [,4] [,5] [,6]
## IDNAM10     1    1    1    1    1    1
## IDNAM11     1    1    1    1    1    1
## IDNAM12     1    1    1    1    0    0
## IDNAM13     1    1    1    1    0    0
## IDNAM14     1    1    1    1    1    1
## IDNAM15     1    1    1    1    1    0
## IDNAM17     0    0    1    1    0    0
## IDNAM18     0    0    0    0    1    0
## IDNAM2      1    1    1    1    0    0
## IDNAM22     1    1    1    1    0    0
## IDNAM23     1    1    1    1    0    0
## IDNAM24     1    1    1    1    1    1
## IDNAM25     1    1    1    1    0    0
## IDNAM26     1    1    1    1    1    1
## IDNAM27     1    1    1    1    0    0
## IDNAM28     0    0    0    0    0    0
## IDNAM29     1    1    1    1    1    1
## IDNAM3      1    1    1    1    1    1
## IDNAM30     1    1    1    1    0    0
## IDNAM31     1    1    1    1    1    1
## IDNAM32     1    1    1    1    1    1
## IDNAM33     1    1    1    1    0    0
## IDNAM34     1    1    0    1    1    1
## IDNAM36     1    1    1    1    1    1
## IDNAM37     0    0    0    0    0    0
## IDNAM38     0    0    1    1    0    0
## IDNAM39     1    1    1    1    0    0
## IDNAM4      1    1    1    1    1    1
## IDNAM40     1    1    1    1    1    1
## IDNAM41     1    1    1    1    1    1
## IDNAM42     1    1    1    1    1    1
## IDNAM46     1    1    1    1    0    0
## IDNAM48     1    1    1    1    1    1
## IDNAM5      1    1    1    1    0    0
## IDNAM50     1    1    1    1    1    1
## IDNAM54     1    1    1    1    1    1
## IDNAM6      1    1    1    1    0    0
## IDNAM64     1    1    1    1    1    1
## IDNAM8      0    0    0    0    0    0
## IDNAM9      1    1    1    1    0    0
```

Below are signs that indicate which genotypes yield better model fit when their effect is differentiated from the RC reference level. A 1 indicates a positive effect, a -1 indicates a negative effect, and a 0 indicates that the genotype was not determined to provide better model when differentiated from the RC reference level.

```
inference_mAqE = sign(coefs_mAqE) * sign(AIC_IDs_mAqE < 0)
inference_mAqE
```

```
##          [,1] [,2] [,3] [,4] [,5] [,6]
## IDNAM10     1    1    1    1    0    0
## IDNAM11     1    1    1    1    0    0
## IDNAM12     0    0    0    0    0    0
## IDNAM13     0    0    1    1    0    0
## IDNAM14     1    1    1    1    0    0
## IDNAM15     1    1    1    1    0    0
## IDNAM17     0    0    0    0   -1   -1
## IDNAM18     0    0    0    0    0    0
## IDNAM2      0    0    0    0    0    0
## IDNAM22     0    0    0    1    0    0
## IDNAM23     0    0    0    0    0    0
## IDNAM24     1    1    1    1    1    1
## IDNAM25     0    0    0    0    0    0
## IDNAM26     1    1    1    1    0    0
## IDNAM27     0    0    0    0    0    0
## IDNAM28    -1   -1    0    0   -1   -1
## IDNAM29     0    0    1    1    0    0
## IDNAM3      1    1    1    1    0    0
## IDNAM30     0    0    1    1    0    0
## IDNAM31     1    1    1    1    0    0
## IDNAM32     0    0    0    0    0    0
## IDNAM33     0    0    0    0    0    0
## IDNAM34     0    0    0    0    0    0
## IDNAM36     0    0    0    0    0    0
## IDNAM37     0    0    0    0    0    0
## IDNAM38     0    0    0    0    0    0
## IDNAM39     0    0    0    0    0    0
## IDNAM4      1    1    0    0    1    1
## IDNAM40     1    1    1    1    0    0
## IDNAM41     1    1    1    1    0    0
## IDNAM42     1    1    1    1    0    0
## IDNAM46     0    0    0    0    0    0
## IDNAM48     0    0    0    0    0    0
## IDNAM5      0    0    0    0    0    0
## IDNAM50     0    0    0    0    0    0
## IDNAM54     1    1    1    1    0    0
## IDNAM6      1    1    1    1    0    0
## IDNAM64     1    1    1    1    0    0
## IDNAM8      0    0    0    0    0    0
## IDNAM9      0    0    0    1    0    0
```

##### Results

Here are the effect size estimates for genotypes that were to determined to differentiate from the RC reference level in all considered models and had the same estimated sign across all considered models.

```
rownames(coefs_mAqE)[abs(rowSums(inference_mAqE)) == 6]
```

```
## [1] "IDNAM24"
```

```
coefs_mAqE[abs(rowSums(inference_mAqE)) == 6, ]
```

```
## [1] 0.171 0.170 0.217 0.217 0.116 0.114
```

##### AqI response

```r
m1_mAqI = lmer(mAqI ~ ID + Date_num + I(Date_num^2) +
               (1|plot_number_year),
          data = dat5, REML = FALSE, control = lmerControl(optimizer ="Nelder_Mead"))
m1_mAqI_full = lmer(mAqI ~ ID + Date_num + I(Date_num^2) +
                      Ta + VPD + Fsd + Precip + Precip_cum +
                      Ta_7day + VPD_7day + Fsd_7day + Precip_7day +
                      (1|plot_number_year),
          data = dat5, REML = FALSE, control = lmerControl(optimizer ="Nelder_Mead"))
AIC(m1_mAqI)
```

```
## [1] 12.24326
```

```r
AIC(m1_mAqI_full)
```

```
## [1] -1033.198
```

```r
m2_mAqI = lmer(mAqI ~ ID + Date_num + I(Date_num^2) +
               (1|plot_number),
          data = dat5 %>% filter(year == 2021), REML = FALSE,
          control = lmerControl(optimizer ="Nelder_Mead"))
m2_mAqI_full = lmer(mAqI ~ ID + Date_num + I(Date_num^2) +
                      Ta + VPD + Fsd + Precip + Precip_cum +
                      Ta_7day + VPD_7day + Fsd_7day + Precip_7day +
                      (1|plot_number),
          data = dat5 %>% filter(year == 2021), REML = FALSE,
          control = lmerControl(optimizer ="Nelder_Mead"))
AIC(m2_mAqI)
```

```
## [1] -1105.714
```

```r
AIC(m2_mAqI_full)
```

```
## [1] -1260.503
```

```r
m3_mAqI = lmer(mAqI ~ ID + Date_num + I(Date_num^2) +
               (1|plot_number),
          data = dat5 %>% filter(year == 2022), REML = FALSE,
          control = lmerControl(optimizer ="Nelder_Mead"))
m3_mAqI_full = lmer(mAqI ~ ID + Date_num + I(Date_num^2) +
                      Ta + VPD + Fsd + Precip + Precip_cum +
                      Ta_7day + VPD_7day + Fsd_7day + Precip_7day +
                      (1|plot_number),
          data = dat5 %>% filter(year == 2022), REML = FALSE,
          control = lmerControl(optimizer ="Nelder_Mead"))
AIC(m3_mAqI)
```

```
## [1] -1.631758
```

```r
AIC(m3_mAqI_full)
```

```
## [1] -84.55435
```

**Diagnostics**

Diagnostic plots for the AqI response are provided (small model followed by full model for models 1 through 3). Modeling assumptions appear to be violated.

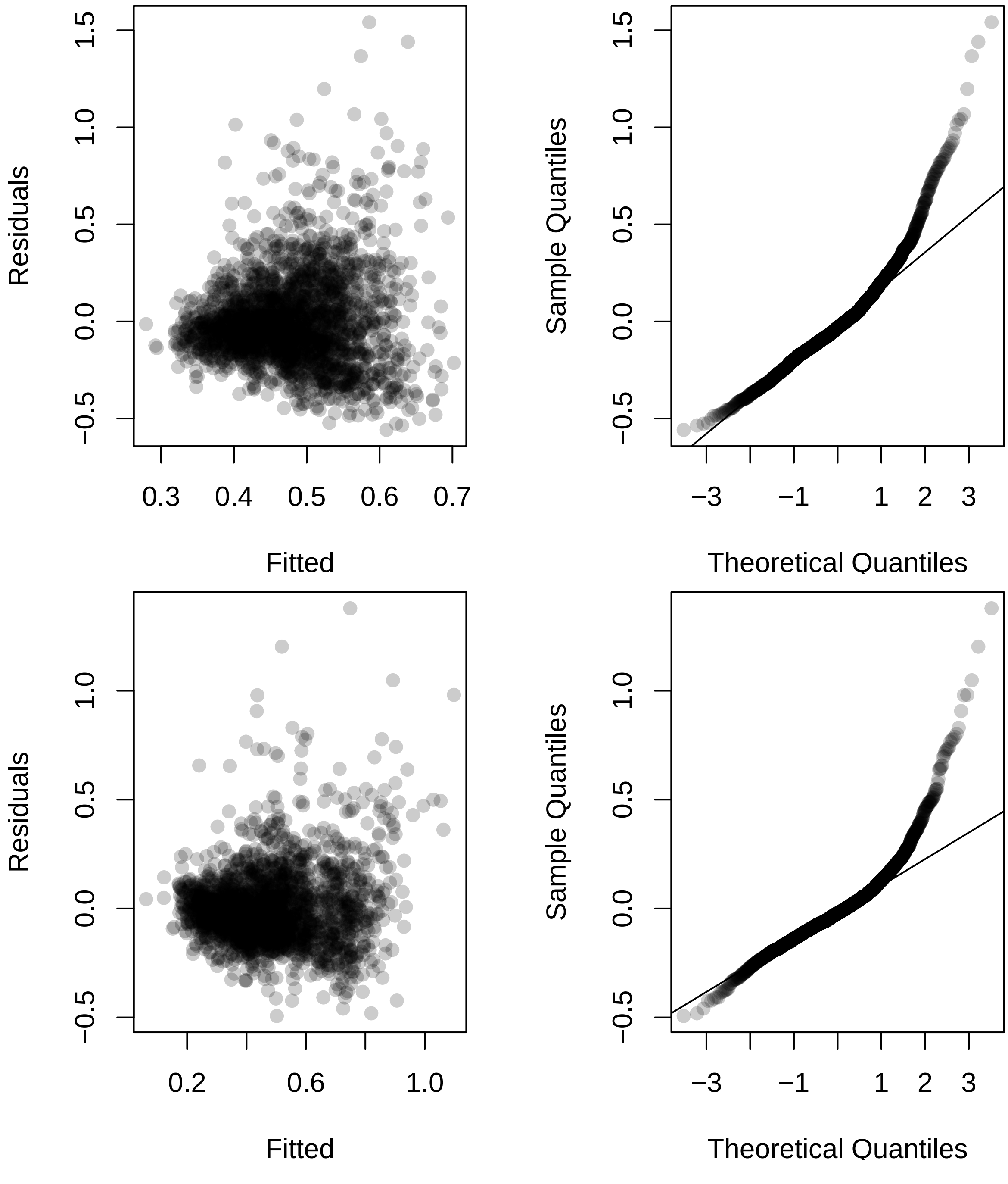

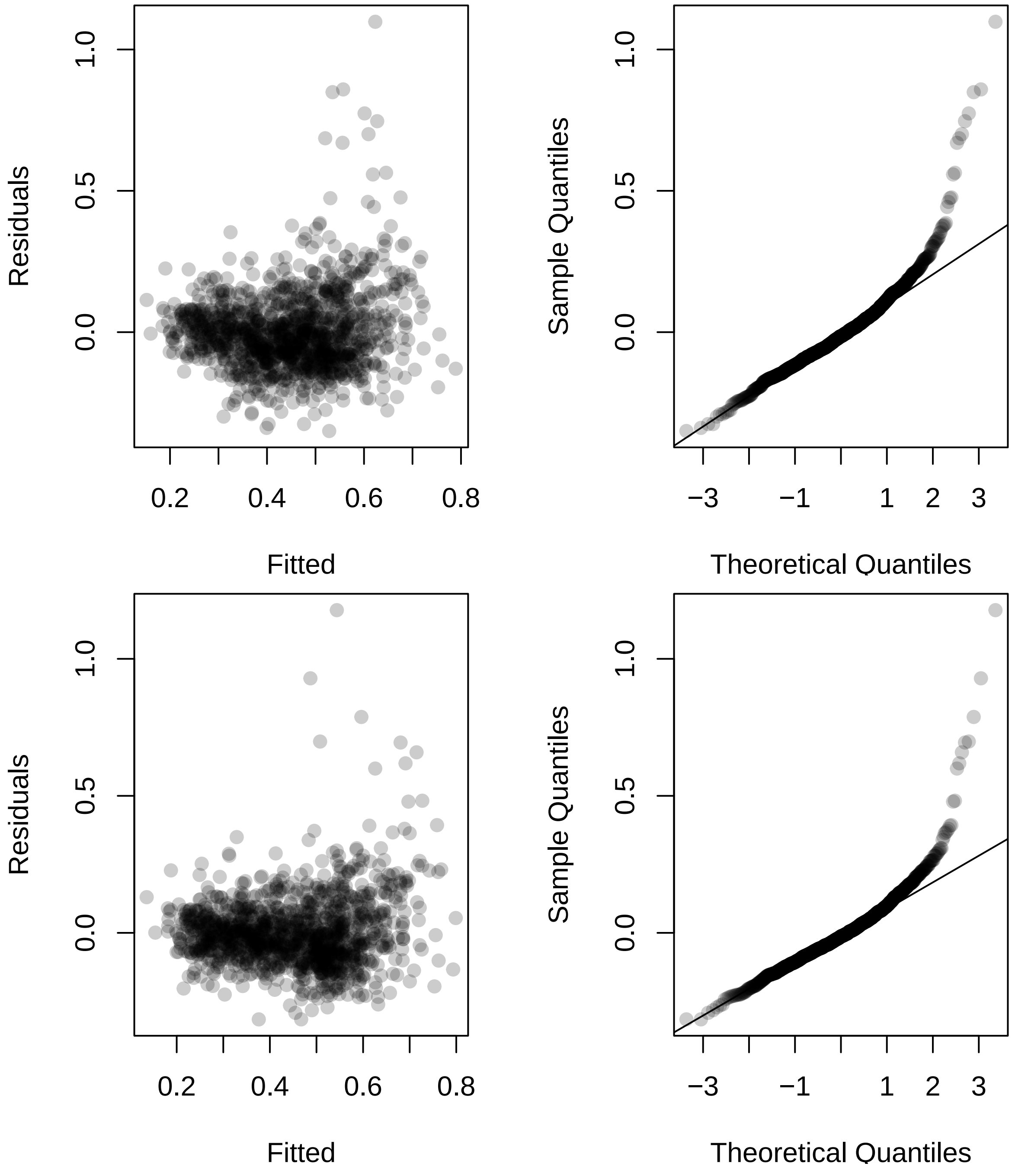
Residuals
Fitted
Sample Quantiles
Theoretical Quantiles
0.0
0.5
1.0
0.2
0.4
0.6
0.8
−3
−1
1
2
3

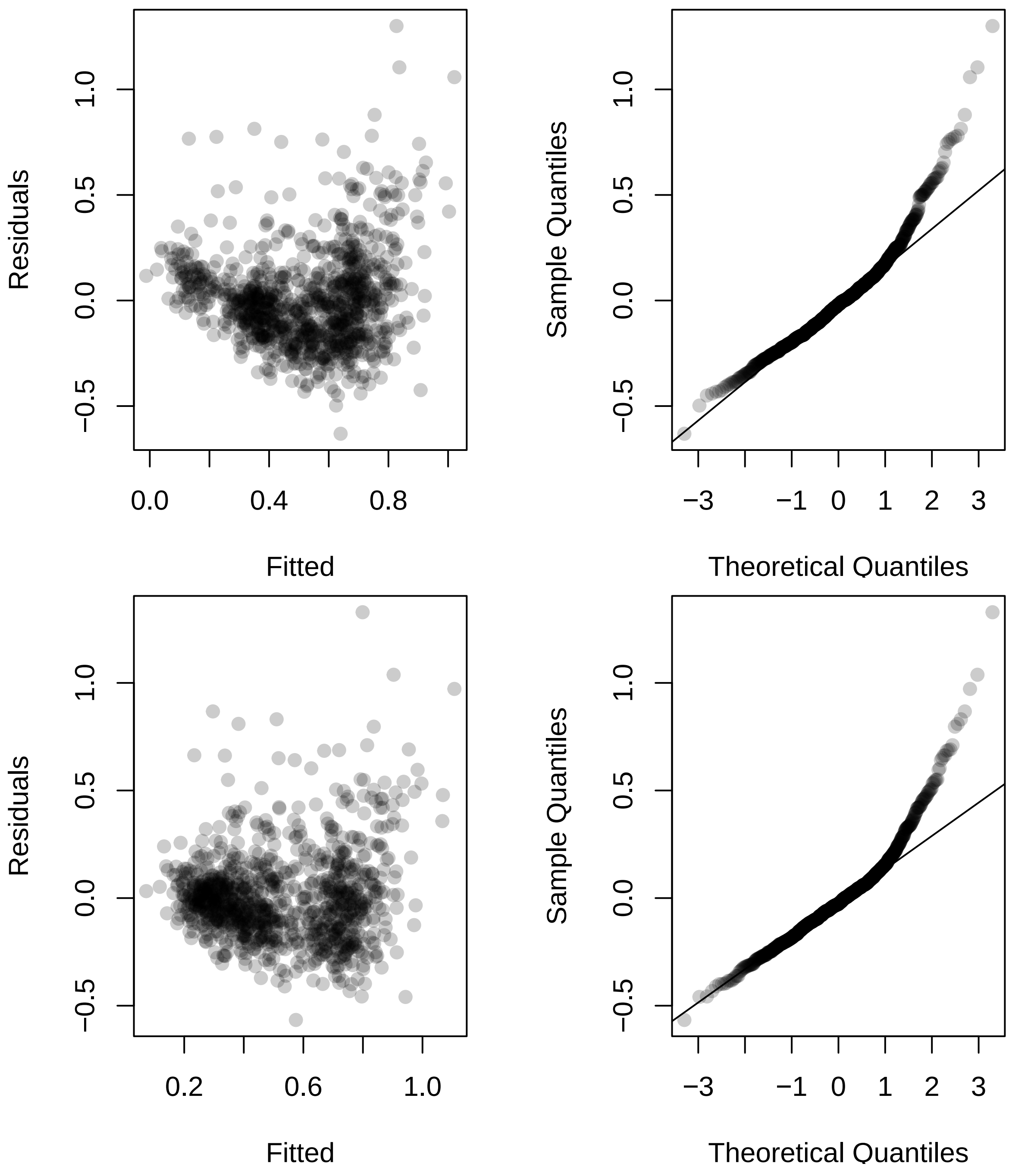
Residuals
Fitted
Sample Quantiles
Theoretical Quantiles
Residuals
Fitted
Sample Quantiles
Theoretical Quantiles

##### Cube root transform

We now consider a cube root response transformation to alleviate modeling problems. This transformation seems to alleviate serious problems with minor modeling assumption deviations remaining. Schielzeth et al (2020) notes that:

> Model estimates were usually robust to violations of assumptions, with the exception of slight upward biases in estimates of random effect variance if the generating distribution was bimodal but was modelled by Gaussian error distributions. Further, estimates for (random effect) components that violated distributional assumptions became less precise but remained unbiased. However, this particular problem did not affect other parameters of the model. The same pattern was found for strongly correlated fixed effects, which led to imprecise, but unbiased estimates, with uncertainty estimates reflecting imprecision. Overall, our results show remarkable robustness of mixed-effects models that should allow researchers to use mixed-effects models even if the distributional assumptions are objectively violated. However, this does not free researchers from careful evaluation of the model.

Here is the reference:

> Holger Schielzeth, Niels J Dingemanse, Shinichi Nakagawa, David F Westneat, Hassen Allegue, Céline Teplitsky, Denis Réale, Ned A Dochtermann, László Zsolt Garamszegi, and Yimen G Araya-Ajoy. Robustness of linear mixed-effects models to violations of distributional assumptions. Methods in Ecology and Evolution, 11(9):1141–1152, 2020.

```
m1_mAqI_cube = lmer(mAqI^(1/3) ~ ID + Date_num + I(Date_num^2) +
               (1|plot_number_year),
          data = dat5, REML = FALSE, control = lmerControl(optimizer ="Nelder_Mead"))
m1_mAqI_cube_full = lmer(mAqI^(1/3) ~ ID + Date_num + I(Date_num^2) +
                      Ta + VPD + Fsd + Precip + Precip_cum +
                      Ta_7day + VPD_7day + Fsd_7day + Precip_7day +
                      (1|plot_number_year),
          data = dat5, REML = FALSE, control = lmerControl(optimizer ="Nelder_Mead"))
AIC(m1_mAqI_cube)
```

```
## [1] -3030.211
```

```
AIC(m1_mAqI_cube_full)
```

```
## [1] -4250.185
```

```
m2_mAqI_cube = lmer(mAqI^(1/3) ~ ID + Date_num + I(Date_num^2) +
               (1|plot_number),
          data = dat5 %>% filter(year == 2021), REML = FALSE,
          control = lmerControl(optimizer ="Nelder_Mead"))
m2_mAqI_cube_full = lmer(mAqI^(1/3) ~ ID + Date_num + I(Date_num^2) +
                      Ta + VPD + Fsd + Precip + Precip_cum +
                      Ta_7day + VPD_7day + Fsd_7day + Precip_7day +
                      (1|plot_number),
          data = dat5 %>% filter(year == 2021), REML = FALSE,
          control = lmerControl(optimizer ="Nelder_Mead"))
AIC(m2_mAqI_cube)
```

```
## [1] -2714.663
```

```
AIC(m2_mAqI_cube_full)
```

```
## [1] -2934.412
```

```r
m3_mAqI_cube = lmer(mAqI^(1/3) ~ ID + Date_num + I(Date_num^2) +
               (1|plot_number),
           data = dat5 %>% filter(year == 2022), REML = FALSE,
           control = lmerControl(optimizer ="Nelder_Mead"))
m3_mAqI_cube_full = lmer(mAqI^(1/3) ~ ID + Date_num + I(Date_num^2) +
                      Ta + VPD + Fsd + Precip + Precip_cum +
                      Ta_7day + VPD_7day + Fsd_7day + Precip_7day +
                      (1|plot_number),
           data = dat5 %>% filter(year == 2022), REML = FALSE,
           control = lmerControl(optimizer ="Nelder_Mead"))
AIC(m3_mAqI_cube)
```

```
## [1] -1485.108
```

```r
AIC(m3_mAqI_cube_full)
```

```
## [1] -1562.261
```

**Diagnostics**

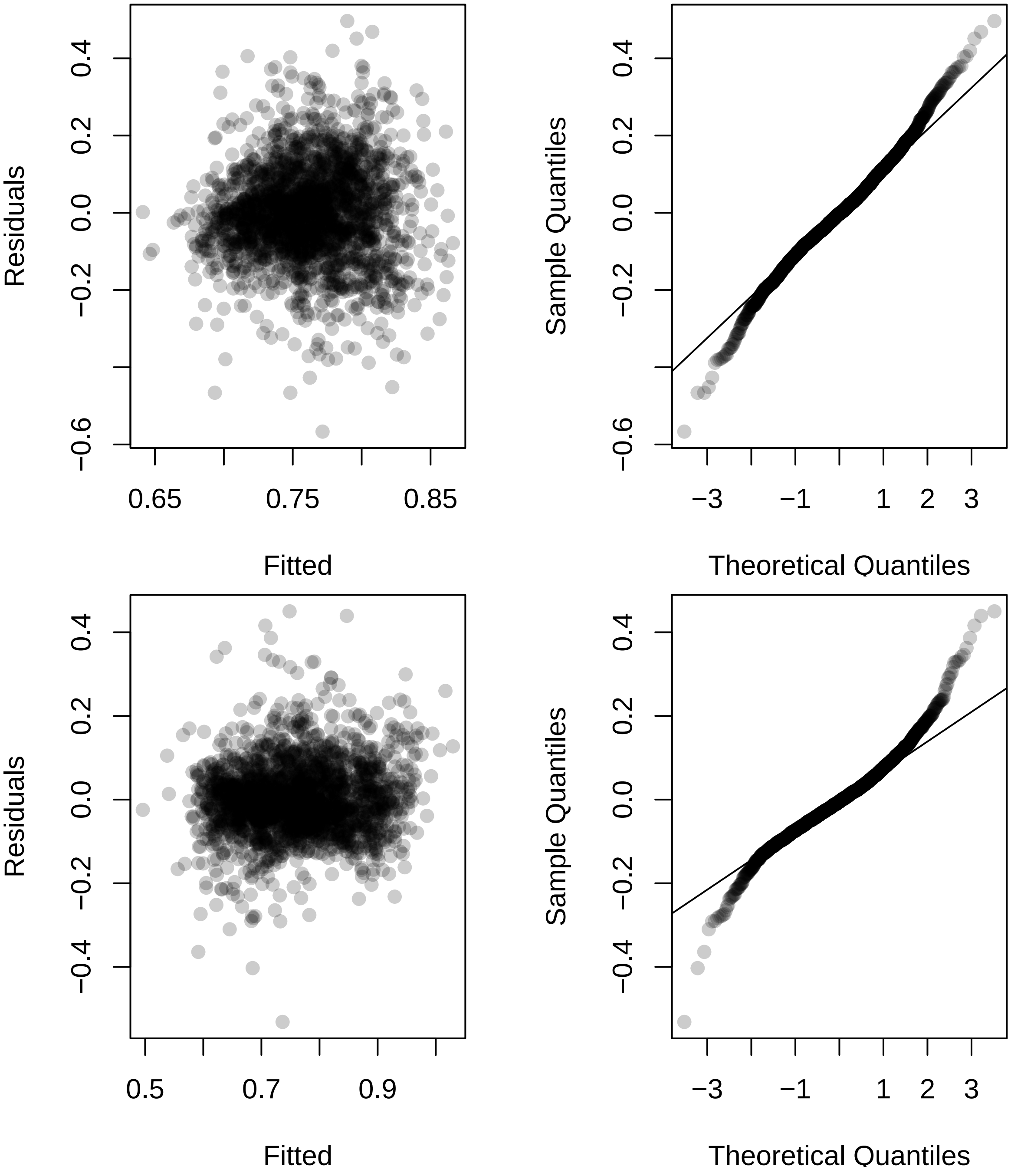

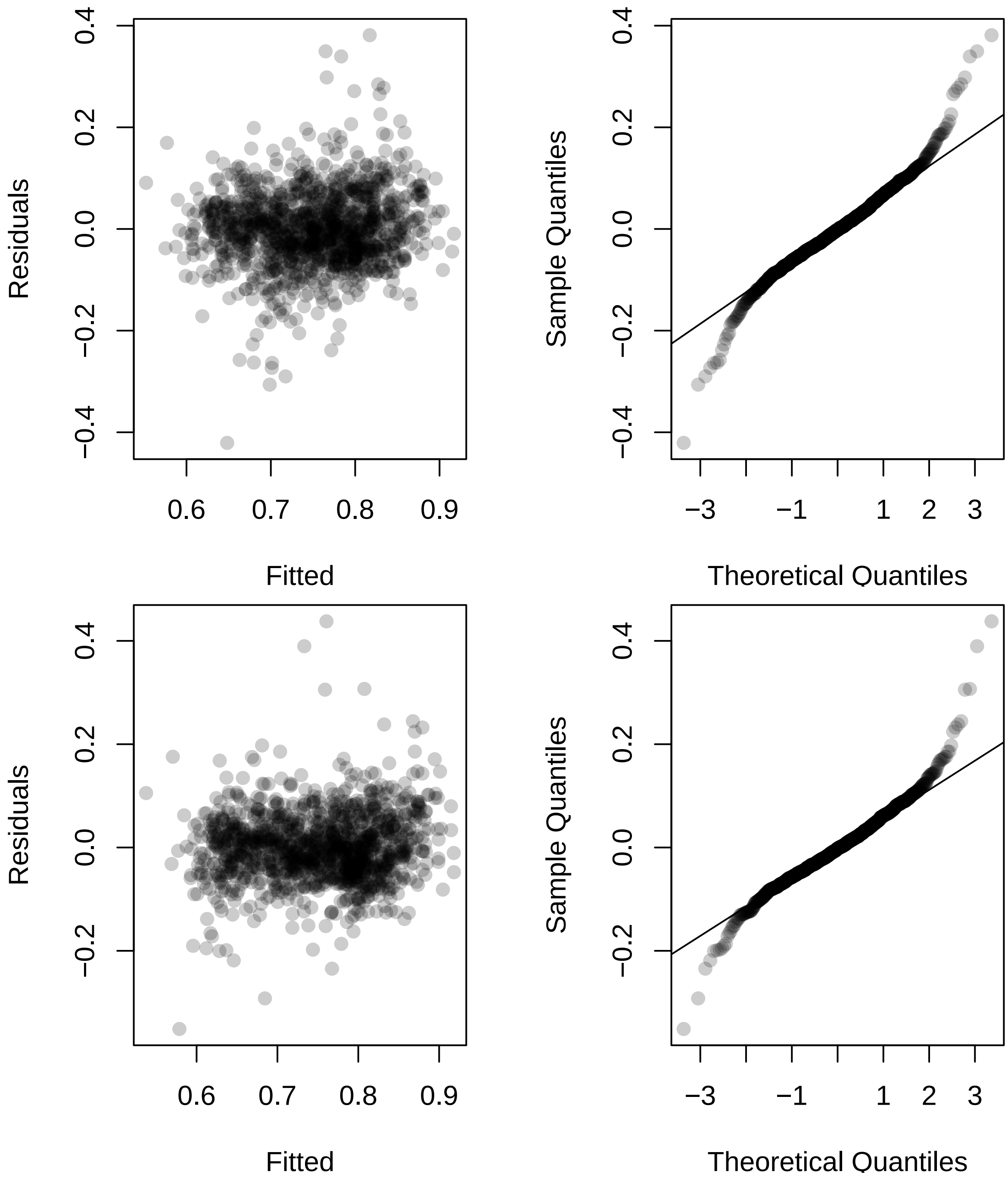
Residuals
Fitted
Sample Quantiles
Theoretical Quantiles
Residuals
Fitted
Sample Quantiles
Theoretical Quantiles

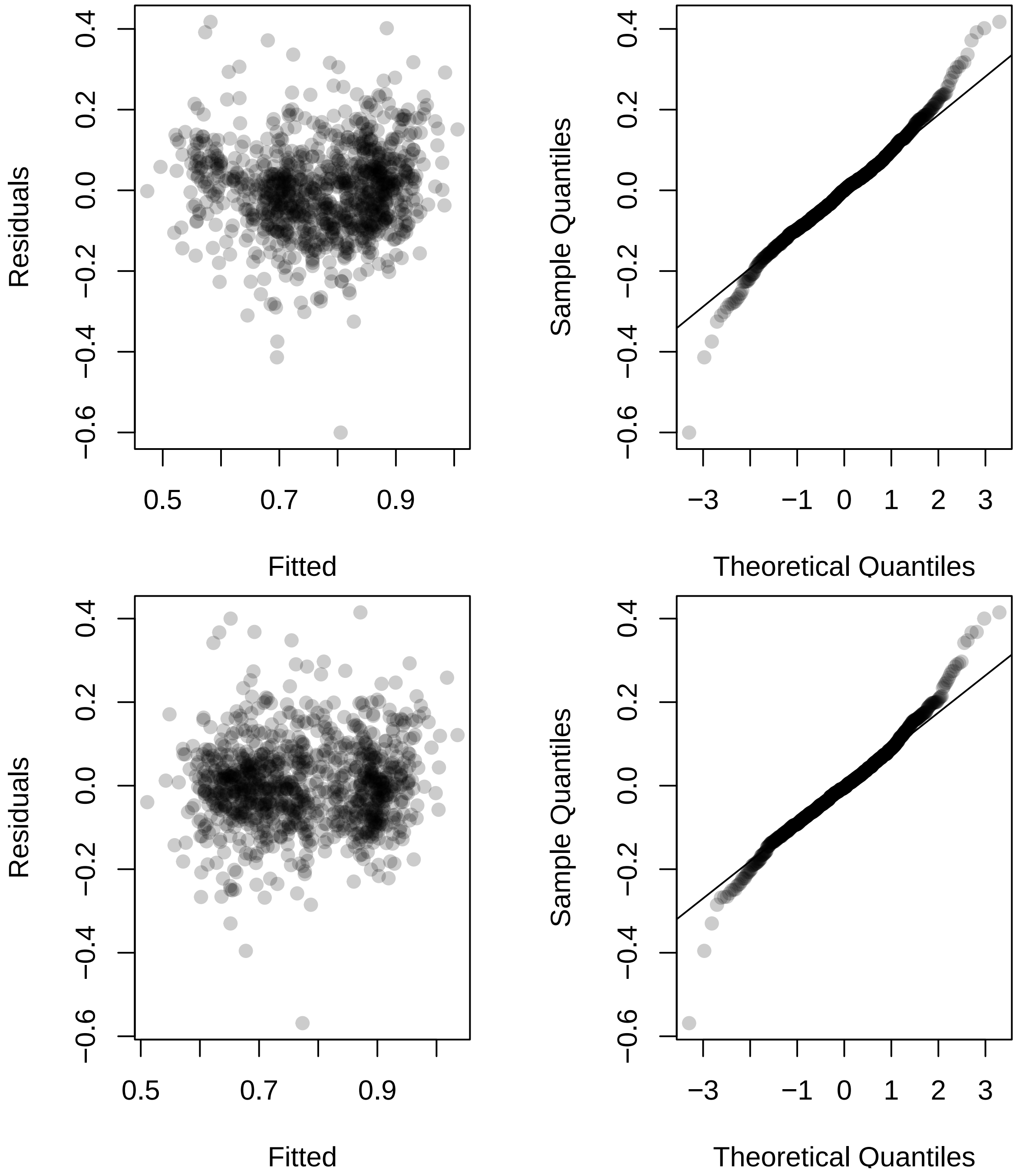
Residuals
Fitted
0.5
0.7
0.9
0.4
0.2
0.0
−0.2
−0.4
−0.6
Sample Quantiles
Theoretical Quantiles
−3
−1
0
1
2
3
Residuals
Fitted
Sample Quantiles
Theoretical Quantiles

##### Investigate genotypes

We now perform an AIC based procedure to find genotypes that differ from the RC reference level.

```
M = model.matrix(mAqI ~ ID + Date_num + I(Date_num^2), data = dat5)
M_full = model.matrix(mAqI ~ ID + Date_num + I(Date_num^2) +
                      Ta + VPD + Fsd + Precip + Precip_cum +
                      Ta_7day + VPD_7day + Fsd_7day + Precip_7day,
                 data = dat5)

M2021 = model.matrix(mAqI ~ ID + Date_num + I(Date_num^2),
                 data = dat5 %>% filter(year == 2021))
M2021_full = model.matrix(mAqI ~ ID + Date_num + I(Date_num^2) +
                      Ta + VPD + Fsd + Precip + Precip_cum +
                      Ta_7day + VPD_7day + Fsd_7day + Precip_7day,
                 data = dat5 %>% filter(year == 2021))

M2022 = model.matrix(mAqI ~ ID + Date_num + I(Date_num^2),
                 data = dat5 %>% filter(year == 2022))
M2022_full = model.matrix(mAqI ~ ID + Date_num + I(Date_num^2) +
                            Ta + VPD + Fsd + Precip + Precip_cum +
                            Ta_7day + VPD_7day + Fsd_7day + Precip_7day,
                 data = dat5 %>% filter(year == 2022))

ncores = detectCores() - 2
AIC_IDs_mAqI = do.call(rbind, mclapply(
  grep("IDNA", colnames(M)), function(j){
    M1 = M[, -j]
    foo = lmer(mAqI^(1/3) ~ -1 + M1 + (1|plot_number_year),
                data = dat5, REML = FALSE)
    M1_full = M_full[, -j]
    foo_full = lmer(mAqI^(1/3) ~ -1 + M1_full + (1|plot_number_year),
                data = dat5, REML = FALSE)
    M12021 = M2021[, -j]
    bar = lmer(mAqI^(1/3) ~ -1 + M12021 + (1|plot_number),
                data = dat5 %>% filter(year == 2021), REML = FALSE)
    M12021_full = M2021_full[, -j]
    bar_full = lmer(mAqI^(1/3) ~ -1 + M12021_full + (1|plot_number),
                data = dat5 %>% filter(year == 2021), REML = FALSE)
    M12022 = M2022[, -j]
    baz = lmer(mAqI^(1/3) ~ -1 + M12022 + (1|plot_number),
                data = dat5 %>% filter(year == 2022), REML = FALSE)
    M12022_full = M2022_full[, -j]
    baz_full = lmer(mAqI^(1/3) ~ -1 + M12022_full + (1|plot_number),
                data = dat5 %>% filter(year == 2022), REML = FALSE)

    c(AIC(m1_mAqI_cube) - AIC(foo),
      AIC(m1_mAqI_cube_full) - AIC(foo_full),
      AIC(m2_mAqI_cube) - AIC(bar),
      AIC(m2_mAqI_cube_full) - AIC(bar_full),
      AIC(m3_mAqI_cube) - AIC(baz),
      AIC(m3_mAqI_cube_full) - AIC(baz_full))
  }, mc.cores = ncores))
rownames(AIC_IDs_mAqI) = colnames(M)[grep("IDNA", colnames(M))]
```

Negative values indicate that the genotype in question provides better model fit when differentiated from the RC reference level.

```r
round(AIC_IDs_mAqI, 3)
```

```
##              [,1]    [,2]   [,3]   [,4]   [,5]   [,6]
## IDNAM10   1.975   1.991  1.806  1.847  1.783  1.904
## IDNAM11   2.081   1.697  1.994  1.977  1.601  1.635
## IDNAM12  -0.018  -1.663  0.490  0.290 -0.174 -0.044
## IDNAM13   2.063   1.655  1.456  1.486  1.980  1.969
## IDNAM14   1.760   1.995  1.951  1.963  1.631  1.798
## IDNAM15   2.054   1.906  1.997  1.977  1.875  1.856
## IDNAM17   0.972   0.020  1.384  1.350  0.202  0.475
## IDNAM18   1.746   1.031  1.446  1.380  1.770  1.682
## IDNAM2   -0.536  -0.428  0.660  0.671  1.140  1.074
## IDNAM22   1.971   1.943  1.996  1.998  1.848  1.927
## IDNAM23   1.750   1.987  0.994  0.655  0.576  0.739
## IDNAM24   1.816   1.409  1.813  1.803  1.435  1.480
## IDNAM25   1.460   1.524 -0.009  0.205  1.981  1.958
## IDNAM26  -0.970  -1.557  1.290  1.332 -2.173 -1.739
## IDNAM27   1.732   1.828  1.974  1.940  1.704  1.839
## IDNAM28   1.199   0.199  1.711  1.699 -0.136 -0.106
## IDNAM29   1.001   0.515  0.788  0.951  1.371  1.493
## IDNAM3   -2.005  -5.475  0.223  0.395 -5.121 -4.551
## IDNAM30   1.863   1.984  1.997  1.997  1.933  1.965
## IDNAM31   1.167   1.677  1.828  1.887  1.859  1.866
## IDNAM32  -2.876  -3.415  1.106  1.045 -3.482 -3.340
## IDNAM33 -10.111 -17.867 -6.753 -8.498 -7.946 -7.654
## IDNAM34  -1.428  -4.053 -1.765 -2.430  0.048 -0.307
## IDNAM36   1.661   0.698  1.700  1.686  0.538  0.633
## IDNAM37  -3.209  -5.375 -3.686 -3.893 -0.214 -0.214
## IDNAM38   1.497   0.782  0.799  0.717  1.673  1.775
## IDNAM39   1.498   1.847  1.923  1.934  1.902  1.885
## IDNAM4    2.011   1.445  1.879  1.792  1.487  1.533
## IDNAM40   0.415  -2.605  0.987  0.888 -2.495 -2.066
## IDNAM41  -0.732  -2.067  1.447  1.311 -2.574 -2.742
## IDNAM42  -0.152  -1.521  0.212  0.090  0.020  0.270
## IDNAM46  -1.165  -2.773 -1.409 -1.470 -0.057 -0.012
## IDNAM48   0.688   0.027  0.823  0.438  1.035  1.101
## IDNAM5    2.023   1.730  1.945  1.933  0.620  0.899
## IDNAM50  -1.737  -2.777 -1.145 -0.747 -0.451 -0.076
## IDNAM54  -3.581  -3.632 -3.643 -3.325  0.162  0.340
## IDNAM6    1.908   1.996  1.776  1.695  1.813  1.819
## IDNAM64  -2.592  -2.864 -2.114 -1.667  0.362  0.390
## IDNAM8    1.140   0.101  1.018  1.006  0.932  0.976
## IDNAM9    1.058   1.585  1.835  1.812  1.780  1.725
```

```r
sign(AIC_IDs_mAqI < 0)
```

```
##         [,1] [,2] [,3] [,4] [,5] [,6]
## IDNAM10    0    0    0    0    0    0
## IDNAM11    0    0    0    0    0    0
## IDNAM12    1    1    0    0    1    1
## IDNAM13    0    0    0    0    0    0
## IDNAM14    0    0    0    0    0    0
```

```
## IDNAM15    0    0    0    0    0    0
## IDNAM17    0    0    0    0    0    0
## IDNAM18    0    0    0    0    0    0
## IDNAM2     1    1    0    0    0    0
## IDNAM22    0    0    0    0    0    0
## IDNAM23    0    0    0    0    0    0
## IDNAM24    0    0    0    0    0    0
## IDNAM25    0    0    1    0    0    0
## IDNAM26    1    1    0    0    1    1
## IDNAM27    0    0    0    0    0    0
## IDNAM28    0    0    0    0    1    1
## IDNAM29    0    0    0    0    0    0
## IDNAM3     1    1    0    0    1    1
## IDNAM30    0    0    0    0    0    0
## IDNAM31    0    0    0    0    0    0
## IDNAM32    1    1    0    0    1    1
## IDNAM33    1    1    1    1    1    1
## IDNAM34    1    1    1    1    0    1
## IDNAM36    0    0    0    0    0    0
## IDNAM37    1    1    1    1    1    1
## IDNAM38    0    0    0    0    0    0
## IDNAM39    0    0    0    0    0    0
## IDNAM4     0    0    0    0    0    0
## IDNAM40    0    1    0    0    1    1
## IDNAM41    1    1    0    0    1    1
## IDNAM42    1    1    0    0    0    0
## IDNAM46    1    1    1    1    1    1
## IDNAM48    0    0    0    0    0    0
## IDNAM5     0    0    0    0    0    0
## IDNAM50    1    1    1    1    1    1
## IDNAM54    1    1    1    1    0    0
## IDNAM6     0    0    0    0    0    0
## IDNAM64    1    1    1    1    0    0
## IDNAM8     0    0    0    0    0    0
## IDNAM9     0    0    0    0    0    0
```

We now investigate the specific coefficient effects from our fitted models. Specific interest is in determining the sign of the effect. Here are the effect estimates from each fitted model:

```r
coefs_mAqI = round(cbind(summary(m1_mAqI_cube)$coefficients[2:41, 1],
                   summary(m1_mAqI_cube_full)$coefficients[2:41, 1],
                   summary(m2_mAqI_cube)$coefficients[2:41, 1],
                   summary(m2_mAqI_cube_full)$coefficients[2:41, 1],
                   summary(m3_mAqI_cube)$coefficients[2:41, 1],
                   summary(m3_mAqI_cube_full)$coefficients[2:41, 1]), 3)
coefs_mAqI
```

```
##            [,1]   [,2]   [,3]   [,4]   [,5]   [,6]
## IDNAM10   0.008 -0.002 -0.013 -0.011  0.017  0.011
## IDNAM11   0.000 -0.012 -0.002 -0.004 -0.022 -0.021
## IDNAM12  -0.034 -0.044 -0.035 -0.037 -0.052 -0.050
## IDNAM13  -0.003 -0.013 -0.021 -0.020 -0.005 -0.006
## IDNAM14   0.013  0.002 -0.006 -0.005  0.021  0.016
## IDNAM15  -0.004 -0.007 -0.002 -0.004 -0.012 -0.013
## IDNAM17  -0.024 -0.032 -0.022 -0.023 -0.046 -0.043
## IDNAM18  -0.013 -0.023 -0.021 -0.022 -0.017 -0.020
## IDNAM2    0.036  0.035  0.033  0.032  0.031  0.032
## IDNAM22   0.008 -0.005  0.002  0.001 -0.014 -0.009
## IDNAM23   0.013  0.003 -0.028 -0.033  0.041  0.039
## IDNAM24  -0.012 -0.017 -0.012 -0.012 -0.025 -0.024
## IDNAM25  -0.018 -0.016 -0.040 -0.038  0.005  0.007
## IDNAM26  -0.041 -0.043 -0.024 -0.023 -0.070 -0.066
## IDNAM27   0.014 -0.010 -0.005 -0.007 -0.020 -0.014
## IDNAM28  -0.022 -0.030 -0.015 -0.015 -0.051 -0.051
## IDNAM29  -0.024 -0.028 -0.031 -0.029 -0.028 -0.025
## IDNAM3   -0.047 -0.062 -0.038 -0.036 -0.092 -0.088
## IDNAM30   0.011 -0.003  0.001  0.001 -0.009 -0.007
## IDNAM31   0.022  0.013  0.012  0.010  0.013  0.013
## IDNAM32   0.051  0.052  0.027  0.027  0.080  0.080
## IDNAM33  -0.092 -0.113 -0.104 -0.112 -0.113 -0.111
## IDNAM34  -0.045 -0.058 -0.055 -0.060 -0.054 -0.059
## IDNAM36  -0.015 -0.026 -0.015 -0.016 -0.042 -0.040
## IDNAM37  -0.053 -0.062 -0.068 -0.069 -0.052 -0.052
## IDNAM38  -0.018 -0.025 -0.031 -0.032 -0.020 -0.016
## IDNAM39   0.018  0.009  0.008  0.007  0.011  0.012
## IDNAM4   -0.006 -0.017 -0.010 -0.013 -0.025 -0.024
## IDNAM40  -0.030 -0.049 -0.029 -0.030 -0.074 -0.071
## IDNAM41  -0.039 -0.046 -0.021 -0.023 -0.075 -0.077
## IDNAM42  -0.034 -0.042 -0.038 -0.039 -0.048 -0.045
## IDNAM46  -0.041 -0.050 -0.052 -0.052 -0.050 -0.049
## IDNAM48  -0.027 -0.032 -0.031 -0.035 -0.034 -0.033
## IDNAM5    0.006 -0.012  0.007  0.007 -0.041 -0.037
## IDNAM50  -0.045 -0.049 -0.051 -0.047 -0.054 -0.049
## IDNAM54  -0.055 -0.054 -0.068 -0.066 -0.046 -0.044
## IDNAM6    0.010 -0.001 -0.013 -0.016  0.015  0.015
## IDNAM64  -0.049 -0.050 -0.058 -0.054 -0.043 -0.043
## IDNAM8   -0.022 -0.031 -0.028 -0.028 -0.036 -0.035
## IDNAM9    0.024  0.015  0.012  0.012  0.017  0.019
```

Here are the corresponding signs:

```
sign(coefs_mAqI > 0)
```

```
##          [,1] [,2] [,3] [,4] [,5] [,6]
## IDNAM10    1    0    0    0    1    1
## IDNAM11    0    0    0    0    0    0
## IDNAM12    0    0    0    0    0    0
## IDNAM13    0    0    0    0    0    0
## IDNAM14    1    1    0    0    1    1
## IDNAM15    0    0    0    0    0    0
## IDNAM17    0    0    0    0    0    0
## IDNAM18    0    0    0    0    0    0
## IDNAM2     1    1    1    1    1    1
## IDNAM22    1    0    1    1    0    0
## IDNAM23    1    1    0    0    1    1
## IDNAM24    0    0    0    0    0    0
## IDNAM25    0    0    0    0    1    1
## IDNAM26    0    0    0    0    0    0
## IDNAM27    1    0    0    0    0    0
## IDNAM28    0    0    0    0    0    0
## IDNAM29    0    0    0    0    0    0
## IDNAM3     0    0    0    0    0    0
## IDNAM30    1    0    1    1    0    0
## IDNAM31    1    1    1    1    1    1
## IDNAM32    1    1    1    1    1    1
## IDNAM33    0    0    0    0    0    0
## IDNAM34    0    0    0    0    0    0
## IDNAM36    0    0    0    0    0    0
## IDNAM37    0    0    0    0    0    0
## IDNAM38    0    0    0    0    0    0
## IDNAM39    1    1    1    1    1    1
## IDNAM4     0    0    0    0    0    0
## IDNAM40    0    0    0    0    0    0
## IDNAM41    0    0    0    0    0    0
## IDNAM42    0    0    0    0    0    0
## IDNAM46    0    0    0    0    0    0
## IDNAM48    0    0    0    0    0    0
## IDNAM5     1    0    1    1    0    0
## IDNAM50    0    0    0    0    0    0
## IDNAM54    0    0    0    0    0    0
## IDNAM6     1    0    0    0    1    1
## IDNAM64    0    0    0    0    0    0
## IDNAM8     0    0    0    0    0    0
## IDNAM9     1    1    1    1    1    1
```

Below are signs that indicate which genotypes yield better model fit when their effect is differentiated from the RC reference level. A 1 indicates a positive effect, a -1 indicates a negative effect, and a 0 indicates that the genotype was not determined to provide better model when differentiated from the RC reference level.

```
inference_mAqI = sign(coefs_mAqI) * sign(AIC_IDs_mAqI < 0)
inference_mAqI
```

```
##          [,1] [,2] [,3] [,4] [,5] [,6]
## IDNAM10     0    0    0    0    0    0
## IDNAM11     0    0    0    0    0    0
## IDNAM12    -1   -1    0    0   -1   -1
## IDNAM13     0    0    0    0    0    0
## IDNAM14     0    0    0    0    0    0
## IDNAM15     0    0    0    0    0    0
## IDNAM17     0    0    0    0    0    0
## IDNAM18     0    0    0    0    0    0
## IDNAM2      1    1    0    0    0    0
## IDNAM22     0    0    0    0    0    0
## IDNAM23     0    0    0    0    0    0
## IDNAM24     0    0    0    0    0    0
## IDNAM25     0    0   -1    0    0    0
## IDNAM26    -1   -1    0    0   -1   -1
## IDNAM27     0    0    0    0    0    0
## IDNAM28     0    0    0    0   -1   -1
## IDNAM29     0    0    0    0    0    0
## IDNAM3     -1   -1    0    0   -1   -1
## IDNAM30     0    0    0    0    0    0
## IDNAM31     0    0    0    0    0    0
## IDNAM32     1    1    0    0    1    1
## IDNAM33    -1   -1   -1   -1   -1   -1
## IDNAM34    -1   -1   -1   -1    0   -1
## IDNAM36     0    0    0    0    0    0
## IDNAM37    -1   -1   -1   -1   -1   -1
## IDNAM38     0    0    0    0    0    0
## IDNAM39     0    0    0    0    0    0
## IDNAM4      0    0    0    0    0    0
## IDNAM40     0   -1    0    0   -1   -1
## IDNAM41    -1   -1    0    0   -1   -1
## IDNAM42    -1   -1    0    0    0    0
## IDNAM46    -1   -1   -1   -1   -1   -1
## IDNAM48     0    0    0    0    0    0
## IDNAM5      0    0    0    0    0    0
## IDNAM50    -1   -1   -1   -1   -1   -1
## IDNAM54    -1   -1   -1   -1    0    0
## IDNAM6      0    0    0    0    0    0
## IDNAM64    -1   -1   -1   -1    0    0
## IDNAM8      0    0    0    0    0    0
## IDNAM9      0    0    0    0    0    0
```

**Results**

Here are the effect size estimates for genotypes that were to determined to differentiate from the RC reference level in all considered models and had the same estimated sign across all considered models.

```
coefs_mAqI[abs(rowSums(inference_mAqI)) == 6, ]
```

```
##           [,1]   [,2]   [,3]   [,4]   [,5]   [,6]
## IDNAM33 -0.092 -0.113 -0.104 -0.112 -0.113 -0.111
## IDNAM37 -0.053 -0.062 -0.068 -0.069 -0.052 -0.052
## IDNAM46 -0.041 -0.050 -0.052 -0.052 -0.050 -0.049
## IDNAM50 -0.045 -0.049 -0.051 -0.047 -0.054 -0.049
```

##### AqM response

```
m1_mAqM = lmer(mAqM ~ ID + Date_num + I(Date_num^2) +
               (1|plot_number_year),
           data = dat5, REML = FALSE, control = lmerControl(optimizer ="Nelder_Mead"))
m1_mAqM_full = lmer(mAqM ~ ID + Date_num + I(Date_num^2) +
                      Ta + VPD + Fsd + Precip + Precip_cum +
                      Ta_7day + VPD_7day + Fsd_7day + Precip_7day +
                      (1|plot_number_year),
           data = dat5, REML = FALSE, control = lmerControl(optimizer ="Nelder_Mead"))
AIC(m1_mAqM)
```

```
## [1] 1562.258
```

```
AIC(m1_mAqM_full)
```

```
## [1] 451.2039
```

```
m2_mAqM = lmer(mAqM ~ ID + Date_num + I(Date_num^2) +
               (1|plot_number),
           data = dat5 %>% filter(year == 2021), REML = FALSE,
           control = lmerControl(optimizer ="Nelder_Mead"))
m2_mAqM_full = lmer(mAqM ~ ID + Date_num + I(Date_num^2) +
                      Ta + VPD + Fsd + Precip + Precip_cum +
                      Ta_7day + VPD_7day + Fsd_7day + Precip_7day +
                      (1|plot_number),
           data = dat5 %>% filter(year == 2021), REML = FALSE,
           control = lmerControl(optimizer ="Nelder_Mead"))
AIC(m2_mAqM)
```

```
## [1] -53.32485
```

```
AIC(m2_mAqM_full)
```

```
## [1] -179.2898
```

```
m3_mAqM = lmer(mAqM ~ ID + Date_num + I(Date_num^2) +
               (1|plot_number),
           data = dat5 %>% filter(year == 2022), REML = FALSE,
           control = lmerControl(optimizer ="Nelder_Mead"))
m3_mAqM_full = lmer(mAqM ~ ID + Date_num + I(Date_num^2) +
                      Ta + VPD + Fsd + Precip + Precip_cum +
                      Ta_7day + VPD_7day + Fsd_7day + Precip_7day +
                      (1|plot_number),
           data = dat5 %>% filter(year == 2022), REML = FALSE,
           control = lmerControl(optimizer ="Nelder_Mead"))
AIC(m3_mAqM)
```

```
## [1] 681.0196
```

```
AIC(m3_mAqM_full)
```

```
## [1] 468.6349
```

**Diagnostics**

Diagnostic plots for the AqM response are provided (small model followed by full model for models 1 through 3). Modeling assumptions appear to be satisfied.

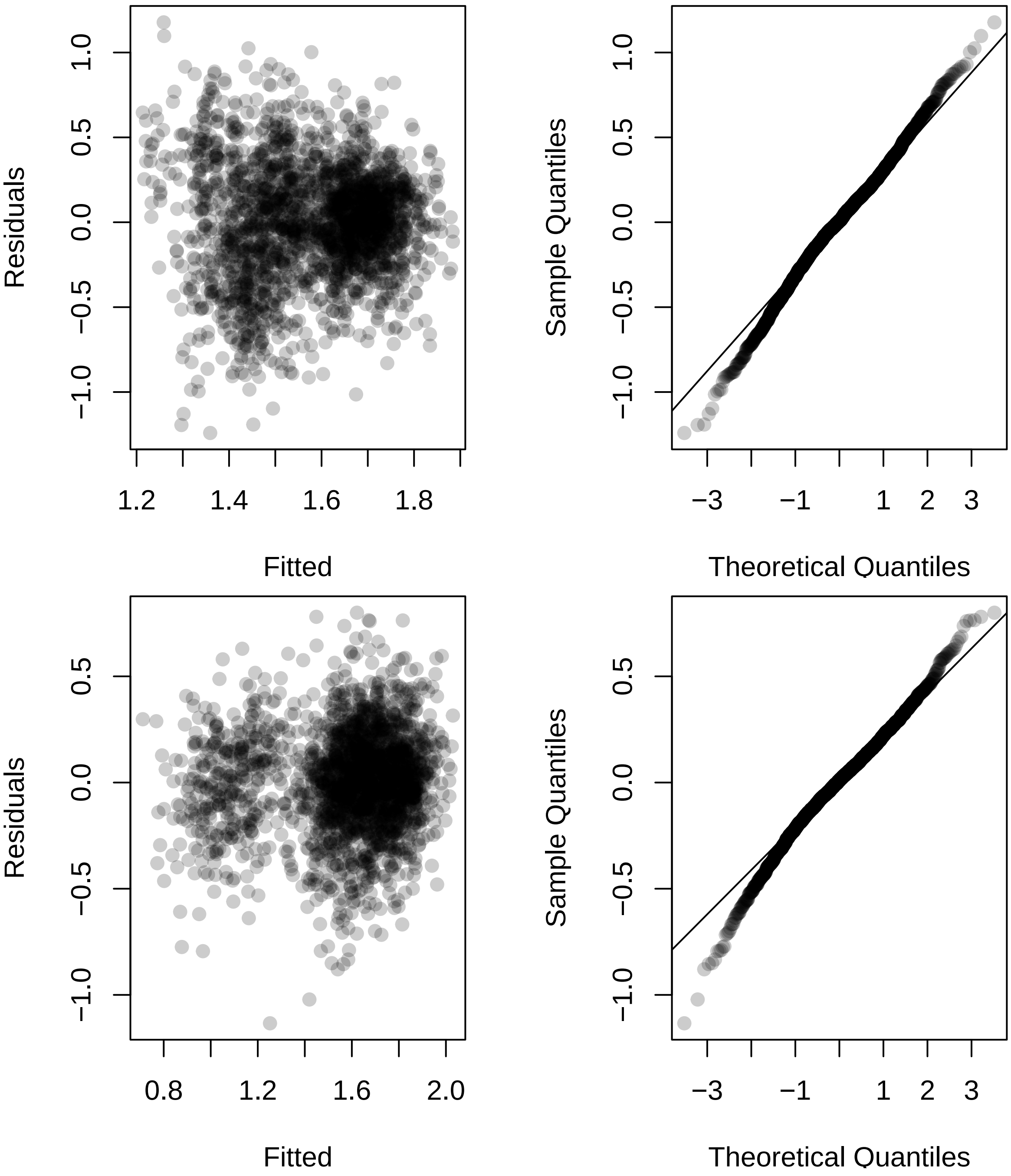

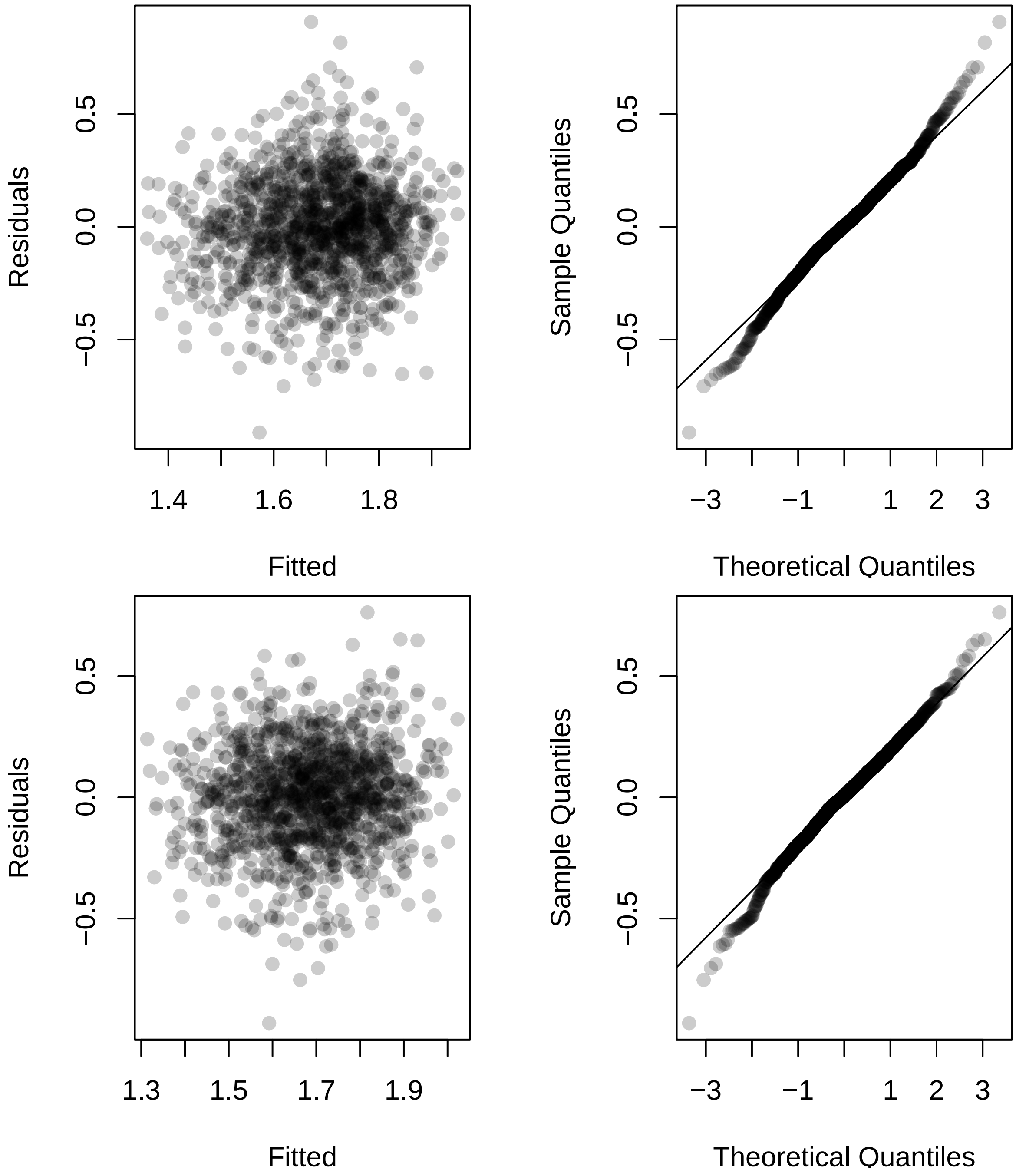

Residuals
0.5
0.0
−0.5
1.4
1.6
1.8
Fitted
Sample Quantiles
0.5
0.0
−0.5
−3
−1
1
2
3
Theoretical Quantiles
Residuals
0.5
0.0
−0.5
1.3
1.5
1.7
1.9
Fitted
Sample Quantiles
0.5
0.0
−0.5
−3
−1
1
2
3
Theoretical Quantiles

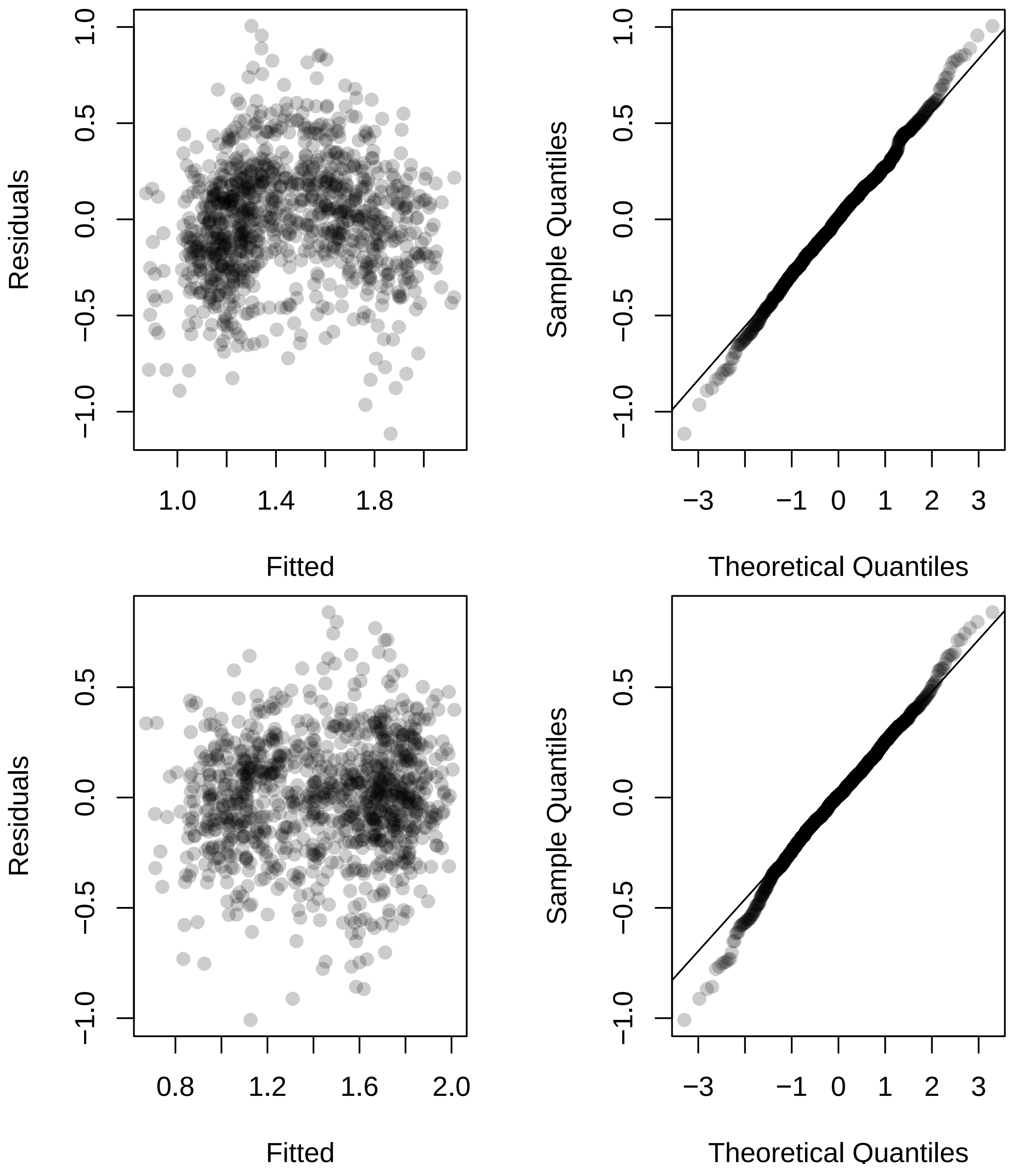

Residuals
Fitted
Sample Quantiles
Theoretical Quantiles
Residuals
Fitted
Sample Quantiles
Theoretical Quantiles

##### Investigate genotypes

We now perform an AIC based procedure to find genotypes that differ from the RC reference level.

```
## AIC for each ID variable from full AqM fixed-effects model
M = model.matrix(mAqM ~ ID + Date_num + I(Date_num^2), data = dat5)
M_full = model.matrix(mAqM ~ ID + Date_num + I(Date_num^2) +
                      Ta + VPD + Fsd + Precip + Precip_cum +
                      Ta_7day + VPD_7day + Fsd_7day + Precip_7day,
                 data = dat5)

M2021 = model.matrix(mAqM ~ ID + Date_num + I(Date_num^2),
                 data = dat5 %>% filter(year == 2021))
M2021_full = model.matrix(mAqM ~ ID + Date_num + I(Date_num^2) +
                      Ta + VPD + Fsd + Precip + Precip_cum +
                      Ta_7day + VPD_7day + Fsd_7day + Precip_7day,
                 data = dat5 %>% filter(year == 2021))

M2022 = model.matrix(mAqM ~ ID + Date_num + I(Date_num^2),
                 data = dat5 %>% filter(year == 2022))
M2022_full = model.matrix(mAqM ~ ID + Date_num + I(Date_num^2) +
                            Ta + VPD + Fsd + Precip + Precip_cum +
                            Ta_7day + VPD_7day + Fsd_7day + Precip_7day,
                 data = dat5 %>% filter(year == 2022))

ncores = detectCores() - 2
AIC_IDs_mAqM = do.call(rbind, mclapply(
  grep("IDNA", colnames(M)), function(j){
    M1 = M[, -j]
    foo = lmer(mAqM ~ -1 + M1 + (1|plot_number_year),
               data = dat5, REML = FALSE)
    M1_full = M_full[, -j]
    foo_full = lmer(mAqM ~ -1 + M1_full + (1|plot_number_year),
               data = dat5, REML = FALSE)
    M12021 = M2021[, -j]
    bar = lmer(mAqM ~ -1 + M12021 + (1|plot_number),
               data = dat5 %>% filter(year == 2021), REML = FALSE)
    M12021_full = M2021_full[, -j]
    bar_full = lmer(mAqM ~ -1 + M12021_full + (1|plot_number),
               data = dat5 %>% filter(year == 2021), REML = FALSE)
    M12022 = M2022[, -j]
    baz = lmer(mAqM ~ -1 + M12022 + (1|plot_number),
               data = dat5 %>% filter(year == 2022), REML = FALSE)
    M12022_full = M2022_full[, -j]
    baz_full = lmer(mAqM ~ -1 + M12022_full + (1|plot_number),
               data = dat5 %>% filter(year == 2022), REML = FALSE)

    c(AIC(m1_mAqM) - AIC(foo),
      AIC(m1_mAqM_full) - AIC(foo_full),
      AIC(m2_mAqM) - AIC(bar),
      AIC(m2_mAqM_full) - AIC(bar_full),
      AIC(m3_mAqM) - AIC(baz),
      AIC(m3_mAqM_full) - AIC(baz_full))
  }, mc.cores = ncores))
```

```r
rownames(AIC_IDs_mAqM) = colnames(M)[grep("IDNA", colnames(M))]
```

Negative values indicate that the genotype in question provides better model fit when differentiated from the RC reference level.

```r
round(AIC_IDs_mAqM, 3)
```

```
##            [,1]   [,2]   [,3]   [,4]   [,5]   [,6]
## IDNAM10  1.654  0.455  1.889  1.918  0.645 -0.096
## IDNAM11  1.908  1.613  1.614  1.518  1.892  1.903
## IDNAM12  1.993  1.829  1.979  1.958  1.855  1.867
## IDNAM13  1.962  1.815  1.969  1.956  1.583  1.492
## IDNAM14  0.585  1.149  1.918  1.888  0.255  0.806
## IDNAM15  0.309 -0.158  1.870  1.788 -0.258 -0.381
## IDNAM17  0.667  0.156  1.984  1.993 -1.229 -0.651
## IDNAM18  1.864  1.803  1.433  1.439  1.998  2.000
## IDNAM2   1.953  1.964  1.877  1.888  1.983  1.991
## IDNAM22 -1.277 -0.944 -3.681 -3.674  1.743  1.599
## IDNAM23 -0.200  0.609  1.808  1.915  0.147  0.427
## IDNAM24  1.471  1.366  0.535  0.575  1.917  1.923
## IDNAM25  1.618  1.491  1.105  1.107  1.865  1.904
## IDNAM26  0.463  0.904 -0.371 -0.372  1.947  1.948
## IDNAM27  1.965  1.396  0.581  0.407  1.905  1.980
## IDNAM28  1.830  1.390  1.837  1.833  0.333  0.204
## IDNAM29  1.967  1.999  1.990  1.976  1.980  2.000
## IDNAM3  -1.940 -3.282  0.798  0.950 -3.567 -2.835
## IDNAM30  1.999  1.906  1.177  1.175  1.965  1.936
## IDNAM31  1.377  0.037 -0.022 -0.310  1.284  1.238
## IDNAM32 -0.426 -0.458  1.994  1.988 -2.027 -1.770
## IDNAM33  1.383  2.000  1.521  1.906  1.991  1.992
## IDNAM34  1.556  1.292 -1.165 -1.702  1.784  1.857
## IDNAM36  0.101 -0.815 -0.099 -0.100  0.466  0.461
## IDNAM37  1.621  1.904  2.000  1.995  1.726  1.769
## IDNAM38  1.978  1.986  1.910  1.913  1.859  1.908
## IDNAM39  1.774  1.993  1.417  1.520  1.873  1.882
## IDNAM4   1.659  0.586 -2.509 -3.173  1.996  2.000
## IDNAM40  1.522  0.862  0.011 -0.252  1.784  1.851
## IDNAM41  1.957  1.588  1.323  1.438  0.285  0.100
## IDNAM42  1.996  1.955  1.747  1.766  1.554  1.754
## IDNAM46  1.971  1.732  0.740  0.779 -0.061 -0.198
## IDNAM48  1.994  1.817  1.624  1.410  1.997  1.995
## IDNAM5   1.573  0.731 -1.312 -1.314  1.879  1.957
## IDNAM50  1.823  1.823  1.375  1.545  1.964  1.995
## IDNAM54 -3.619 -3.528 -6.103 -5.758  0.466  0.569
## IDNAM6  -0.924 -1.734 -0.621 -0.840  0.201  0.078
## IDNAM64  1.902  1.936  1.010  0.728  1.014  1.050
## IDNAM8  -0.554 -1.335 -2.965 -3.105  1.383  1.338
## IDNAM9   1.383  1.893 -0.585 -0.617  1.448  1.520
```

```r
sign(AIC_IDs_mAqM < 0)
```

```
##         [,1] [,2] [,3] [,4] [,5] [,6]
## IDNAM10    0    0    0    0    0    1
## IDNAM11    0    0    0    0    0    0
## IDNAM12    0    0    0    0    0    0
```

```
## IDNAM13    0    0    0    0    0    0
## IDNAM14    0    0    0    0    0    0
## IDNAM15    0    1    0    0    1    1
## IDNAM17    0    0    0    0    1    1
## IDNAM18    0    0    0    0    0    0
## IDNAM2     0    0    0    0    0    0
## IDNAM22    1    1    1    1    0    0
## IDNAM23    1    0    0    0    0    0
## IDNAM24    0    0    0    0    0    0
## IDNAM25    0    0    0    0    0    0
## IDNAM26    0    0    1    1    0    0
## IDNAM27    0    0    0    0    0    0
## IDNAM28    0    0    0    0    0    0
## IDNAM29    0    0    0    0    0    0
## IDNAM3     1    1    0    0    1    1
## IDNAM30    0    0    0    0    0    0
## IDNAM31    0    0    1    1    0    0
## IDNAM32    1    1    0    0    1    1
## IDNAM33    0    0    0    0    0    0
## IDNAM34    0    0    1    1    0    0
## IDNAM36    0    1    1    1    0    0
## IDNAM37    0    0    0    0    0    0
## IDNAM38    0    0    0    0    0    0
## IDNAM39    0    0    0    0    0    0
## IDNAM4     0    0    1    1    0    0
## IDNAM40    0    0    0    1    0    0
## IDNAM41    0    0    0    0    0    0
## IDNAM42    0    0    0    0    0    0
## IDNAM46    0    0    0    0    1    1
## IDNAM48    0    0    0    0    0    0
## IDNAM5     0    0    1    1    0    0
## IDNAM50    0    0    0    0    0    0
## IDNAM54    1    1    1    1    0    0
## IDNAM6     1    1    1    1    0    0
## IDNAM64    0    0    0    0    0    0
## IDNAM8     1    1    1    1    0    0
## IDNAM9     0    0    1    1    0    0
```

We now investigate the specific coefficient effects from our fitted models. Specific interest is in determining the sign of the effect. Here are the effect estimates from each fitted model:

```
coefs_mAqM = round(cbind(summary(m1_mAqM)$coefficients[2:41, 1],
                   summary(m1_mAqM_full)$coefficients[2:41, 1],
                   summary(m2_mAqM)$coefficients[2:41, 1],
                   summary(m2_mAqM_full)$coefficients[2:41, 1],
                   summary(m3_mAqM)$coefficients[2:41, 1],
                   summary(m3_mAqM_full)$coefficients[2:41, 1]), 3)
coefs_mAqM
```

```
##            [,1]   [,2]   [,3]   [,4]   [,5]   [,6]
## IDNAM10   0.037  0.080  0.021  0.018  0.126  0.155
## IDNAM11   0.019  0.040  0.038  0.043  0.035  0.033
## IDNAM12   0.005  0.026  0.009  0.013  0.041  0.039
## IDNAM13   0.012  0.027 -0.011 -0.013  0.067  0.074
## IDNAM14  -0.073 -0.059 -0.018 -0.021 -0.139 -0.115
## IDNAM15   0.079  0.092  0.022  0.029  0.154  0.159
## IDNAM17   0.072  0.087  0.008  0.005  0.190  0.172
## IDNAM18   0.023  0.029  0.048  0.047 -0.005  0.001
## IDNAM2    0.013  0.012  0.022  0.021  0.013  0.010
## IDNAM22  -0.113 -0.110 -0.151 -0.150 -0.054 -0.067
## IDNAM23  -0.091 -0.075 -0.027 -0.018 -0.142 -0.131
## IDNAM24   0.044  0.050  0.075  0.074  0.030  0.029
## IDNAM25   0.038  0.045  0.059  0.059  0.039  0.033
## IDNAM26  -0.077 -0.067 -0.097 -0.097 -0.024 -0.024
## IDNAM27   0.012  0.050  0.074  0.078  0.034  0.015
## IDNAM28   0.025  0.050 -0.025 -0.025  0.137  0.142
## IDNAM29  -0.011 -0.002 -0.006 -0.010  0.015  0.001
## IDNAM3    0.124  0.147  0.070  0.065  0.247  0.230
## IDNAM30  -0.001  0.019  0.056  0.056 -0.020 -0.027
## IDNAM31   0.049  0.090  0.089  0.094  0.090  0.093
## IDNAM32  -0.095 -0.099 -0.005 -0.007 -0.210 -0.204
## IDNAM33  -0.055 -0.001 -0.055 -0.024  0.010  0.009
## IDNAM34   0.043  0.056  0.111  0.120 -0.055 -0.044
## IDNAM36   0.084  0.106  0.089  0.089  0.130  0.130
## IDNAM37  -0.038 -0.020  0.000  0.004 -0.055 -0.051
## IDNAM38  -0.009  0.007 -0.019 -0.018  0.039  0.032
## IDNAM39  -0.029 -0.005 -0.047 -0.043  0.037  0.036
## IDNAM4    0.037  0.077  0.133  0.142  0.007  0.002
## IDNAM40   0.043  0.068  0.088  0.093  0.049  0.040
## IDNAM41   0.013  0.041 -0.051 -0.046  0.140  0.147
## IDNAM42   0.004  0.013 -0.031 -0.030  0.070  0.052
## IDNAM46   0.010  0.033 -0.069 -0.068  0.152  0.157
## IDNAM48  -0.005  0.027  0.038  0.048  0.005  0.007
## IDNAM5    0.041  0.072  0.113  0.113  0.037  0.022
## IDNAM50   0.026  0.027  0.050  0.042  0.020  0.007
## IDNAM54   0.146  0.149  0.179  0.174  0.129  0.124
## IDNAM6    0.105  0.123  0.101  0.104  0.141  0.146
## IDNAM64   0.019  0.016 -0.062 -0.070  0.102  0.101
## IDNAM8    0.100  0.117  0.140  0.141  0.083  0.086
## IDNAM9   -0.050 -0.021 -0.102 -0.102  0.082  0.076
```

Here are the corresponding signs:

```r
sign(coefs_mAqM > 0)
```

```
##          [,1] [,2] [,3] [,4] [,5] [,6]
## IDNAM10     1    1    1    1    1    1
## IDNAM11     1    1    1    1    1    1
## IDNAM12     1    1    1    1    1    1
## IDNAM13     1    1    0    0    1    1
## IDNAM14     0    0    0    0    0    0
## IDNAM15     1    1    1    1    1    1
## IDNAM17     1    1    1    1    1    1
## IDNAM18     1    1    1    1    0    1
## IDNAM2      1    1    1    1    1    1
## IDNAM22     0    0    0    0    0    0
## IDNAM23     0    0    0    0    0    0
## IDNAM24     1    1    1    1    1    1
## IDNAM25     1    1    1    1    1    1
## IDNAM26     0    0    0    0    0    0
## IDNAM27     1    1    1    1    1    1
## IDNAM28     1    1    0    0    1    1
## IDNAM29     0    0    0    0    1    1
## IDNAM3      1    1    1    1    1    1
## IDNAM30     0    1    1    1    0    0
## IDNAM31     1    1    1    1    1    1
## IDNAM32     0    0    0    0    0    0
## IDNAM33     0    0    0    0    1    1
## IDNAM34     1    1    1    1    0    0
## IDNAM36     1    1    1    1    1    1
## IDNAM37     0    0    0    1    0    0
## IDNAM38     0    1    0    0    1    1
## IDNAM39     0    0    0    0    1    1
## IDNAM4      1    1    1    1    1    1
## IDNAM40     1    1    1    1    1    1
## IDNAM41     1    1    0    0    1    1
## IDNAM42     1    1    0    0    1    1
## IDNAM46     1    1    0    0    1    1
## IDNAM48     0    1    1    1    1    1
## IDNAM5      1    1    1    1    1    1
## IDNAM50     1    1    1    1    1    1
## IDNAM54     1    1    1    1    1    1
## IDNAM6      1    1    1    1    1    1
## IDNAM64     1    1    0    0    1    1
## IDNAM8      1    1    1    1    1    1
## IDNAM9      0    0    0    0    1    1
```

Below are signs that indicate which genotypes yield better model fit when their effect is differentiated from the RC reference level. A 1 indicates a positive effect, a -1 indicates a negative effect, and a 0 indicates that the genotype was not determined to provide better model when differentiated from the RC reference level.

```r
inference_mAqM = sign(coefs_mAqM) * sign(AIC_IDs_mAqM < 0)
inference_mAqM
```

```
##          [,1] [,2] [,3] [,4] [,5] [,6]
## IDNAM10     0    0    0    0    0    1
## IDNAM11     0    0    0    0    0    0
## IDNAM12     0    0    0    0    0    0
## IDNAM13     0    0    0    0    0    0
## IDNAM14     0    0    0    0    0    0
## IDNAM15     0    1    0    0    1    1
## IDNAM17     0    0    0    0    1    1
## IDNAM18     0    0    0    0    0    0
## IDNAM2      0    0    0    0    0    0
## IDNAM22    -1   -1   -1   -1    0    0
## IDNAM23    -1    0    0    0    0    0
## IDNAM24     0    0    0    0    0    0
## IDNAM25     0    0    0    0    0    0
## IDNAM26     0    0   -1   -1    0    0
## IDNAM27     0    0    0    0    0    0
## IDNAM28     0    0    0    0    0    0
## IDNAM29     0    0    0    0    0    0
## IDNAM3      1    1    0    0    1    1
## IDNAM30     0    0    0    0    0    0
## IDNAM31     0    0    1    1    0    0
## IDNAM32    -1   -1    0    0   -1   -1
## IDNAM33     0    0    0    0    0    0
## IDNAM34     0    0    1    1    0    0
## IDNAM36     0    1    1    1    0    0
## IDNAM37     0    0    0    0    0    0
## IDNAM38     0    0    0    0    0    0
## IDNAM39     0    0    0    0    0    0
## IDNAM4      0    0    1    1    0    0
## IDNAM40     0    0    0    1    0    0
## IDNAM41     0    0    0    0    0    0
## IDNAM42     0    0    0    0    0    0
## IDNAM46     0    0    0    0    1    1
## IDNAM48     0    0    0    0    0    0
## IDNAM5      0    0    1    1    0    0
## IDNAM50     0    0    0    0    0    0
## IDNAM54     1    1    1    1    0    0
## IDNAM6      1    1    1    1    0    0
## IDNAM64     0    0    0    0    0    0
## IDNAM8      1    1    1    1    0    0
## IDNAM9      0    0   -1   -1    0    0
```

**Results**

Here are the effect size estimates for genotypes that were to determined to differentiate from the RC reference level in all considered models and had the same estimated sign across all considered models. There are no such genotypes for the AqM response

```
coefs_mAqM[abs(rowSums(inference_mAqM)) == 6, ]
```

```
##      [,1] [,2] [,3] [,4] [,5] [,6]
```

##### tqE response

```
m1_mtqE = lmer(mtqE ~ ID + Date_num + I(Date_num^2) +
               (1|plot_number_year),
          data = dat5, REML = FALSE, control = lmerControl(optimizer ="Nelder_Mead"))
m1_mtqE_full = lmer(mtqE ~ ID + Date_num + I(Date_num^2) +
                      Ta + VPD + Fsd + Precip + Precip_cum +
                      Ta_7day + VPD_7day + Fsd_7day + Precip_7day +
                      (1|plot_number_year),
          data = dat5, REML = FALSE, control = lmerControl(optimizer ="Nelder_Mead"))
AIC(m1_mtqE)
```

```
## [1] 1453.057
```

```
AIC(m1_mtqE_full)
```

```
## [1] 794.7726
```

```
m2_mtqE = lmer(mtqE ~ ID + Date_num + I(Date_num^2) +
               (1|plot_number),
          data = dat5 %>% filter(year == 2021), REML = FALSE,
          control = lmerControl(optimizer ="Nelder_Mead"))
m2_mtqE_full = lmer(mtqE ~ ID + Date_num + I(Date_num^2) +
                      Ta + VPD + Fsd + Precip + Precip_cum +
                      Ta_7day + VPD_7day + Fsd_7day + Precip_7day +
                      (1|plot_number),
          data = dat5 %>% filter(year == 2021), REML = FALSE,
          control = lmerControl(optimizer ="Nelder_Mead"))
AIC(m2_mtqE)
```

```
## [1] 190.6442
```

```
AIC(m2_mtqE_full)
```

```
## [1] 65.54412
```

```
m3_mtqE = lmer(mtqE ~ ID + Date_num + I(Date_num^2) +
               (1|plot_number),
          data = dat5 %>% filter(year == 2022), REML = FALSE,
          control = lmerControl(optimizer ="Nelder_Mead"))
m3_mtqE_full = lmer(mtqE ~ ID + Date_num + I(Date_num^2) +
                      Ta + VPD + Fsd + Precip + Precip_cum +
                      Ta_7day + VPD_7day + Fsd_7day + Precip_7day +
                      (1|plot_number),
          data = dat5 %>% filter(year == 2022), REML = FALSE,
          control = lmerControl(optimizer ="Nelder_Mead"))
AIC(m3_mtqE)
```

```
## [1] 742.7664
```

```
AIC(m3_mtqE_full)
```

```
## [1] 652.6279
```

**Diagnostics**

Diagnostic plots for the tqE response are provided (small model followed by full model for models 1 through 3). Deviations from modeling assumptions are mild with most noticeable deviations occurring in the tails of the distributions for residuals.

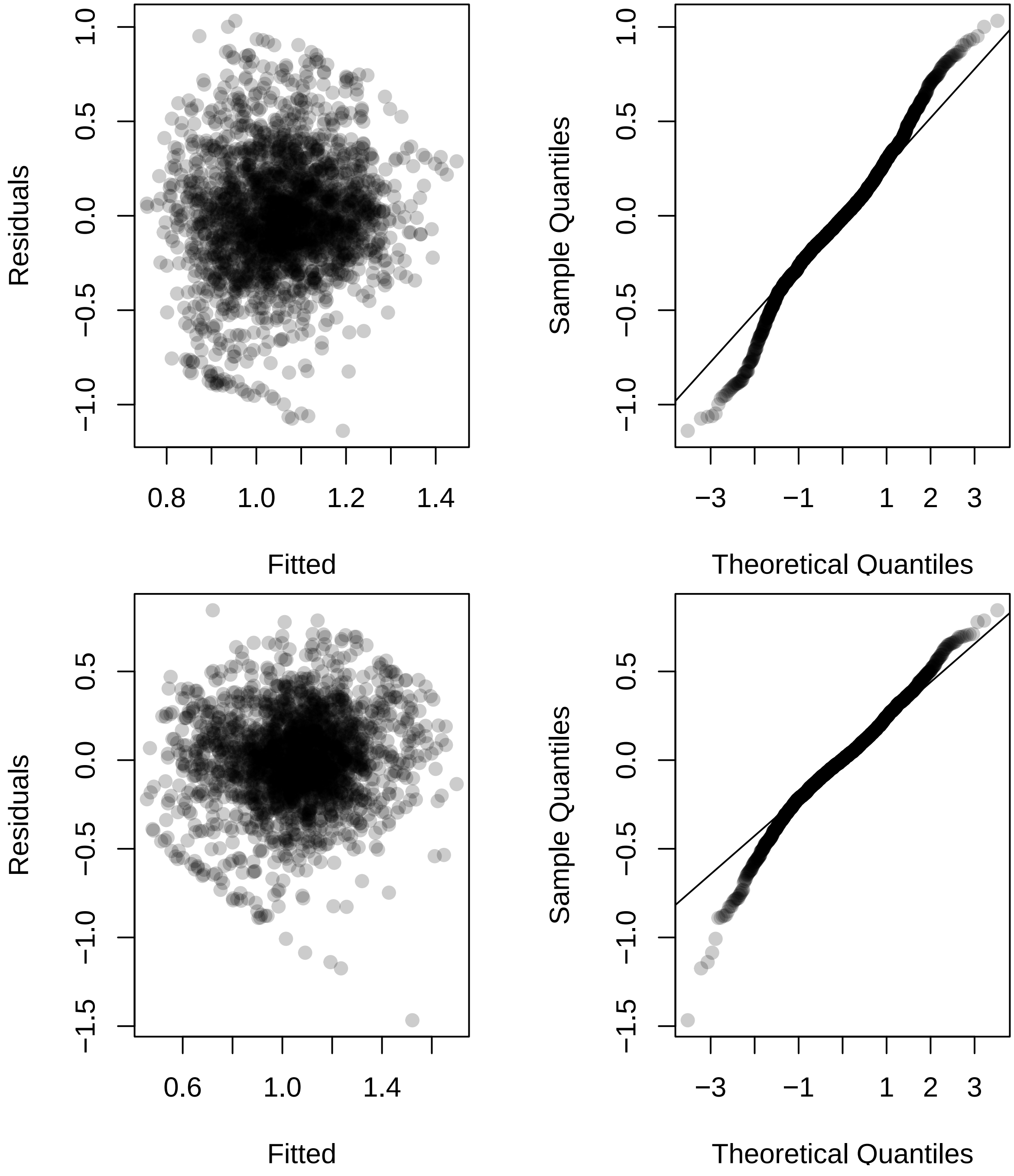

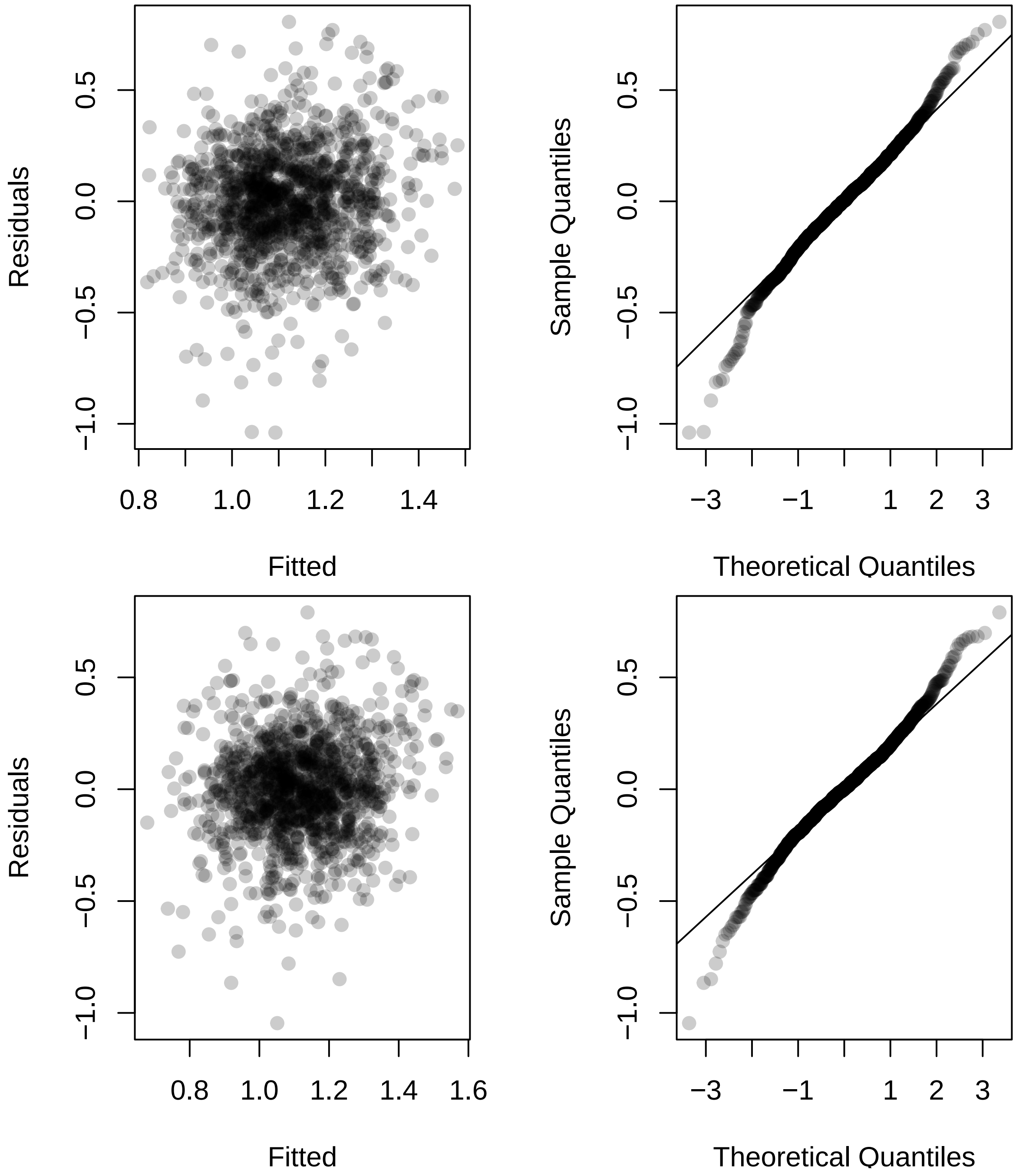
Residuals
Fitted
Sample Quantiles
Theoretical Quantiles
Residuals
Fitted
Sample Quantiles
Theoretical Quantiles

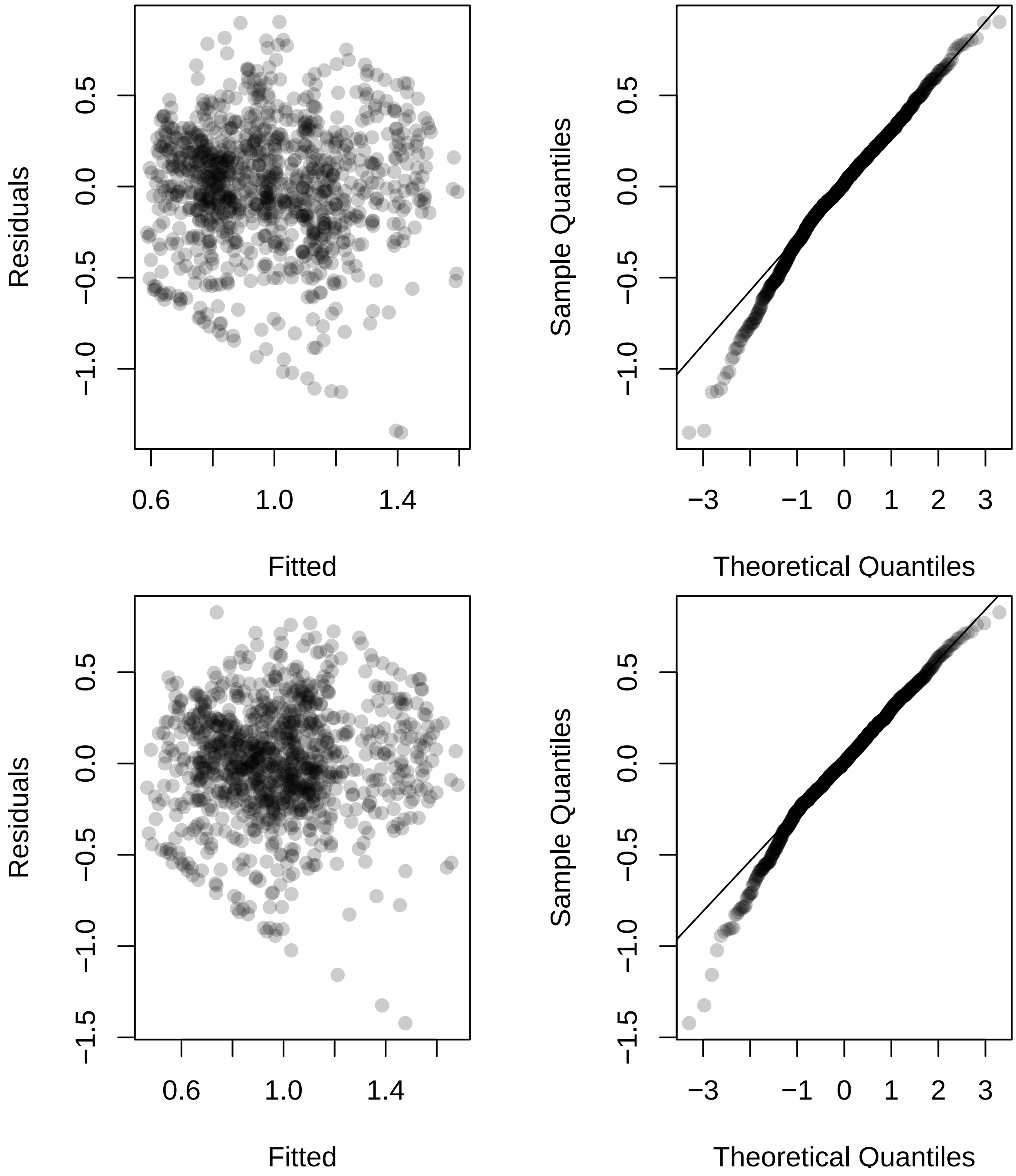
Residuals
Fitted
0.6
1.0
1.4
0.5
0.0
−0.5
−1.0
Sample Quantiles
Theoretical Quantiles
−3
−1
0
1
2
3
Residuals
Fitted
0.6
1.0
1.4
0.5
0.0
−0.5
−1.0
−1.5
Sample Quantiles
Theoretical Quantiles
−3
−1
0
1
2
3
0.5
0.0
−0.5
−1.0
−1.5

##### Investigate genotypes

We now perform an AIC based procedure to find genotypes that differ from the RC reference level.

```
## AIC for each ID variable from full tqE fixed-effects model
M = model.matrix(mtqE ~ ID + Date_num + I(Date_num^2), data = dat5)
M_full = model.matrix(mtqE ~ ID + Date_num + I(Date_num^2) +
                      Ta + VPD + Fsd + Precip + Precip_cum +
                      Ta_7day + VPD_7day + Fsd_7day + Precip_7day,
                 data = dat5)

M2021 = model.matrix(mtqE ~ ID + Date_num + I(Date_num^2),
                 data = dat5 %>% filter(year == 2021))
M2021_full = model.matrix(mtqE ~ ID + Date_num + I(Date_num^2) +
                      Ta + VPD + Fsd + Precip + Precip_cum +
                      Ta_7day + VPD_7day + Fsd_7day + Precip_7day,
                 data = dat5 %>% filter(year == 2021))

M2022 = model.matrix(mtqE ~ ID + Date_num + I(Date_num^2),
                 data = dat5 %>% filter(year == 2022))
M2022_full = model.matrix(mtqE ~ ID + Date_num + I(Date_num^2) +
                            Ta + VPD + Fsd + Precip + Precip_cum +
                            Ta_7day + VPD_7day + Fsd_7day + Precip_7day,
                 data = dat5 %>% filter(year == 2022))

ncores = detectCores() - 2
AIC_IDs_mtqE = do.call(rbind, mclapply(
  grep("IDNA", colnames(M)), function(j){
    M1 = M[, -j]
    foo = lmer(mtqE ~ -1 + M1 + (1|plot_number_year),
                data = dat5, REML = FALSE)
    M1_full = M_full[, -j]
    foo_full = lmer(mtqE ~ -1 + M1_full + (1|plot_number_year),
                data = dat5, REML = FALSE)
    M12021 = M2021[, -j]
    bar = lmer(mtqE ~ -1 + M12021 + (1|plot_number),
                data = dat5 %>% filter(year == 2021), REML = FALSE)
    M12021_full = M2021_full[, -j]
    bar_full = lmer(mtqE ~ -1 + M12021_full + (1|plot_number),
                data = dat5 %>% filter(year == 2021), REML = FALSE)
    M12022 = M2022[, -j]
    baz = lmer(mtqE ~ -1 + M12022 + (1|plot_number),
                data = dat5 %>% filter(year == 2022), REML = FALSE)
    M12022_full = M2022_full[, -j]
    baz_full = lmer(mtqE ~ -1 + M12022_full + (1|plot_number),
                data = dat5 %>% filter(year == 2022), REML = FALSE)

    c(AIC(m1_mtqE) - AIC(foo),
      AIC(m1_mtqE_full) - AIC(foo_full),
      AIC(m2_mtqE) - AIC(bar),
      AIC(m2_mtqE_full) - AIC(bar_full),
      AIC(m3_mtqE) - AIC(baz),
      AIC(m3_mtqE_full) - AIC(baz_full))
  }, mc.cores = ncores))
```

```r
rownames(AIC_IDs_mtqE) = colnames(M)[grep("IDNA", colnames(M))]
```

Negative values indicate that the genotype in question provides better model fit when differentiated from the RC reference level.

```r
round(AIC_IDs_mtqE, 3)
```

```
##            [,1]    [,2]    [,3]    [,4]   [,5]   [,6]
## IDNAM10  0.991   1.308   1.955   1.968  0.851  0.041
## IDNAM11  1.904   1.554   1.912   1.909  2.193  1.443
## IDNAM12 -9.006 -11.594 -13.206 -13.756 -0.234 -1.228
## IDNAM13 -1.578  -0.693   1.393   1.409 -0.185 -0.896
## IDNAM14  1.706   1.853   1.894   1.889  1.548  1.196
## IDNAM15  1.855   1.871   0.780   0.834  2.417  1.831
## IDNAM17  0.118  -0.652  -1.742  -1.975  2.298  1.619
## IDNAM18  0.683   1.114   1.984   1.971  0.210  0.054
## IDNAM2  -2.042  -1.713  -0.515  -0.413  0.628 -0.119
## IDNAM22  0.504  -0.547  -4.119  -4.278  2.581  1.994
## IDNAM23  1.766   1.719  -7.957  -8.071 -1.113 -2.147
## IDNAM24  1.622   1.427   1.723   1.695  2.199  1.590
## IDNAM25 -0.126  -0.014  -4.438  -3.643  2.583  2.000
## IDNAM26  0.405   0.510  -1.760  -1.407  2.512  1.978
## IDNAM27  1.840   1.879   1.909   1.907  2.527  1.849
## IDNAM28  1.885   1.967   1.639   1.668  2.006  1.438
## IDNAM29  1.609   1.318   0.956   1.138  2.503  1.833
## IDNAM3  -1.697  -3.615  -1.644  -1.663  0.094 -0.843
## IDNAM30  1.202   1.814   1.765   1.761  2.506  1.983
## IDNAM31  1.747   1.981   1.786   1.763  2.464  1.882
## IDNAM32  0.976   0.558   1.751   1.781 -1.602 -2.589
## IDNAM33  1.867   1.329   1.459   1.720  2.501  1.753
## IDNAM34  1.166   0.884  -2.029  -2.076  2.469  1.872
## IDNAM36 -1.049   0.026  -1.834  -1.879  2.538  1.953
## IDNAM37 -1.101  -1.884  -5.902  -5.409  2.463  1.748
## IDNAM38  1.995   1.900   1.060   0.934  2.397  1.830
## IDNAM39  1.186   0.831  -1.914  -1.487  2.563  1.998
## IDNAM4   1.114   1.791  -0.008  -0.336  1.990  1.281
## IDNAM40  1.013   1.648   1.994   1.965  2.008  1.566
## IDNAM41  1.178   1.340   2.000   1.999  1.412  0.863
## IDNAM42  1.666   1.886   1.979   1.975  2.139  1.598
## IDNAM46  1.724   1.636   0.438   0.542  2.580  1.971
## IDNAM48 -1.540  -0.928   1.162   1.122  0.005 -0.595
## IDNAM5   1.205   1.879   1.997   1.999  2.239  1.841
## IDNAM50  1.598   1.673  -0.145  -0.040 -1.800 -2.258
## IDNAM54  0.610   0.696   1.868   1.809  1.113  0.605
## IDNAM6  -0.722   0.071   1.903   1.915 -0.445 -0.958
## IDNAM64  0.775   0.919  -2.849  -2.717  2.435  1.858
## IDNAM8   0.212   0.586   0.192  -0.105  2.476  1.890
## IDNAM9   1.939   2.000   0.771   0.841  1.640  1.249
```

```r
sign(AIC_IDs_mtqE < 0)
```

```
##         [,1] [,2] [,3] [,4] [,5] [,6]
## IDNAM10    0    0    0    0    0    0
## IDNAM11    0    0    0    0    0    0
## IDNAM12    1    1    1    1    1    1
```

```
## IDNAM13    1    1    0    0    1    1
## IDNAM14    0    0    0    0    0    0
## IDNAM15    0    0    0    0    0    0
## IDNAM17    0    1    1    1    0    0
## IDNAM18    0    0    0    0    0    0
## IDNAM2     1    1    1    1    0    1
## IDNAM22    0    1    1    1    0    0
## IDNAM23    0    0    1    1    1    1
## IDNAM24    0    0    0    0    0    0
## IDNAM25    1    1    1    1    0    0
## IDNAM26    0    0    1    1    0    0
## IDNAM27    0    0    0    0    0    0
## IDNAM28    0    0    0    0    0    0
## IDNAM29    0    0    0    0    0    0
## IDNAM3     1    1    1    1    0    1
## IDNAM30    0    0    0    0    0    0
## IDNAM31    0    0    0    0    0    0
## IDNAM32    0    0    0    0    1    1
## IDNAM33    0    0    0    0    0    0
## IDNAM34    0    0    1    1    0    0
## IDNAM36    1    0    1    1    0    0
## IDNAM37    1    1    1    1    0    0
## IDNAM38    0    0    0    0    0    0
## IDNAM39    0    0    1    1    0    0
## IDNAM4     0    0    1    1    0    0
## IDNAM40    0    0    0    0    0    0
## IDNAM41    0    0    0    0    0    0
## IDNAM42    0    0    0    0    0    0
## IDNAM46    0    0    0    0    0    0
## IDNAM48    1    1    0    0    0    1
## IDNAM5     0    0    0    0    0    0
## IDNAM50    0    0    1    1    1    1
## IDNAM54    0    0    0    0    0    0
## IDNAM6     1    0    0    0    1    1
## IDNAM64    0    0    1    1    0    0
## IDNAM8     0    0    0    1    0    0
## IDNAM9     0    0    0    0    0    0
```

We now investigate the specific coefficient effects from our fitted models. Specific interest is in determining the sign of the effect. Here are the effect estimates from each fitted model:

```
coefs_mtqE = round(cbind(summary(m1_mtqE)$coefficients[2:41, 1],
                   summary(m1_mtqE_full)$coefficients[2:41, 1],
                   summary(m2_mtqE)$coefficients[2:41, 1],
                   summary(m2_mtqE_full)$coefficients[2:41, 1],
                   summary(m3_mtqE)$coefficients[2:41, 1],
                   summary(m3_mtqE_full)$coefficients[2:41, 1]), 3)
coefs_mtqE
```

```
##           [,1]   [,2]   [,3]   [,4]   [,5]   [,6]
## IDNAM10 -0.067 -0.056  0.016  0.014 -0.132 -0.139
## IDNAM11  0.020  0.044  0.022  0.023  0.061  0.071
## IDNAM12  0.221  0.246  0.303  0.308  0.165  0.175
## IDNAM13 -0.125 -0.108 -0.060 -0.059 -0.158 -0.162
## IDNAM14 -0.035 -0.025  0.025  0.025 -0.099 -0.086
## IDNAM15  0.025  0.023  0.084  0.082 -0.036 -0.038
## IDNAM17  0.091  0.108  0.150  0.154  0.053  0.059
## IDNAM18 -0.077 -0.063  0.010  0.013 -0.150 -0.138
## IDNAM2  -0.129 -0.124 -0.121 -0.118 -0.128 -0.134
## IDNAM22  0.081  0.106  0.193  0.194 -0.004  0.007
## IDNAM23  0.032  0.035  0.244  0.244 -0.182 -0.194
## IDNAM24  0.040  0.049  0.040  0.042  0.060  0.060
## IDNAM25  0.096  0.094  0.195  0.182  0.002  0.000
## IDNAM26  0.083  0.080  0.151  0.143  0.026  0.014
## IDNAM27 -0.027  0.023  0.023  0.023  0.026  0.039
## IDNAM28 -0.022 -0.012  0.045  0.043 -0.073 -0.073
## IDNAM29  0.041  0.055  0.078  0.070  0.029  0.040
## IDNAM3   0.127  0.157  0.149  0.148  0.152  0.161
## IDNAM30 -0.058 -0.028 -0.037 -0.037 -0.026 -0.013
## IDNAM31 -0.033 -0.009 -0.035 -0.037  0.036  0.033
## IDNAM32 -0.066 -0.078  0.038  0.035 -0.194 -0.204
## IDNAM33  0.027  0.060  0.070  0.050  0.031  0.049
## IDNAM34  0.063  0.073  0.154  0.154 -0.036 -0.039
## IDNAM36 -0.114 -0.092 -0.149 -0.149 -0.019 -0.021
## IDNAM37  0.116  0.130  0.217  0.209  0.037  0.048
## IDNAM38  0.005  0.021  0.074  0.079 -0.039 -0.039
## IDNAM39  0.059  0.071  0.151  0.142 -0.011 -0.004
## IDNAM4  -0.063 -0.030 -0.108 -0.117  0.078  0.084
## IDNAM40 -0.065 -0.039 -0.006 -0.014 -0.073 -0.064
## IDNAM41 -0.060 -0.054  0.000 -0.002 -0.102 -0.104
## IDNAM42 -0.038 -0.022  0.011  0.012 -0.062 -0.060
## IDNAM46  0.034  0.040  0.095  0.091 -0.005 -0.016
## IDNAM48 -0.124 -0.112 -0.070 -0.071 -0.154 -0.155
## IDNAM5  -0.059 -0.023  0.004 -0.003 -0.056 -0.039
## IDNAM50 -0.042 -0.037  0.113  0.109 -0.201 -0.197
## IDNAM54 -0.077 -0.075 -0.028 -0.033 -0.112 -0.112
## IDNAM6  -0.108 -0.091 -0.024 -0.022 -0.165 -0.164
## IDNAM64  0.072  0.068  0.170  0.167 -0.035 -0.035
## IDNAM8  -0.088 -0.079 -0.103 -0.111 -0.030 -0.032
## IDNAM9  -0.017  0.000  0.086  0.083 -0.095 -0.088
```

Here are the corresponding signs:

```r
sign(coefs_mtqE > 0)
```

```
##          [,1] [,2] [,3] [,4] [,5] [,6]
## IDNAM10     0    0    1    1    0    0
## IDNAM11     1    1    1    1    1    1
## IDNAM12     1    1    1    1    1    1
## IDNAM13     0    0    0    0    0    0
## IDNAM14     0    0    1    1    0    0
## IDNAM15     1    1    1    1    0    0
## IDNAM17     1    1    1    1    1    1
## IDNAM18     0    0    1    1    0    0
## IDNAM2      0    0    0    0    0    0
## IDNAM22     1    1    1    1    0    1
## IDNAM23     1    1    1    1    0    0
## IDNAM24     1    1    1    1    1    1
## IDNAM25     1    1    1    1    1    0
## IDNAM26     1    1    1    1    1    1
## IDNAM27     0    1    1    1    1    1
## IDNAM28     0    0    1    1    0    0
## IDNAM29     1    1    1    1    1    1
## IDNAM3      1    1    1    1    1    1
## IDNAM30     0    0    0    0    0    0
## IDNAM31     0    0    0    0    1    1
## IDNAM32     0    0    1    1    0    0
## IDNAM33     1    1    1    1    1    1
## IDNAM34     1    1    1    1    0    0
## IDNAM36     0    0    0    0    0    0
## IDNAM37     1    1    1    1    1    1
## IDNAM38     1    1    1    1    0    0
## IDNAM39     1    1    1    1    0    0
## IDNAM4      0    0    0    0    1    1
## IDNAM40     0    0    0    0    0    0
## IDNAM41     0    0    0    0    0    0
## IDNAM42     0    0    1    1    0    0
## IDNAM46     1    1    1    1    0    0
## IDNAM48     0    0    0    0    0    0
## IDNAM5      0    0    1    0    0    0
## IDNAM50     0    0    1    1    0    0
## IDNAM54     0    0    0    0    0    0
## IDNAM6      0    0    0    0    0    0
## IDNAM64     1    1    1    1    0    0
## IDNAM8      0    0    0    0    0    0
## IDNAM9      0    0    1    1    0    0
```

Below are signs that indicate which genotypes yield better model fit when their effect is differentiated from the RC reference level. A 1 indicates a positive effect, a -1 indicates a negative effect, and a 0 indicates that the genotype was not determined to provide better model when differentiated from the RC reference level.

```r
inference_mtqE = sign(coefs_mtqE) * sign(AIC_IDs_mtqE < 0)
inference_mtqE
```

```
##          [,1] [,2] [,3] [,4] [,5] [,6]
## IDNAM10     0    0    0    0    0    0
## IDNAM11     0    0    0    0    0    0
## IDNAM12     1    1    1    1    1    1
## IDNAM13    -1   -1    0    0   -1   -1
## IDNAM14     0    0    0    0    0    0
## IDNAM15     0    0    0    0    0    0
## IDNAM17     0    1    1    1    0    0
## IDNAM18     0    0    0    0    0    0
## IDNAM2     -1   -1   -1   -1    0   -1
## IDNAM22     0    1    1    1    0    0
## IDNAM23     0    0    1    1   -1   -1
## IDNAM24     0    0    0    0    0    0
## IDNAM25     1    1    1    1    0    0
## IDNAM26     0    0    1    1    0    0
## IDNAM27     0    0    0    0    0    0
## IDNAM28     0    0    0    0    0    0
## IDNAM29     0    0    0    0    0    0
## IDNAM3      1    1    1    1    0    1
## IDNAM30     0    0    0    0    0    0
## IDNAM31     0    0    0    0    0    0
## IDNAM32     0    0    0    0   -1   -1
## IDNAM33     0    0    0    0    0    0
## IDNAM34     0    0    1    1    0    0
## IDNAM36    -1    0   -1   -1    0    0
## IDNAM37     1    1    1    1    0    0
## IDNAM38     0    0    0    0    0    0
## IDNAM39     0    0    1    1    0    0
## IDNAM4      0    0   -1   -1    0    0
## IDNAM40     0    0    0    0    0    0
## IDNAM41     0    0    0    0    0    0
## IDNAM42     0    0    0    0    0    0
## IDNAM46     0    0    0    0    0    0
## IDNAM48    -1   -1    0    0    0   -1
## IDNAM5      0    0    0    0    0    0
## IDNAM50     0    0    1    1   -1   -1
## IDNAM54     0    0    0    0    0    0
## IDNAM6     -1    0    0    0   -1   -1
## IDNAM64     0    0    1    1    0    0
## IDNAM8      0    0    0   -1    0    0
## IDNAM9      0    0    0    0    0    0
```

**Results**

Here are the effect size estimates for genotypes that were to determined to differentiate from the RC reference level in all considered models and had the same estimated sign across all considered models.

```
rownames(coefs_mtqE)[abs(rowSums(inference_mtqE)) == 6]
```

```
## [1] "IDNAM12"
```

```
coefs_mtqE[abs(rowSums(inference_mtqE)) == 6, ]
```

```
## [1] 0.221 0.246 0.303 0.308 0.165 0.175
```

##### tqM response

```
m1_mtqM = lmer(mtqM ~ ID + Date_num + I(Date_num^2) +
               (1|plot_number_year),
           data = dat5, REML = FALSE, control = lmerControl(optimizer ="Nelder_Mead"))
m1_mtqM_full = lmer(mtqM ~ ID + Date_num + I(Date_num^2) +
                      Ta + VPD + Fsd + Precip + Precip_cum +
                      Ta_7day + VPD_7day + Fsd_7day + Precip_7day +
                      (1|plot_number_year),
           data = dat5, REML = FALSE, control = lmerControl(optimizer ="Nelder_Mead"))
AIC(m1_mtqM)
```

```
## [1] 16068.24
```

```
AIC(m1_mtqM_full)
```

```
## [1] 15773.49
```

```
m2_mtqM = lmer(mtqM ~ ID + Date_num + I(Date_num^2) +
               (1|plot_number),
           data = dat5 %>% filter(year == 2021), REML = FALSE,
           control = lmerControl(optimizer ="Nelder_Mead"))
m2_mtqM_full = lmer(mtqM ~ ID + Date_num + I(Date_num^2) +
                      Ta + VPD + Fsd + Precip + Precip_cum +
                      Ta_7day + VPD_7day + Fsd_7day + Precip_7day +
                      (1|plot_number),
           data = dat5 %>% filter(year == 2021), REML = FALSE,
           control = lmerControl(optimizer ="Nelder_Mead"))
AIC(m2_mtqM)
```

```
## [1] 8783.907
```

```
AIC(m2_mtqM_full)
```

```
## [1] 8533.791
```

```
m3_mtqM = lmer(mtqM ~ ID + Date_num + I(Date_num^2) +
               (1|plot_number),
           data = dat5 %>% filter(year == 2022), REML = FALSE,
           control = lmerControl(optimizer ="Nelder_Mead"))
m3_mtqM_full = lmer(mtqM ~ ID + Date_num + I(Date_num^2) +
                      Ta + VPD + Fsd + Precip + Precip_cum +
                      Ta_7day + VPD_7day + Fsd_7day + Precip_7day +
                      (1|plot_number),
           data = dat5 %>% filter(year == 2022), REML = FALSE,
           control = lmerControl(optimizer ="Nelder_Mead"))
AIC(m3_mtqM)
```

```
## [1] 7263.609
```

```
AIC(m3_mtqM_full)
```

```
## [1] 7203.446
```

**Diagnostics**

Diagnostic plots for the tqM response are provided (small model followed by full model for models 1 through 3). Modeling assumptions appear to be violated.

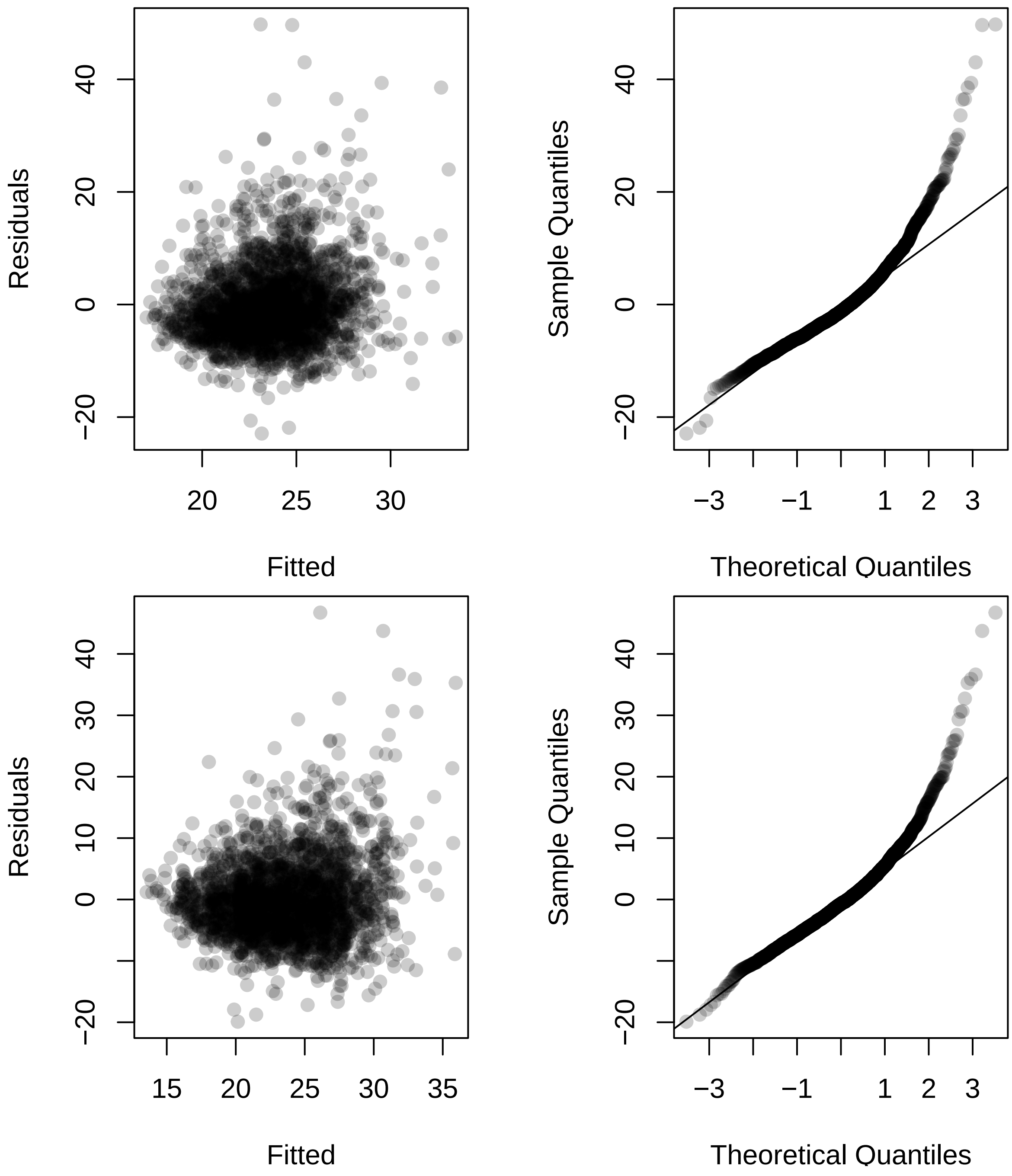

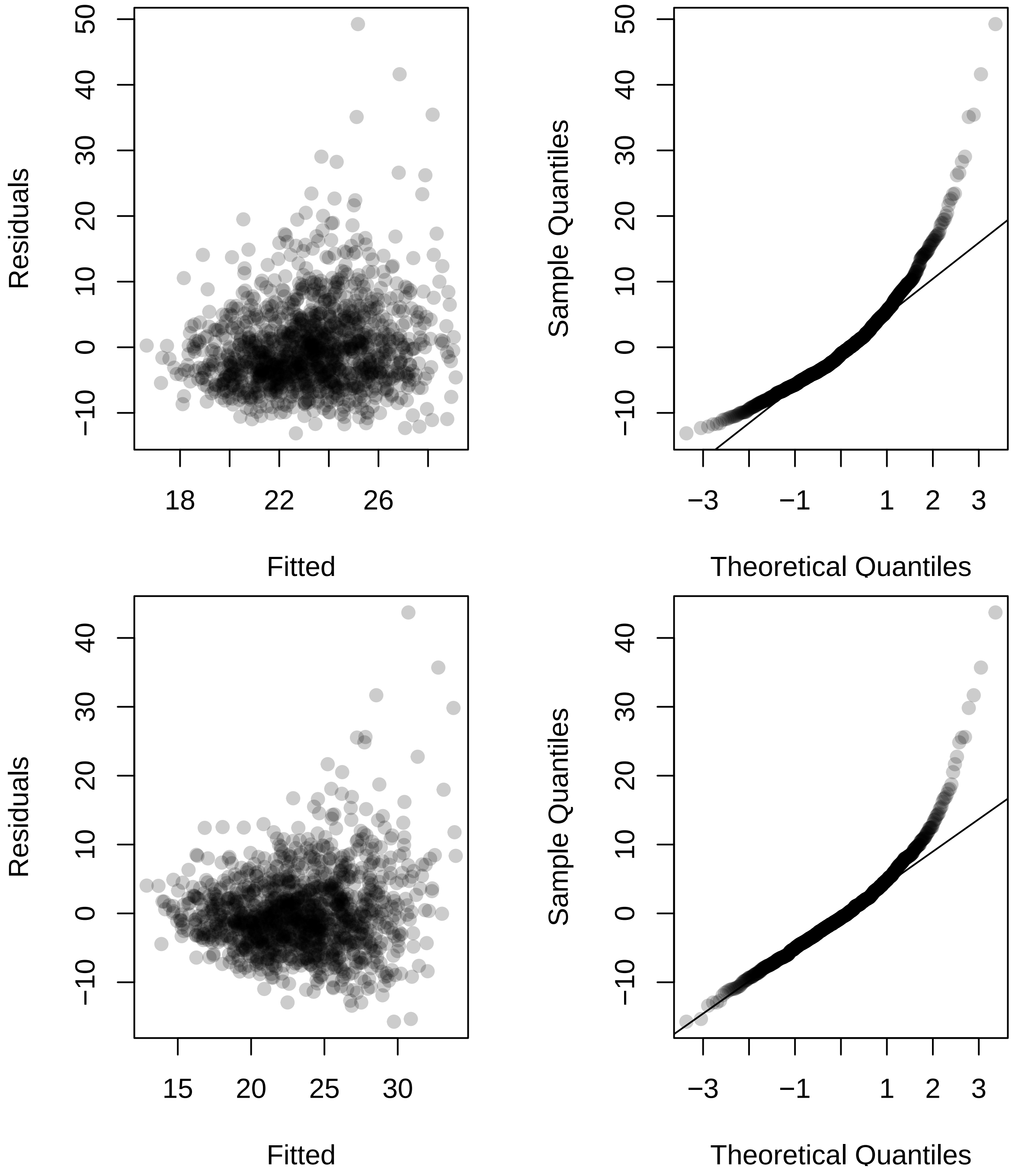
Residuals
Fitted
Sample Quantiles
Theoretical Quantiles
Residuals
Fitted
Sample Quantiles
Theoretical Quantiles

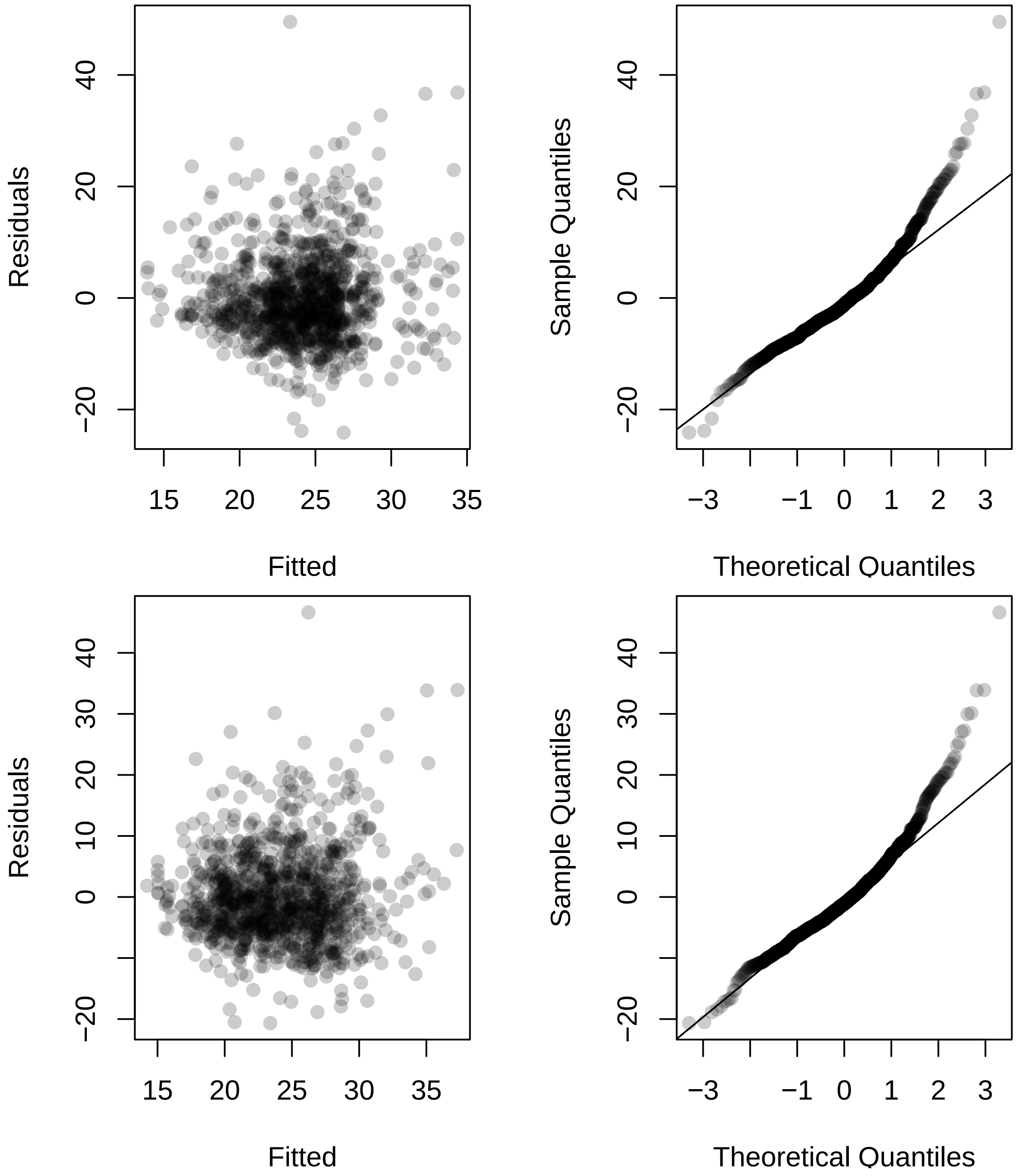
Residuals
Fitted
Sample Quantiles
Theoretical Quantiles
−20 0 20 40
15 20 25 30 35
−3 −1 0 1 2 3
−20 0 10 20 30 40

##### Log transform

We now consider a log response transformation to alleviate modeling problems. This transformation seems to alleviate serious problems with minor modeling assumption deviations remaining.

```
m1_mtqM_log = lmer(log(mtqM) ~ ID + Date_num + I(Date_num^2) +
               (1|plot_number_year),
           data = dat5, REML = FALSE, control = lmerControl(optimizer ="Nelder_Mead"))
m1_mtqM_log_full = lmer(log(mtqM) ~ ID + Date_num + I(Date_num^2) +
                      Ta + VPD + Fsd + Precip + Precip_cum +
                      Ta_7day + VPD_7day + Fsd_7day + Precip_7day +
                      (1|plot_number_year),
           data = dat5, REML = FALSE, control = lmerControl(optimizer ="Nelder_Mead"))
AIC(m1_mtqM_log)
```

```
## [1] 1386.456
```

```
AIC(m1_mtqM_log_full)
```

```
## [1] 1110.765
```

```
m2_mtqM_log = lmer(log(mtqM) ~ ID + Date_num + I(Date_num^2) +
               (1|plot_number),
           data = dat5 %>% filter(year == 2021), REML = FALSE,
           control = lmerControl(optimizer ="Nelder_Mead"))
m2_mtqM_log_full = lmer(log(mtqM) ~ ID + Date_num + I(Date_num^2) +
                      Ta + VPD + Fsd + Precip + Precip_cum +
                      Ta_7day + VPD_7day + Fsd_7day + Precip_7day +
                      (1|plot_number),
           data = dat5 %>% filter(year == 2021), REML = FALSE,
           control = lmerControl(optimizer ="Nelder_Mead"))
AIC(m2_mtqM_log)
```

```
## [1] 412.6305
```

```
AIC(m2_mtqM_log_full)
```

```
## [1] 155.8909
```

```
m3_mtqM_log = lmer(log(mtqM) ~ ID + Date_num + I(Date_num^2) +
               (1|plot_number),
           data = dat5 %>% filter(year == 2022), REML = FALSE,
           control = lmerControl(optimizer ="Nelder_Mead"))
m3_mtqM_log_full = lmer(log(mtqM) ~ ID + Date_num + I(Date_num^2) +
                      Ta + VPD + Fsd + Precip + Precip_cum +
                      Ta_7day + VPD_7day + Fsd_7day + Precip_7day +
                      (1|plot_number),
           data = dat5 %>% filter(year == 2022), REML = FALSE,
           control = lmerControl(optimizer ="Nelder_Mead"))
AIC(m3_mtqM_log)
```

```
## [1] 900.6977
```

```
AIC(m3_mtqM_log_full)
```

```
## [1] 840.5858
```

**Diagnostics**

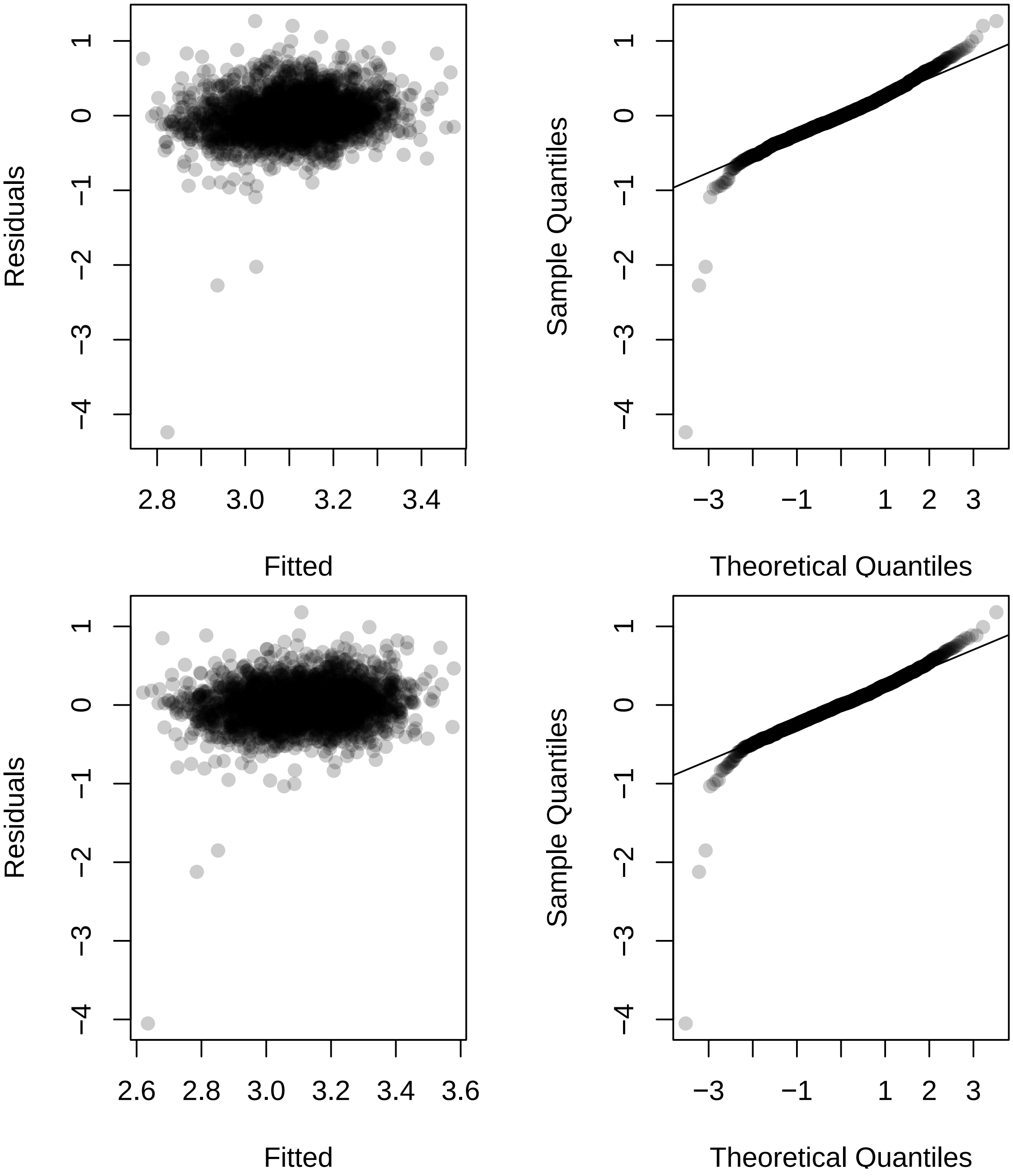

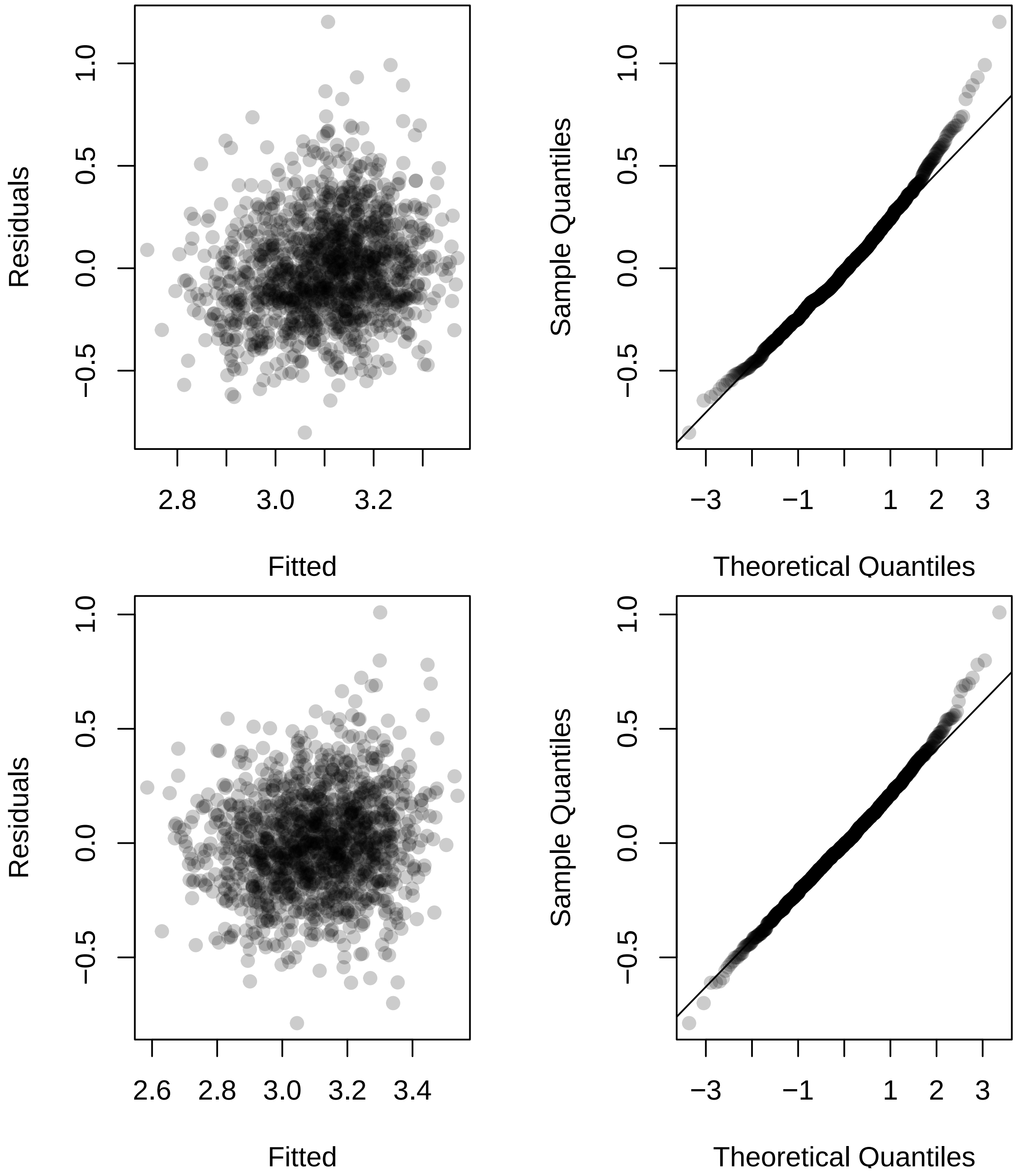
Residuals
Fitted
Sample Quantiles
Theoretical Quantiles
Residuals
Fitted
Sample Quantiles
Theoretical Quantiles

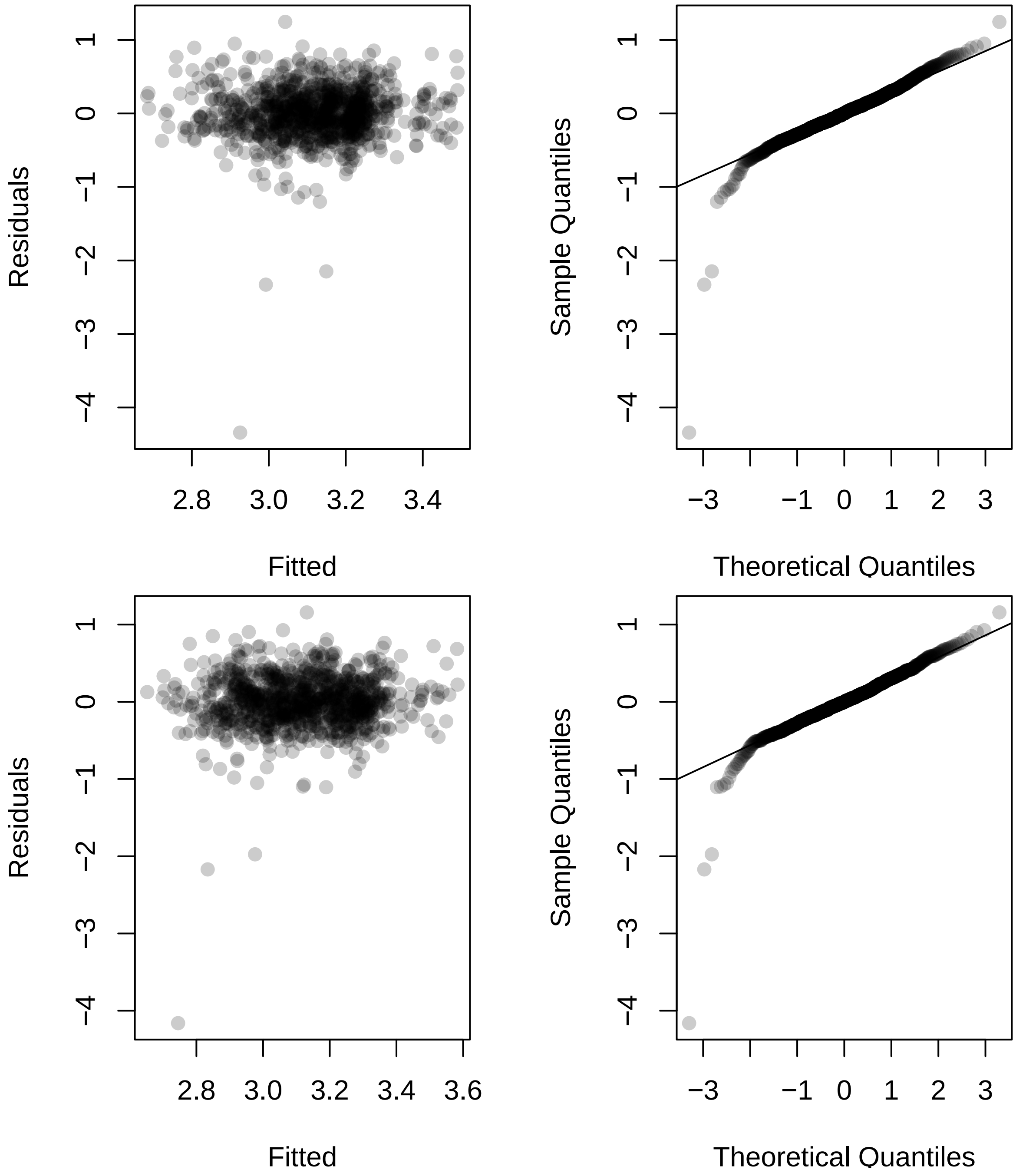
Residuals
1
0
−1
−2
−3
−4
2.8
3.0
3.2
3.4
Fitted
Sample Quantiles
Theoretical Quantiles
−3
−1
0
1
2
3
Residuals
2.8
3.0
3.2
3.4
3.6
Fitted
Sample Quantiles
Theoretical Quantiles

**Investigate genotypes**

We now perform an AIC based procedure to find genotypes that differ from the RC reference level.

```
M = model.matrix(mtqM ~ ID + Date_num + I(Date_num^2), data = dat5)
M_full = model.matrix(mtqM ~ ID + Date_num + I(Date_num^2) +
                      Ta + VPD + Fsd + Precip + Precip_cum +
                      Ta_7day + VPD_7day + Fsd_7day + Precip_7day,
                 data = dat5)

M2021 = model.matrix(mtqM ~ ID + Date_num + I(Date_num^2),
                 data = dat5 %>% filter(year == 2021))
M2021_full = model.matrix(mtqM ~ ID + Date_num + I(Date_num^2) +
                      Ta + VPD + Fsd + Precip + Precip_cum +
                      Ta_7day + VPD_7day + Fsd_7day + Precip_7day,
                 data = dat5 %>% filter(year == 2021))

M2022 = model.matrix(mtqM ~ ID + Date_num + I(Date_num^2),
                 data = dat5 %>% filter(year == 2022))
M2022_full = model.matrix(mtqM ~ ID + Date_num + I(Date_num^2) +
                            Ta + VPD + Fsd + Precip + Precip_cum +
                            Ta_7day + VPD_7day + Fsd_7day + Precip_7day,
                 data = dat5 %>% filter(year == 2022))

ncores = detectCores() - 2
AIC_IDs_mtqM = do.call(rbind, mclapply(
  grep("IDNA", colnames(M)), function(j){
    M1 = M[, -j]
    foo = lmer(log(mtqM) ~ -1 + M1 + (1|plot_number_year),
                data = dat5, REML = FALSE)
    M1_full = M_full[, -j]
    foo_full = lmer(log(mtqM) ~ -1 + M1_full + (1|plot_number_year),
                data = dat5, REML = FALSE)
    M12021 = M2021[, -j]
    bar = lmer(log(mtqM) ~ -1 + M12021 + (1|plot_number),
                data = dat5 %>% filter(year == 2021), REML = FALSE)
    M12021_full = M2021_full[, -j]
    bar_full = lmer(log(mtqM) ~ -1 + M12021_full + (1|plot_number),
                data = dat5 %>% filter(year == 2021), REML = FALSE)
    M12022 = M2022[, -j]
    baz = lmer(log(mtqM) ~ -1 + M12022 + (1|plot_number),
                data = dat5 %>% filter(year == 2022), REML = FALSE)
    M12022_full = M2022_full[, -j]
    baz_full = lmer(log(mtqM) ~ -1 + M12022_full + (1|plot_number),
                data = dat5 %>% filter(year == 2022), REML = FALSE)

    c(AIC(m1_mtqM_log) - AIC(foo),
      AIC(m1_mtqM_log_full) - AIC(foo_full),
      AIC(m2_mtqM_log) - AIC(bar),
      AIC(m2_mtqM_log_full) - AIC(bar_full),
      AIC(m3_mtqM_log) - AIC(baz),
      AIC(m3_mtqM_log_full) - AIC(baz_full))
  }, mc.cores = ncores))
rownames(AIC_IDs_mtqM) = colnames(M)[grep("IDNA", colnames(M))]
```

Negative values indicate that the genotype in question provides better model fit when differentiated from the RC reference level.

```r
round(AIC_IDs_mtqM, 3)
```

```
##              [,1]    [,2]   [,3]   [,4]    [,5]    [,6]
## IDNAM10  -0.809  -1.120  1.519  1.415  -0.626  -0.955
## IDNAM11   1.839   1.812  1.880  1.922   1.163   1.163
## IDNAM12  -0.962  -0.822  0.828  0.775  -0.049   0.049
## IDNAM13   1.719   1.750  1.212  1.244   1.992   1.988
## IDNAM14   1.478   1.400  1.623  1.663   1.793   1.688
## IDNAM15   0.110  -0.120  1.889  1.852  -0.906  -0.988
## IDNAM17 -13.513 -13.402 -1.581 -1.428 -13.228 -13.253
## IDNAM18  -3.504  -3.916 -2.499 -2.540  -0.157  -0.500
## IDNAM2    1.709   1.765  1.933  1.959   1.615   1.644
## IDNAM22  -6.539  -6.321 -1.623 -1.743  -3.996  -3.480
## IDNAM23   1.794   1.645  1.756  1.603   1.942   1.933
## IDNAM24   1.093   1.004  1.794  1.729   1.180   1.125
## IDNAM25   1.798   1.747  1.948  1.993   1.290   1.314
## IDNAM26   1.778   1.838  1.766  1.805   0.773   1.130
## IDNAM27  -1.115  -0.969  1.865  1.820  -2.985  -2.618
## IDNAM28  -1.651  -1.554  0.944  0.991  -1.293  -1.163
## IDNAM29  -0.889  -0.879  1.397  1.316  -0.938  -0.884
## IDNAM3    1.274   1.484  1.957  1.990   0.988   1.141
## IDNAM30   1.826   1.866  1.994  1.994   1.579   1.632
## IDNAM31  -1.797  -1.965  1.563  1.466  -2.622  -2.781
## IDNAM32   1.996   1.999  1.968  1.965   1.941   1.941
## IDNAM33  -0.082  -0.822  1.386  0.388   0.074   0.036
## IDNAM34   1.408   1.034  1.943  1.875   1.275   0.929
## IDNAM36   0.390   0.417  1.985  1.988  -0.987  -0.957
## IDNAM37  -2.275  -2.684 -2.629 -3.328   1.206   1.154
## IDNAM38  -3.587  -3.292 -1.051 -1.091  -1.102  -0.821
## IDNAM39  -0.864  -1.163  1.348  1.087  -0.952  -0.936
## IDNAM4    1.704   1.671  1.955  1.826   1.874   1.882
## IDNAM40   1.624   1.577  1.805  1.646   1.757   1.842
## IDNAM41  -0.171  -0.535  1.926  1.897  -1.776  -2.196
## IDNAM42   1.999   1.998  1.660  1.700   1.484   1.624
## IDNAM46  -3.335  -3.500  1.983  1.971  -8.853  -8.527
## IDNAM48   0.535   0.598 -0.618 -0.273   1.913   1.902
## IDNAM5    0.082  -0.189  1.352  1.366   0.095  -0.146
## IDNAM50   1.236   1.146  1.063  1.051   1.883   1.809
## IDNAM54  -3.477  -3.599 -1.774 -1.822  -0.682  -0.741
## IDNAM6    1.011   0.885 -0.330 -0.541   1.995   1.995
## IDNAM64   1.977   1.984  1.956  1.951   1.994   1.989
## IDNAM8   -1.873  -2.101 -0.035 -0.425  -0.274  -0.156
## IDNAM9    1.827   1.819 -0.117 -0.044   0.878   0.927
```

```r
sign(AIC_IDs_mtqM < 0)
```

```
##         [,1] [,2] [,3] [,4] [,5] [,6]
## IDNAM10    1    1    0    0    1    1
## IDNAM11    0    0    0    0    0    0
## IDNAM12    1    1    0    0    1    0
## IDNAM13    0    0    0    0    0    0
## IDNAM14    0    0    0    0    0    0
```

```
## IDNAM15    0    1    0    0    1    1
## IDNAM17    1    1    1    1    1    1
## IDNAM18    1    1    1    1    1    1
## IDNAM2     0    0    0    0    0    0
## IDNAM22    1    1    1    1    1    1
## IDNAM23    0    0    0    0    0    0
## IDNAM24    0    0    0    0    0    0
## IDNAM25    0    0    0    0    0    0
## IDNAM26    0    0    0    0    0    0
## IDNAM27    1    1    0    0    1    1
## IDNAM28    1    1    0    0    1    1
## IDNAM29    1    1    0    0    1    1
## IDNAM3     0    0    0    0    0    0
## IDNAM30    0    0    0    0    0    0
## IDNAM31    1    1    0    0    1    1
## IDNAM32    0    0    0    0    0    0
## IDNAM33    1    1    0    0    0    0
## IDNAM34    0    0    0    0    0    0
## IDNAM36    0    0    0    0    1    1
## IDNAM37    1    1    1    1    0    0
## IDNAM38    1    1    1    1    1    1
## IDNAM39    1    1    0    0    1    1
## IDNAM4     0    0    0    0    0    0
## IDNAM40    0    0    0    0    0    0
## IDNAM41    1    1    0    0    1    1
## IDNAM42    0    0    0    0    0    0
## IDNAM46    1    1    0    0    1    1
## IDNAM48    0    0    1    1    0    0
## IDNAM5     0    1    0    0    0    1
## IDNAM50    0    0    0    0    0    0
## IDNAM54    1    1    1    1    1    1
## IDNAM6     0    0    1    1    0    0
## IDNAM64    0    0    0    0    0    0
## IDNAM8     1    1    1    1    1    1
## IDNAM9     0    0    1    1    0    0
```

We now investigate the specific coefficient effects from our fitted models. Specific interest is in determining the sign of the effect. Here are the effect estimates from each fitted model:

```
coefs_mtqM = round(cbind(summary(m1_mtqM_log)$coefficients[2:41, 1],
                   summary(m1_mtqM_log_full)$coefficients[2:41, 1],
                   summary(m2_mtqM_log)$coefficients[2:41, 1],
                   summary(m2_mtqM_log_full)$coefficients[2:41, 1],
                   summary(m3_mtqM_log)$coefficients[2:41, 1],
                   summary(m3_mtqM_log_full)$coefficients[2:41, 1]), 3)
coefs_mtqM
```

```
##           [,1]   [,2]   [,3]   [,4]   [,5]   [,6]
## IDNAM10  0.121  0.127  0.062  0.068  0.173  0.182
## IDNAM11  0.028  0.031 -0.031 -0.025  0.094  0.093
## IDNAM12  0.123  0.119  0.096  0.098  0.150  0.145
## IDNAM13 -0.038 -0.035 -0.080 -0.078  0.009  0.011
## IDNAM14  0.051  0.054  0.054  0.051  0.047  0.057
## IDNAM15  0.096  0.101  0.030  0.034  0.170  0.172
## IDNAM17  0.284  0.281  0.170  0.166  0.409  0.407
## IDNAM18  0.170  0.175  0.191  0.191  0.158  0.168
## IDNAM2   0.037  0.033  0.023  0.018  0.061  0.059
## IDNAM22  0.210  0.206  0.171  0.173  0.256  0.243
## IDNAM23  0.032  0.042  0.044  0.056  0.025  0.026
## IDNAM24 -0.067 -0.070 -0.040 -0.046 -0.091 -0.093
## IDNAM25  0.032  0.036 -0.020 -0.008  0.087  0.085
## IDNAM26  0.034  0.028 -0.043 -0.039  0.114  0.095
## IDNAM27  0.127  0.123  0.033  0.037  0.244  0.232
## IDNAM28  0.136  0.133  0.091  0.088  0.191  0.185
## IDNAM29  0.122  0.121  0.069  0.073  0.182  0.179
## IDNAM3   0.061  0.051  0.019  0.009  0.103  0.094
## IDNAM30  0.029  0.026 -0.007 -0.007  0.067  0.062
## IDNAM31  0.139  0.141  0.059  0.065  0.226  0.228
## IDNAM32 -0.004 -0.002  0.016  0.016 -0.025 -0.025
## IDNAM33  0.114  0.131  0.086  0.137  0.147  0.147
## IDNAM34  0.057  0.072  0.021  0.031  0.100  0.120
## IDNAM36  0.090  0.088  0.011  0.010  0.178  0.176
## IDNAM37  0.148  0.153  0.192  0.206  0.092  0.095
## IDNAM38  0.169  0.163  0.156  0.157  0.181  0.172
## IDNAM39  0.120  0.125  0.071  0.084  0.177  0.176
## IDNAM4   0.039  0.041  0.019  0.037  0.038  0.036
## IDNAM40  0.044  0.046  0.039  0.053  0.052  0.041
## IDNAM41  0.105  0.113  0.024  0.028  0.205  0.214
## IDNAM42  0.002 -0.003 -0.052 -0.048  0.074  0.063
## IDNAM46  0.164  0.165  0.011  0.015  0.346  0.338
## IDNAM48 -0.086 -0.083 -0.144 -0.134 -0.031 -0.032
## IDNAM5  -0.099 -0.105 -0.071 -0.070 -0.146 -0.154
## IDNAM50 -0.062 -0.065 -0.086 -0.086 -0.035 -0.044
## IDNAM54  0.166  0.167  0.173  0.174  0.168  0.168
## IDNAM6   0.070  0.074  0.135  0.141 -0.007 -0.007
## IDNAM64 -0.011 -0.009 -0.019 -0.020  0.008  0.010
## IDNAM8   0.141  0.144  0.127  0.138  0.158  0.153
## IDNAM9  -0.030 -0.031 -0.131 -0.128  0.117  0.113
```

Here are the corresponding signs:

```r
sign(coefs_mtqM > 0)
```

```
##          [,1] [,2] [,3] [,4] [,5] [,6]
## IDNAM10     1    1    1    1    1    1
## IDNAM11     1    1    0    0    1    1
## IDNAM12     1    1    1    1    1    1
## IDNAM13     0    0    0    0    1    1
## IDNAM14     1    1    1    1    1    1
## IDNAM15     1    1    1    1    1    1
## IDNAM17     1    1    1    1    1    1
## IDNAM18     1    1    1    1    1    1
## IDNAM2      1    1    1    1    1    1
## IDNAM22     1    1    1    1    1    1
## IDNAM23     1    1    1    1    1    1
## IDNAM24     0    0    0    0    0    0
## IDNAM25     1    1    0    0    1    1
## IDNAM26     1    1    0    0    1    1
## IDNAM27     1    1    1    1    1    1
## IDNAM28     1    1    1    1    1    1
## IDNAM29     1    1    1    1    1    1
## IDNAM3      1    1    1    1    1    1
## IDNAM30     1    1    0    0    1    1
## IDNAM31     1    1    1    1    1    1
## IDNAM32     0    0    1    1    0    0
## IDNAM33     1    1    1    1    1    1
## IDNAM34     1    1    1    1    1    1
## IDNAM36     1    1    1    1    1    1
## IDNAM37     1    1    1    1    1    1
## IDNAM38     1    1    1    1    1    1
## IDNAM39     1    1    1    1    1    1
## IDNAM4      1    1    1    1    1    1
## IDNAM40     1    1    1    1    1    1
## IDNAM41     1    1    1    1    1    1
## IDNAM42     1    0    0    0    1    1
## IDNAM46     1    1    1    1    1    1
## IDNAM48     0    0    0    0    0    0
## IDNAM5      0    0    0    0    0    0
## IDNAM50     0    0    0    0    0    0
## IDNAM54     1    1    1    1    1    1
## IDNAM6      1    1    1    1    0    0
## IDNAM64     0    0    0    0    1    1
## IDNAM8      1    1    1    1    1    1
## IDNAM9      0    0    0    0    1    1
```

Below are signs that indicate which genotypes yield better model fit when their effect is differentiated from the RC reference level. A 1 indicates a positive effect, a -1 indicates a negative effect, and a 0 indicates that the genotype was not determined to provide better model when differentiated from the RC reference level.

```
inference_mtqM = sign(coefs_mtqM) * sign(AIC_IDs_mtqM < 0)
inference_mtqM
```

```
##          [,1] [,2] [,3] [,4] [,5] [,6]
## IDNAM10     1    1    0    0    1    1
## IDNAM11     0    0    0    0    0    0
## IDNAM12     1    1    0    0    1    0
## IDNAM13     0    0    0    0    0    0
## IDNAM14     0    0    0    0    0    0
## IDNAM15     0    1    0    0    1    1
## IDNAM17     1    1    1    1    1    1
## IDNAM18     1    1    1    1    1    1
## IDNAM2      0    0    0    0    0    0
## IDNAM22     1    1    1    1    1    1
## IDNAM23     0    0    0    0    0    0
## IDNAM24     0    0    0    0    0    0
## IDNAM25     0    0    0    0    0    0
## IDNAM26     0    0    0    0    0    0
## IDNAM27     1    1    0    0    1    1
## IDNAM28     1    1    0    0    1    1
## IDNAM29     1    1    0    0    1    1
## IDNAM3      0    0    0    0    0    0
## IDNAM30     0    0    0    0    0    0
## IDNAM31     1    1    0    0    1    1
## IDNAM32     0    0    0    0    0    0
## IDNAM33     1    1    0    0    0    0
## IDNAM34     0    0    0    0    0    0
## IDNAM36     0    0    0    0    1    1
## IDNAM37     1    1    1    1    0    0
## IDNAM38     1    1    1    1    1    1
## IDNAM39     1    1    0    0    1    1
## IDNAM4      0    0    0    0    0    0
## IDNAM40     0    0    0    0    0    0
## IDNAM41     1    1    0    0    1    1
## IDNAM42     0    0    0    0    0    0
## IDNAM46     1    1    0    0    1    1
## IDNAM48     0    0   -1   -1    0    0
## IDNAM5      0   -1    0    0    0   -1
## IDNAM50     0    0    0    0    0    0
## IDNAM54     1    1    1    1    1    1
## IDNAM6      0    0    1    1    0    0
## IDNAM64     0    0    0    0    0    0
## IDNAM8      1    1    1    1    1    1
## IDNAM9      0    0   -1   -1    0    0
```

##### Results

Here are the effect size estimates for genotypes that were to determined to differentiate from the RC reference level in all considered models and had the same estimated sign across all considered models.

```
coefs_mtqM[abs(rowSums(inference_mtqM)) == 6, ]
```

```
##          [,1]  [,2]  [,3]  [,4]  [,5]  [,6]
## IDNAM17 0.284 0.281 0.170 0.166 0.409 0.407
## IDNAM18 0.170 0.175 0.191 0.191 0.158 0.168
## IDNAM22 0.210 0.206 0.171 0.173 0.256 0.243
## IDNAM38 0.169 0.163 0.156 0.157 0.181 0.172
## IDNAM54 0.166 0.167 0.173 0.174 0.168 0.168
## IDNAM8  0.141 0.144 0.127 0.138 0.158 0.153
```

##### maxNPQ response

```
m1_mmaxNPQ = lmer(maxNPQ ~ ID + Date_num + I(Date_num^2) +
              (1|plot_number_year),
           data = dat5, REML = FALSE, control = lmerControl(optimizer ="Nelder_Mead"))
m1_mmaxNPQ_full = lmer(maxNPQ ~ ID + Date_num + I(Date_num^2) +
                      Ta + VPD + Fsd + Precip + Precip_cum +
                      Ta_7day + VPD_7day + Fsd_7day + Precip_7day +
                      (1|plot_number_year),
           data = dat5, REML = FALSE, control = lmerControl(optimizer ="Nelder_Mead"))
AIC(m1_mmaxNPQ)
```

```
## [1] 267.5389
```

```
AIC(m1_mmaxNPQ_full)
```

```
## [1] -395.5314
```

```
m2_mmaxNPQ = lmer(maxNPQ ~ ID + Date_num + I(Date_num^2) +
              (1|plot_number),
           data = dat5 %>% filter(year == 2021), REML = FALSE,
           control = lmerControl(optimizer ="Nelder_Mead"))
m2_mmaxNPQ_full = lmer(maxNPQ ~ ID + Date_num + I(Date_num^2) +
                      Ta + VPD + Fsd + Precip + Precip_cum +
                      Ta_7day + VPD_7day + Fsd_7day + Precip_7day +
                      (1|plot_number),
           data = dat5 %>% filter(year == 2021), REML = FALSE,
           control = lmerControl(optimizer ="Nelder_Mead"))
AIC(m2_mmaxNPQ)
```

```
## [1] 57.385
```

```
AIC(m2_mmaxNPQ_full)
```

```
## [1] -191.3429
```

```
m3_mmaxNPQ = lmer(maxNPQ ~ ID + Date_num + I(Date_num^2) +
              (1|plot_number),
           data = dat5 %>% filter(year == 2022), REML = FALSE,
           control = lmerControl(optimizer ="Nelder_Mead"))
m3_mmaxNPQ_full = lmer(maxNPQ ~ ID + Date_num + I(Date_num^2) +
                      Ta + VPD + Fsd + Precip + Precip_cum +
                      Ta_7day + VPD_7day + Fsd_7day + Precip_7day +
                      (1|plot_number),
           data = dat5 %>% filter(year == 2022), REML = FALSE,
           control = lmerControl(optimizer ="Nelder_Mead"))
AIC(m3_mmaxNPQ)
```

```
## [1] -134.5632
```

```
AIC(m3_mmaxNPQ_full)
```

```
## [1] -233.9881
```

**Diagnostics**

Diagnostic plots for the maxNPQ response are provided (small model followed by full model for models 1 through 3). Deviations from modeling assumptions are mild with most noticeable deviations occurring in the tails of the distributions for residuals.

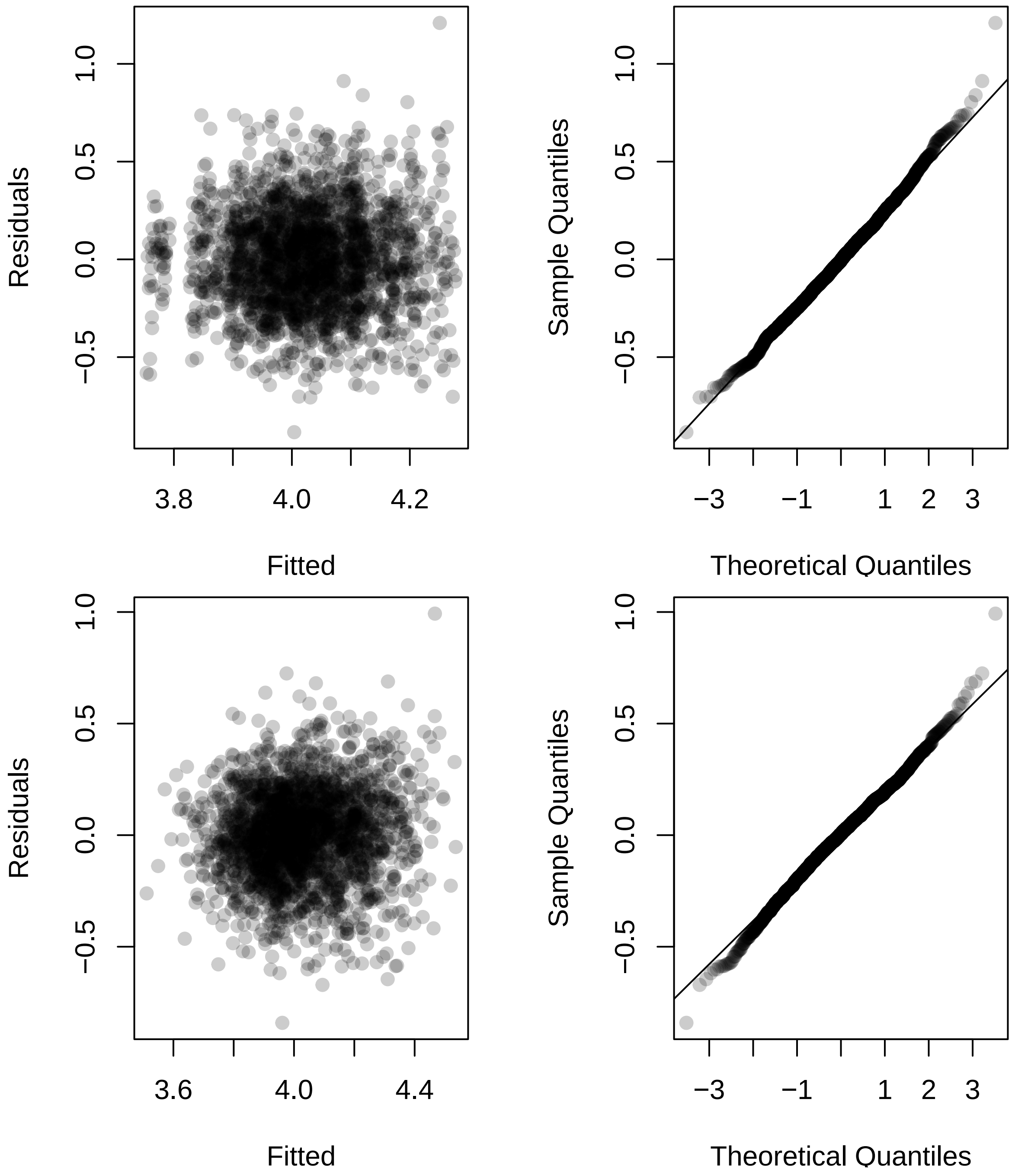

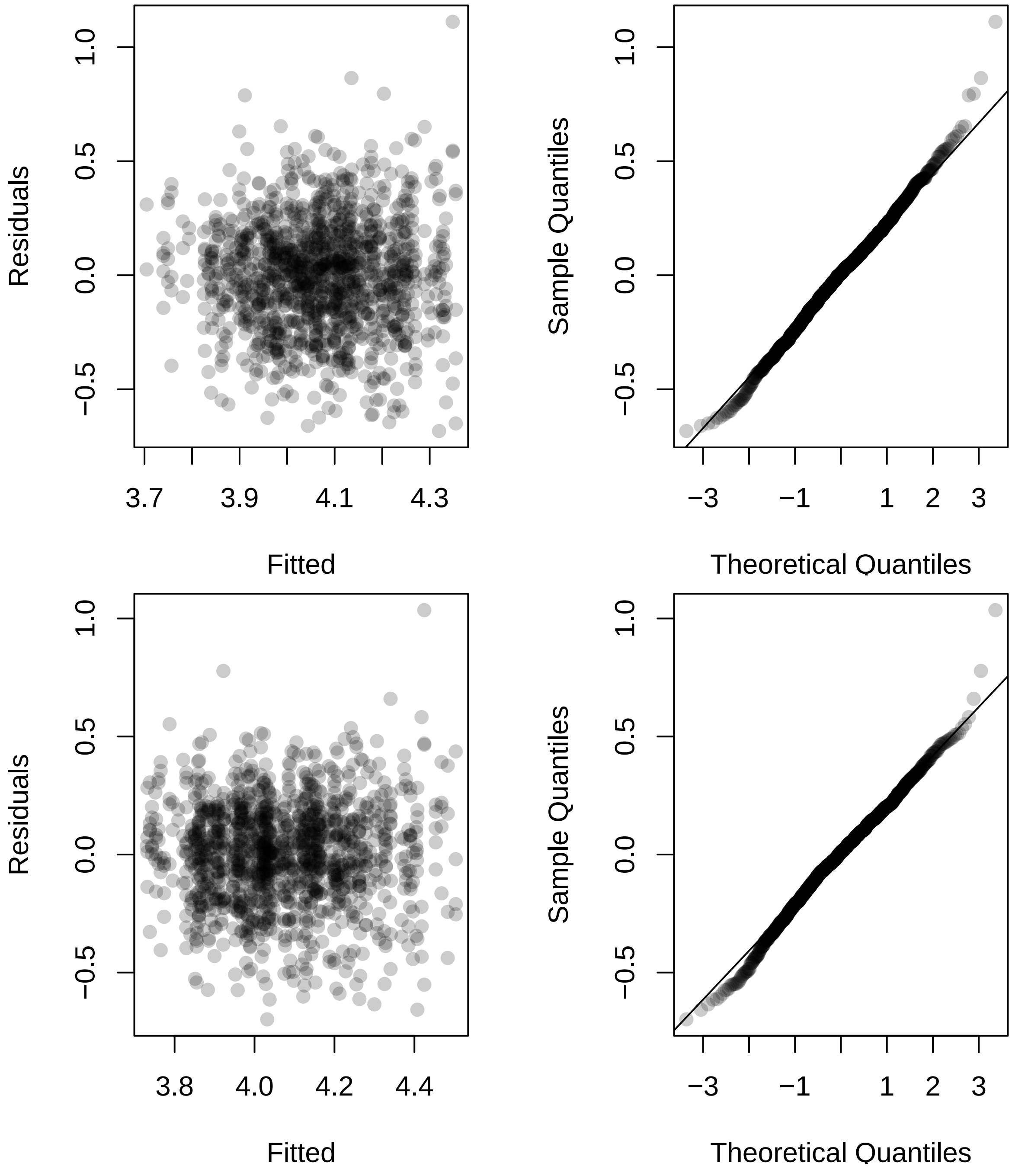
Residuals
Fitted
Sample Quantiles
Theoretical Quantiles
Residuals
Fitted
Sample Quantiles
Theoretical Quantiles

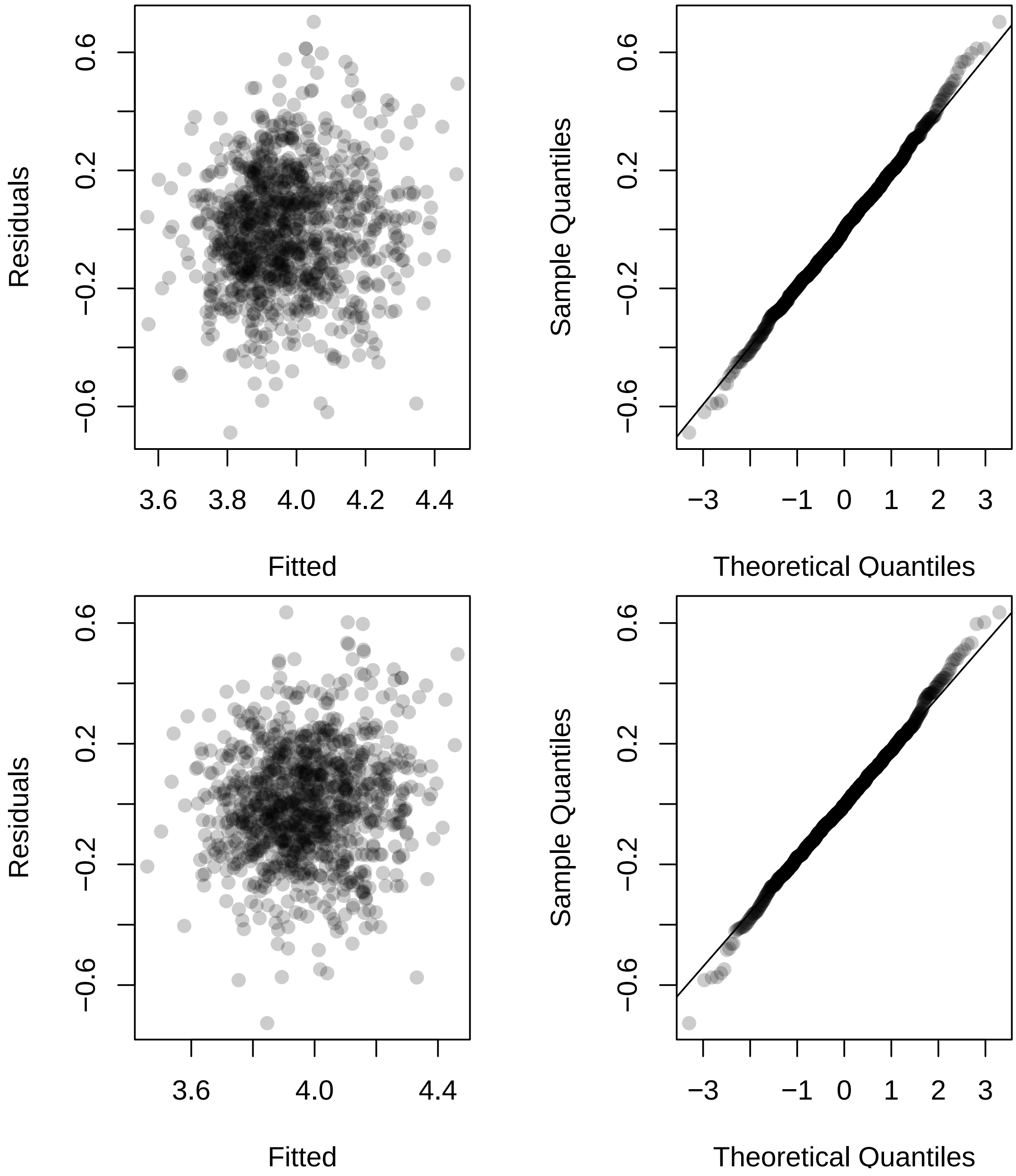

Residuals
Fitted
Sample Quantiles
Theoretical Quantiles

##### Investigate genotypes

We now perform an AIC based procedure to find genotypes that differ from the RC reference level.

```
## AIC for each ID variable from full maxNPQ fixed-effects model
M = model.matrix(maxNPQ ~ ID + Date_num + I(Date_num^2), data = dat5)
M_full = model.matrix(maxNPQ ~ ID + Date_num + I(Date_num^2) +
                      Ta + VPD + Fsd + Precip + Precip_cum +
                      Ta_7day + VPD_7day + Fsd_7day + Precip_7day,
                 data = dat5)

M2021 = model.matrix(maxNPQ ~ ID + Date_num + I(Date_num^2),
                 data = dat5 %>% filter(year == 2021))
M2021_full = model.matrix(maxNPQ ~ ID + Date_num + I(Date_num^2) +
                      Ta + VPD + Fsd + Precip + Precip_cum +
                      Ta_7day + VPD_7day + Fsd_7day + Precip_7day,
                 data = dat5 %>% filter(year == 2021))

M2022 = model.matrix(maxNPQ ~ ID + Date_num + I(Date_num^2),
                 data = dat5 %>% filter(year == 2022))
M2022_full = model.matrix(maxNPQ ~ ID + Date_num + I(Date_num^2) +
                            Ta + VPD + Fsd + Precip + Precip_cum +
                            Ta_7day + VPD_7day + Fsd_7day + Precip_7day,
                 data = dat5 %>% filter(year == 2022))

ncores = detectCores() - 2
AIC_IDs_mmaxNPQ = do.call(rbind, mclapply(
  grep("IDNA", colnames(M)), function(j){
    M1 = M[, -j]
    foo = lmer(maxNPQ ~ -1 + M1 + (1|plot_number_year),
               data = dat5, REML = FALSE)
    M1_full = M_full[, -j]
    foo_full = lmer(maxNPQ ~ -1 + M1_full + (1|plot_number_year),
               data = dat5, REML = FALSE)
    M12021 = M2021[, -j]
    bar = lmer(maxNPQ ~ -1 + M12021 + (1|plot_number),
               data = dat5 %>% filter(year == 2021), REML = FALSE)
    M12021_full = M2021_full[, -j]
    bar_full = lmer(maxNPQ ~ -1 + M12021_full + (1|plot_number),
               data = dat5 %>% filter(year == 2021), REML = FALSE)
    M12022 = M2022[, -j]
    baz = lmer(maxNPQ ~ -1 + M12022 + (1|plot_number),
               data = dat5 %>% filter(year == 2022), REML = FALSE)
    M12022_full = M2022_full[, -j]
    baz_full = lmer(maxNPQ ~ -1 + M12022_full + (1|plot_number),
               data = dat5 %>% filter(year == 2022), REML = FALSE)

    c(AIC(m1_mmaxNPQ) - AIC(foo),
      AIC(m1_mmaxNPQ_full) - AIC(foo_full),
      AIC(m2_mmaxNPQ) - AIC(bar),
      AIC(m2_mmaxNPQ_full) - AIC(bar_full),
      AIC(m3_mmaxNPQ) - AIC(baz),
      AIC(m3_mmaxNPQ_full) - AIC(baz_full))
  }, mc.cores = ncores))
```

```r
rownames(AIC_IDs_mmaxNPQ) = colnames(M)[grep("IDNA", colnames(M))]
```

Negative values indicate that the genotype in question provides better model fit when differentiated from the RC reference level.

```r
round(AIC_IDs_mmaxNPQ, 3)
```

```
##             [,1]    [,2]    [,3]    [,4]   [,5]    [,6]
## IDNAM10 -11.623 -13.107  -3.177  -4.004 -9.632 -10.065
## IDNAM11  -2.866  -2.575  -6.509  -9.131  1.957   1.986
## IDNAM12   1.618   1.612   1.890   1.755  0.262   0.121
## IDNAM13   1.248   1.374   0.632   0.304  1.999   2.000
## IDNAM14   0.128   0.637  -3.375  -4.244  1.577   1.605
## IDNAM15 -11.477 -11.978  -9.943 -13.339 -1.768  -1.692
## IDNAM17   1.991   1.997   1.685   1.674  1.862   1.740
## IDNAM18   2.000   1.977   1.975   1.966  1.915   1.817
## IDNAM2   -5.084  -4.309  -5.436  -7.097  1.072   1.097
## IDNAM22   0.648  -0.005   1.392   1.394  0.226   0.074
## IDNAM23   1.812   1.889   1.949   2.000  1.702   1.790
## IDNAM24 -13.004 -11.583 -17.982 -21.386  0.665   0.743
## IDNAM25  -0.303  -0.209   0.115   0.065  1.001   1.089
## IDNAM26   1.829   1.797   1.632   1.653  0.563   0.818
## IDNAM27   1.305   1.342  -1.570  -2.861  1.531   1.341
## IDNAM28  -2.193  -2.016  -0.078  -0.690 -0.021   0.092
## IDNAM29   1.883   1.916   0.770   0.785  1.763   1.659
## IDNAM3   -6.894  -6.158  -6.579  -7.669 -0.091   0.206
## IDNAM30  -0.973  -0.450  -7.863  -9.994  1.421   1.274
## IDNAM31 -23.274 -25.711 -21.944 -27.381 -6.679  -6.397
## IDNAM32   0.973   1.118  -0.876  -1.264  1.922   1.951
## IDNAM33  -5.701  -4.264  -1.196  -0.090 -3.430  -3.854
## IDNAM34   1.514   1.606   1.924   1.700  0.077  -0.016
## IDNAM36  -2.925  -2.896  -2.084  -2.977  0.284   0.292
## IDNAM37 -10.348 -10.580  -0.912  -1.349 -9.891 -10.185
## IDNAM38  -0.795  -0.606   0.991   0.917 -0.129  -0.142
## IDNAM39   1.455   1.390   0.969   0.610  1.978   1.991
## IDNAM4   -6.191  -8.866  -8.228 -11.700 -0.672  -0.515
## IDNAM40  -2.000  -1.516  -7.875 -10.380  1.876   1.842
## IDNAM41   0.832   0.554   1.187   0.866  1.170   1.265
## IDNAM42  -0.867  -0.910   0.701   0.262  0.356   0.586
## IDNAM46   1.413   1.528  -0.464  -1.043  1.711   1.618
## IDNAM48   1.666   0.936   1.406   0.911  1.663   1.611
## IDNAM5   -0.618  -0.192 -10.251 -12.080  0.669   0.351
## IDNAM50   1.901   1.760   1.974   2.000  1.525   1.467
## IDNAM54 -14.500 -13.322 -16.536 -18.639 -1.200  -0.928
## IDNAM6  -21.000 -19.948 -19.034 -24.026 -3.490  -3.386
## IDNAM64   1.713   1.887   1.999   1.978  1.707   1.757
## IDNAM8    1.780   1.776   0.552   0.242  1.753   1.785
## IDNAM9    1.610   1.487   1.935   1.943  0.947   1.021
```

```r
sign(AIC_IDs_mmaxNPQ < 0)
```

```
##         [,1] [,2] [,3] [,4] [,5] [,6]
## IDNAM10    1    1    1    1    1    1
## IDNAM11    1    1    1    1    0    0
## IDNAM12    0    0    0    0    0    0
```

```
## IDNAM13    0    0    0    0    0    0
## IDNAM14    0    0    1    1    0    0
## IDNAM15    1    1    1    1    1    1
## IDNAM17    0    0    0    0    0    0
## IDNAM18    0    0    0    0    0    0
## IDNAM2     1    1    1    1    0    0
## IDNAM22    0    1    0    0    0    0
## IDNAM23    0    0    0    0    0    0
## IDNAM24    1    1    1    1    0    0
## IDNAM25    1    1    0    0    0    0
## IDNAM26    0    0    0    0    0    0
## IDNAM27    0    0    1    1    0    0
## IDNAM28    1    1    1    1    1    0
## IDNAM29    0    0    0    0    0    0
## IDNAM3     1    1    1    1    1    0
## IDNAM30    1    1    1    1    0    0
## IDNAM31    1    1    1    1    1    1
## IDNAM32    0    0    1    1    0    0
## IDNAM33    1    1    1    1    1    1
## IDNAM34    0    0    0    0    0    1
## IDNAM36    1    1    1    1    0    0
## IDNAM37    1    1    1    1    1    1
## IDNAM38    1    1    0    0    1    1
## IDNAM39    0    0    0    0    0    0
## IDNAM4     1    1    1    1    1    1
## IDNAM40    1    1    1    1    0    0
## IDNAM41    0    0    0    0    0    0
## IDNAM42    1    1    0    0    0    0
## IDNAM46    0    0    1    1    0    0
## IDNAM48    0    0    0    0    0    0
## IDNAM5     1    1    1    1    0    0
## IDNAM50    0    0    0    0    0    0
## IDNAM54    1    1    1    1    1    1
## IDNAM6     1    1    1    1    1    1
## IDNAM64    0    0    0    0    0    0
## IDNAM8     0    0    0    0    0    0
## IDNAM9     0    0    0    0    0    0
```

We now investigate the specific coefficient effects from our fitted models. Specific interest is in determining the sign of the effect. Here are the effect estimates from each fitted model:

```
coefs_mmaxNPQ = round(cbind(summary(m1_mmaxNPQ)$coefficients[2:41, 1],
                    summary(m1_mmaxNPQ_full)$coefficients[2:41, 1],
                    summary(m2_mmaxNPQ)$coefficients[2:41, 1],
                    summary(m2_mmaxNPQ_full)$coefficients[2:41, 1],
                    summary(m3_mmaxNPQ)$coefficients[2:41, 1],
                    summary(m3_mmaxNPQ_full)$coefficients[2:41, 1]), 3)
coefs_mmaxNPQ
```

```
##            [,1]   [,2]   [,3]   [,4]   [,5]   [,6]
## IDNAM10  0.178  0.189  0.136  0.133  0.251  0.257
## IDNAM11  0.103  0.101  0.171  0.177  0.014  0.008
## IDNAM12 -0.029 -0.030  0.020  0.026 -0.094 -0.098
## IDNAM13  0.041  0.037  0.070  0.071 -0.002  0.000
## IDNAM14  0.064  0.055  0.136  0.132 -0.046 -0.045
## IDNAM15  0.170  0.176  0.204  0.210  0.133  0.134
## IDNAM17  0.004  0.002  0.034  0.031 -0.026 -0.036
## IDNAM18  0.001 -0.007  0.009  0.010 -0.021 -0.031
## IDNAM2   0.122  0.117  0.159  0.160  0.065  0.065
## IDNAM22 -0.055 -0.067 -0.047 -0.042 -0.094 -0.099
## IDNAM23 -0.020 -0.016 -0.013  0.000 -0.038 -0.032
## IDNAM24  0.180  0.173  0.262  0.260  0.080  0.078
## IDNAM25  0.071  0.070  0.081  0.074  0.071  0.068
## IDNAM26 -0.019 -0.021  0.036  0.032 -0.084 -0.076
## IDNAM27  0.040  0.039  0.110  0.117 -0.050 -0.060
## IDNAM28 -0.095 -0.095 -0.084 -0.086 -0.101 -0.099
## IDNAM29  0.016  0.014  0.065  0.058 -0.035 -0.042
## IDNAM3   0.142  0.136  0.177  0.170  0.101  0.094
## IDNAM30  0.080  0.074  0.182  0.182 -0.054 -0.061
## IDNAM31  0.241  0.255  0.289  0.295  0.212  0.210
## IDNAM32  0.047  0.044  0.098  0.095 -0.019 -0.016
## IDNAM33 -0.147 -0.132 -0.136 -0.100 -0.168 -0.176
## IDNAM34 -0.034 -0.031  0.016  0.029 -0.109 -0.112
## IDNAM36  0.103  0.104  0.117  0.117  0.092  0.093
## IDNAM37 -0.165 -0.169 -0.100 -0.098 -0.247 -0.252
## IDNAM38 -0.078 -0.076 -0.060 -0.056 -0.102 -0.104
## IDNAM39  0.034  0.037  0.059  0.062  0.010  0.007
## IDNAM4   0.137  0.159  0.189  0.198  0.119  0.116
## IDNAM40  0.094  0.089  0.185  0.188 -0.025 -0.028
## IDNAM41  0.051  0.057  0.053  0.056  0.065  0.062
## IDNAM42  0.079  0.081  0.067  0.070  0.090  0.084
## IDNAM46 -0.036 -0.032 -0.091 -0.092  0.038  0.044
## IDNAM48  0.027  0.049  0.045  0.056  0.041  0.044
## IDNAM5   0.076  0.070  0.205  0.199 -0.083 -0.093
## IDNAM50 -0.015 -0.023  0.010  0.000 -0.048 -0.051
## IDNAM54  0.191  0.186  0.254  0.245  0.125  0.120
## IDNAM6   0.226  0.224  0.269  0.275  0.165  0.165
## IDNAM64  0.025  0.016  0.002 -0.008  0.037  0.034
## IDNAM8   0.022  0.022  0.071  0.071 -0.035 -0.033
## IDNAM9   0.030  0.035  0.015  0.013  0.076  0.074
```

Here are the corresponding signs:

```r
sign(coefs_mmaxNPQ > 0)
```

```
##          [,1] [,2] [,3] [,4] [,5] [,6]
## IDNAM10     1    1    1    1    1    1
## IDNAM11     1    1    1    1    1    1
## IDNAM12     0    0    1    1    0    0
## IDNAM13     1    1    1    1    0    0
## IDNAM14     1    1    1    1    0    0
## IDNAM15     1    1    1    1    1    1
## IDNAM17     1    1    1    1    0    0
## IDNAM18     1    0    1    1    0    0
## IDNAM2      1    1    1    1    1    1
## IDNAM22     0    0    0    0    0    0
## IDNAM23     0    0    0    0    0    0
## IDNAM24     1    1    1    1    1    1
## IDNAM25     1    1    1    1    1    1
## IDNAM26     0    0    1    1    0    0
## IDNAM27     1    1    1    1    0    0
## IDNAM28     0    0    0    0    0    0
## IDNAM29     1    1    1    1    0    0
## IDNAM3      1    1    1    1    1    1
## IDNAM30     1    1    1    1    0    0
## IDNAM31     1    1    1    1    1    1
## IDNAM32     1    1    1    1    0    0
## IDNAM33     0    0    0    0    0    0
## IDNAM34     0    0    1    1    0    0
## IDNAM36     1    1    1    1    1    1
## IDNAM37     0    0    0    0    0    0
## IDNAM38     0    0    0    0    0    0
## IDNAM39     1    1    1    1    1    1
## IDNAM4      1    1    1    1    1    1
## IDNAM40     1    1    1    1    0    0
## IDNAM41     1    1    1    1    1    1
## IDNAM42     1    1    1    1    1    1
## IDNAM46     0    0    0    0    1    1
## IDNAM48     1    1    1    1    1    1
## IDNAM5      1    1    1    1    0    0
## IDNAM50     0    0    1    0    0    0
## IDNAM54     1    1    1    1    1    1
## IDNAM6      1    1    1    1    1    1
## IDNAM64     1    1    1    0    1    1
## IDNAM8      1    1    1    1    0    0
## IDNAM9      1    1    1    1    1    1
```

Below are signs that indicate which genotypes yield better model fit when their effect is differentiated from the RC reference level. A 1 indicates a positive effect, a -1 indicates a negative effect, and a 0 indicates that the genotype was not determined to provide better model when differentiated from the RC reference level.

```
inference_mmaxNPQ = sign(coefs_mmaxNPQ) * sign(AIC_IDs_mmaxNPQ < 0)
inference_mmaxNPQ
```

```
##          [,1] [,2] [,3] [,4] [,5] [,6]
## IDNAM10     1    1    1    1    1    1
## IDNAM11     1    1    1    1    0    0
## IDNAM12     0    0    0    0    0    0
## IDNAM13     0    0    0    0    0    0
## IDNAM14     0    0    1    1    0    0
## IDNAM15     1    1    1    1    1    1
## IDNAM17     0    0    0    0    0    0
## IDNAM18     0    0    0    0    0    0
## IDNAM2      1    1    1    1    0    0
## IDNAM22     0   -1    0    0    0    0
## IDNAM23     0    0    0    0    0    0
## IDNAM24     1    1    1    1    0    0
## IDNAM25     1    1    0    0    0    0
## IDNAM26     0    0    0    0    0    0
## IDNAM27     0    0    1    1    0    0
## IDNAM28    -1   -1   -1   -1   -1    0
## IDNAM29     0    0    0    0    0    0
## IDNAM3      1    1    1    1    1    0
## IDNAM30     1    1    1    1    0    0
## IDNAM31     1    1    1    1    1    1
## IDNAM32     0    0    1    1    0    0
## IDNAM33    -1   -1   -1   -1   -1   -1
## IDNAM34     0    0    0    0    0   -1
## IDNAM36     1    1    1    1    0    0
## IDNAM37    -1   -1   -1   -1   -1   -1
## IDNAM38    -1   -1    0    0   -1   -1
## IDNAM39     0    0    0    0    0    0
## IDNAM4      1    1    1    1    1    1
## IDNAM40     1    1    1    1    0    0
## IDNAM41     0    0    0    0    0    0
## IDNAM42     1    1    0    0    0    0
## IDNAM46     0    0   -1   -1    0    0
## IDNAM48     0    0    0    0    0    0
## IDNAM5      1    1    1    1    0    0
## IDNAM50     0    0    0    0    0    0
## IDNAM54     1    1    1    1    1    1
## IDNAM6      1    1    1    1    1    1
## IDNAM64     0    0    0    0    0    0
## IDNAM8      0    0    0    0    0    0
## IDNAM9      0    0    0    0    0    0
```

##### Results

Here are the effect size estimates for genotypes that were to determined to differentiate from the RC reference level in all considered models and had the same estimated sign across all considered models.

```
coefs_mmaxNPQ[abs(rowSums(inference_mmaxNPQ)) == 6, ]
```

```
##            [,1]   [,2]   [,3]   [,4]   [,5]   [,6]
## IDNAM10  0.178  0.189  0.136  0.133  0.251  0.257
## IDNAM15  0.170  0.176  0.204  0.210  0.133  0.134
## IDNAM31  0.241  0.255  0.289  0.295  0.212  0.210
## IDNAM33 -0.147 -0.132 -0.136 -0.100 -0.168 -0.176
## IDNAM37 -0.165 -0.169 -0.100 -0.098 -0.247 -0.252
## IDNAM4   0.137  0.159  0.189  0.198  0.119  0.116
## IDNAM54  0.191  0.186  0.254  0.245  0.125  0.120
## IDNAM6   0.226  0.224  0.269  0.275  0.165  0.165
```

#### Summary of influential genotypes across all models

Displayed below are all the genotypes that were determined to differ from the RC reference level in all six models and have an estimated effect with same estimated sign in all six models. The coding is as follows: a 1 indicates a positive effect in all six models, a -1 indicates a negative effect in all six models, and a 0 indicates a null result (either signs flipped across models or the genotype was not determined to differ from the RC reference level in one or models). A condensed table consisting of only influential genotypes can be seen on a new page following the table displayed below.

```
nam = paste("ID",
  (dat %>% pull(ID) %>% unique())[dat %>% pull(ID) %>% unique() %>% grepl("NAM", .)] %>% sort(),
  sep = "")
all6samesign = matrix(0, nrow = length(nam), ncol = 6)
rownames(all6samesign) = nam
colnames(all6samesign) = c("AqE", "AqI", "AqM", "tqE", "tqM", "maxNPQ")

all6samesign[rownames(all6samesign) %in%
               rownames(coefs_mAqE)[abs(rowSums(inference_mAqE)) == 6], 1] =
  mean(sign(coefs_mAqE[abs(rowSums(inference_mAqE)) == 6, ]))

all6samesign[rownames(all6samesign) %in%
               rownames(coefs_mAqI)[abs(rowSums(inference_mAqI)) == 6], 2] =
  sign(coefs_mAqI[abs(rowSums(inference_mAqI)) == 6, ]) %>% rowMeans()

all6samesign[rownames(all6samesign) %in%
               rownames(coefs_mtqE)[abs(rowSums(inference_mtqE)) == 6], 4] =
  sign(coefs_mtqE[abs(rowSums(inference_mtqE)) == 6, ]) %>% mean()

all6samesign[rownames(all6samesign) %in%
               rownames(coefs_mtqM)[abs(rowSums(inference_mtqM)) == 6], 5] =
  sign(coefs_mtqM[abs(rowSums(inference_mtqM)) == 6, ]) %>% rowMeans()

all6samesign[rownames(all6samesign) %in%
               rownames(coefs_mmaxNPQ)[abs(rowSums(inference_mmaxNPQ)) == 6], 6] =
  sign(coefs_mmaxNPQ[abs(rowSums(inference_mmaxNPQ)) == 6, ]) %>% rowMeans()

all6samesign
```

```
##          AqE AqI AqM tqE tqM maxNPQ
## IDNAM10   0   0   0   0   0      1
## IDNAM11   0   0   0   0   0      0
## IDNAM12   0   0   0   1   0      0
## IDNAM13   0   0   0   0   0      0
## IDNAM14   0   0   0   0   0      0
## IDNAM15   0   0   0   0   0      1
## IDNAM17   0   0   0   0   1      0
## IDNAM18   0   0   0   0   1      0
## IDNAM2    0   0   0   0   0      0
## IDNAM22   0   0   0   0   1      0
## IDNAM23   0   0   0   0   0      0
## IDNAM24   1   0   0   0   0      0
## IDNAM25   0   0   0   0   0      0
## IDNAM26   0   0   0   0   0      0
## IDNAM27   0   0   0   0   0      0
```

```
## IDNAM28   0   0   0   0   0       0
## IDNAM29   0   0   0   0   0       0
## IDNAM3    0   0   0   0   0       0
## IDNAM30   0   0   0   0   0       0
## IDNAM31   0   0   0   0   0       1
## IDNAM32   0   0   0   0   0       0
## IDNAM33   0  -1   0   0   0      -1
## IDNAM34   0   0   0   0   0       0
## IDNAM36   0   0   0   0   0       0
## IDNAM37   0  -1   0   0   0      -1
## IDNAM38   0   0   0   0   1       0
## IDNAM39   0   0   0   0   0       0
## IDNAM4    0   0   0   0   0       1
## IDNAM40   0   0   0   0   0       0
## IDNAM41   0   0   0   0   0       0
## IDNAM42   0   0   0   0   0       0
## IDNAM46   0  -1   0   0   0       0
## IDNAM48   0   0   0   0   0       0
## IDNAM5    0   0   0   0   0       0
## IDNAM50   0  -1   0   0   0       0
## IDNAM54   0   0   0   0   1       1
## IDNAM6    0   0   0   0   0       1
## IDNAM64   0   0   0   0   0       0
## IDNAM8    0   0   0   0   1       0
## IDNAM9    0   0   0   0   0       0
```

Here is a smaller table consisting of only genotypes that are influential.

```
all6samesign[rowSums(abs(all6samesign)) > 0, ]
```

```
##          AqE AqI AqM tqE tqM maxNPQ
## IDNAM10    0   0   0   0   0      1
## IDNAM12    0   0   0   1   0      0
## IDNAM15    0   0   0   0   0      1
## IDNAM17    0   0   0   0   1      0
## IDNAM18    0   0   0   0   1      0
## IDNAM22    0   0   0   0   1      0
## IDNAM24    1   0   0   0   0      0
## IDNAM31    0   0   0   0   0      1
## IDNAM33    0  -1   0   0   0     -1
## IDNAM37    0  -1   0   0   0     -1
## IDNAM38    0   0   0   0   1      0
## IDNAM4     0   0   0   0   0      1
## IDNAM46    0  -1   0   0   0      0
## IDNAM50    0  -1   0   0   0      0
## IDNAM54    0   0   0   0   1      1
## IDNAM6     0   0   0   0   0      1
## IDNAM8     0   0   0   0   1      0
```

Note that there are several genotypes that were very close to being selected as influential if it weren't for near-zero positive AIC differences or near-zero estimated coefficients in with the opposite sign of estimated coefficients in other candidate models.

### Single-day analysis for AqE response

```
dat5_single = dat5 %>% filter(Date %in% c("2021-07-20", "2022-07-19"))
#str(dat3_single)

dat5_single = dat5_single %>%
  mutate(Ta = as.numeric(scale(Ta)),
         VPD = as.numeric(scale(VPD)),
         Precip = as.numeric(scale(Precip)),
         Fsd = as.numeric(scale(Fsd)),
         Ta_7day = as.numeric(scale(Ta_7day)),
         VPD_7day = as.numeric(scale(VPD_7day)),
         Precip_7day = as.numeric(scale(Precip_7day)),
         Fsd_7day = as.numeric(scale(Fsd_7day)),
         Precip_cum = as.numeric(scale(Precip_cum)))
```

#### Modeling

We fit and compare models under three circumstances:

1. model fit to all data
2. model fit to 2021 data
3. model fit to 2022 data

We do not consider a reduced quadratic model in time since there is only one day under consideration.

```
m1_mAqE_single = lmer(mAqE ~ ID + Ta + VPD + Fsd + Precip + Precip_cum +
             Ta_7day + VPD_7day + Fsd_7day + Precip_7day +
               (1|plot_number),
           data = dat5_single)
AIC(m1_mAqE_single)
```

```
## [1] 158.2018
```

```
m2_mAqE_single = lm(mAqE ~ ID + Ta + VPD + Fsd + Precip + Precip_cum +
             Ta_7day + VPD_7day + Fsd_7day + Precip_7day +
               (1|plot_number),
           data = dat5_single %>% filter(year == 2021))
AIC(m2_mAqE_single)
```

```
## [1] 73.80967
```

```
m3_mAqE_single = lm(mAqE ~ ID + Ta + VPD + Fsd + Precip + Precip_cum +
             Ta_7day + VPD_7day + Fsd_7day + Precip_7day +
               (1|plot_number),
           data = dat5_single %>% filter(year == 2022))
AIC(m3_mAqE_single)
```

```
## [1] -15.61528
```

#### Diagnostics

Diagnostic plots for the AqE response are provided. Modeling assumptions appear to be satisfied.

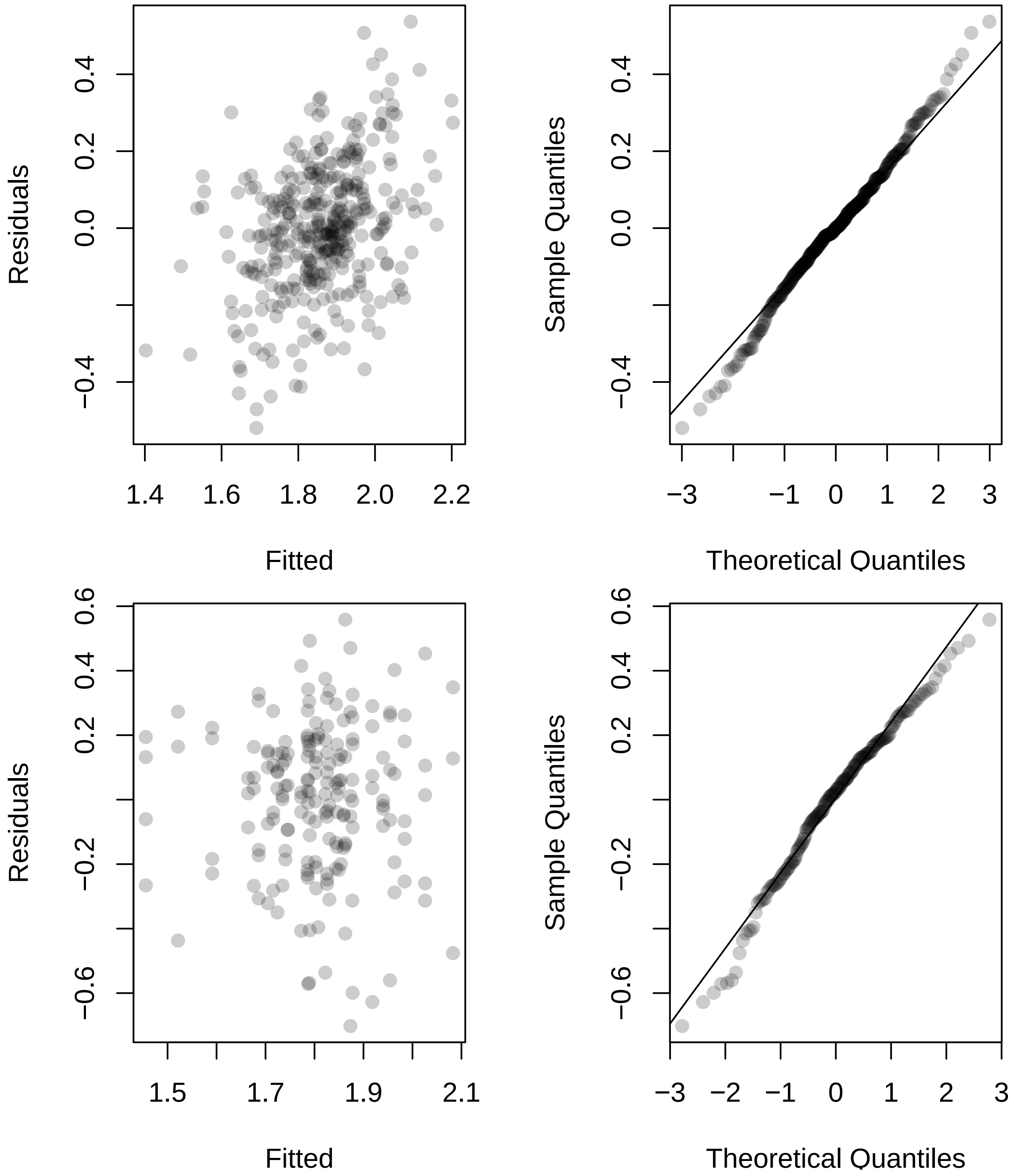
Residuals
Fitted
Sample Quantiles
Theoretical Quantiles
Residuals
Fitted
Sample Quantiles
Theoretical Quantiles

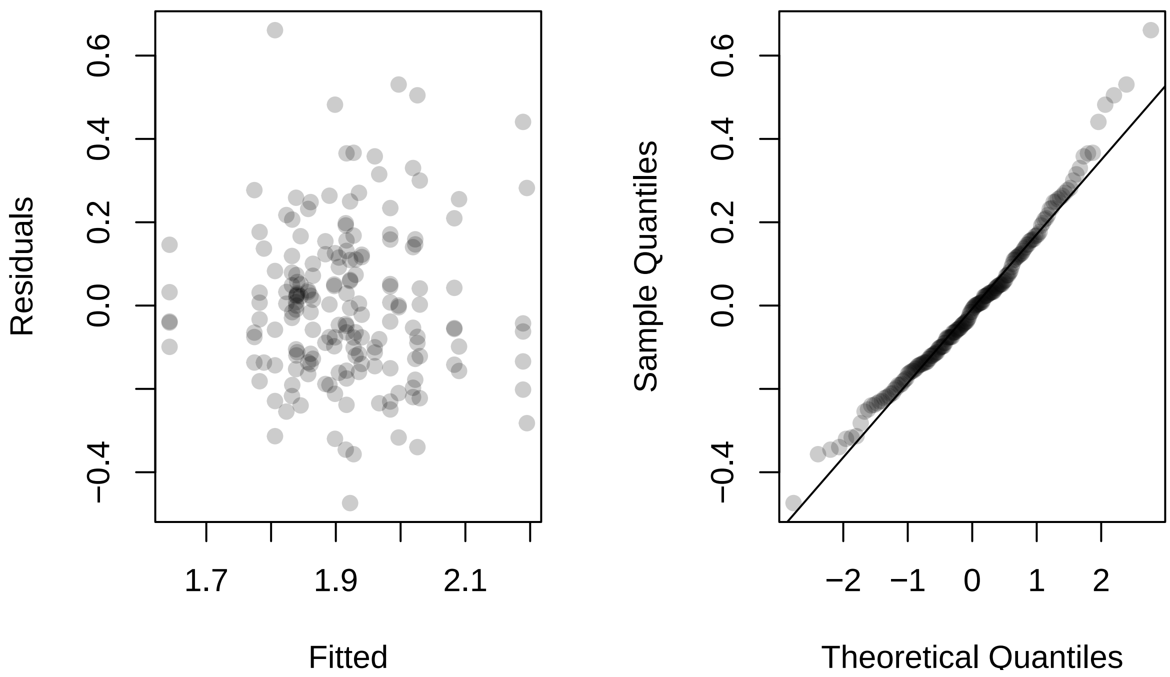
Residuals
Fitted
0.6
0.4
0.2
0.0
−0.4
1.7
1.9
2.1
Sample Quantiles
Theoretical Quantiles
−2
−1
0
1
2

#### Results

We now perform an AIC based procedure to find genotypes that differ from the RC reference level.

```
## AIC for each ID variable from full AqE fixed-effects model
M = model.matrix(mAqE ~ ID + Ta + VPD + Fsd + Precip + Precip_cum +
             Ta_7day + VPD_7day + Fsd_7day + Precip_7day,
                 data = dat5_single)
M2021 = model.matrix(mAqE ~ ID + Ta + VPD + Fsd + Precip + Precip_cum +
             Ta_7day + VPD_7day + Fsd_7day + Precip_7day,
                     data = dat5_single %>% filter(year == 2021))
M2022 = model.matrix(mAqE ~ ID + Ta + VPD + Fsd + Precip + Precip_cum +
             Ta_7day + VPD_7day + Fsd_7day + Precip_7day,
                     data = dat5_single %>% filter(year == 2022))

ncores = detectCores() - 2
AIC_IDs_mAqE_single = do.call(rbind, mclapply(
  grep("IDNA", colnames(M)), function(j){
    M1 = M[, -j]
    foo = lmer(mAqE ~ -1 + M1 + (1|plot_number),
                data = dat5_single)
    M12021 = M2021[, -j]
    bar = lm(mAqE ~ -1 + M12021 + (1|plot_number),
                data = dat5_single %>% filter(year == 2021))
    M12022 = M2022[, -j]
    baz = lm(mAqE ~ -1 + M12022 + (1|plot_number),
                data = dat5_single %>% filter(year == 2022))

    c(AIC(m1_mAqE_single) - AIC(foo),
      AIC(m2_mAqE_single) - AIC(bar),
      AIC(m3_mAqE_single) - AIC(baz))
  }, mc.cores = ncores))
rownames(AIC_IDs_mAqE_single) = colnames(M)[grep("IDNA", colnames(M))]
colnames(AIC_IDs_mAqE_single) = c("all", "2021", "2022")
```

Here are the effect size estimates for genotypes that were to determined to differentiate from the RC reference level in all considered models and had the same estimated sign across all considered models in the single-day analysis. There are no such genotypes.

```
coefs_mAqE_single = cbind(summary(m1_mAqE_single)$coefficients[2:41, 1],
                   summary(m2_mAqE_single)$coefficients[2:41, 1],
                   summary(m3_mAqE_single)$coefficients[2:41, 1])
inference_mAqE_single = sign(coefs_mAqE_single) * sign(AIC_IDs_mAqE_single < 0)

rownames(coefs_mAqE_single)[abs(rowSums(inference_mAqE_single)) == 3]
```

```
## character(0)
```

```
coefs_mAqE_single[abs(rowSums(inference_mAqE_single)) == 3, ]
```

```
##      [,1] [,2] [,3]
```

Notice that the analysis involving all of the data suggested that there was an influential genotype for the

AqE response.

```r
rownames(coefs_mAqE)[abs(rowSums(inference_mAqE)) == 6]
```

```
## [1] "IDNAM24"
```

```r
coefs_mAqE[abs(rowSums(inference_mAqE)) == 6, ]
```

```
## [1] 0.171 0.170 0.217 0.217 0.116 0.114
```
